# Cerebellar nuclei evolved by repeatedly duplicating a conserved cell type set

**DOI:** 10.1101/2020.06.25.170118

**Authors:** Justus M Kebschull, Noam Ringach, Ethan B Richman, Drew Friedmann, Sai Saroja Kolluru, Robert C Jones, William E Allen, Ying Wang, Huaijun Zhou, Seung Woo Cho, Howard Y Chang, Karl Deisseroth, Stephen R Quake, Liqun Luo

**Affiliations:** Department of Biology, Stanford University, Stanford, CA 94305, USA; Neurosciences Program, Stanford University, CA 94305, USA; Department of Bioengineering, Stanford University, Stanford, CA 94305, USA; Department of Applied Physics, Stanford University, Stanford, CA 94305, USA; Chan Zuckerberg Biohub, Stanford, CA 94305, USA; Society of Fellows, Harvard University, Cambridge, MA 02138, USA; Department of Animal Science, University of California Davis, CA 95616, USA; Center for Personal Dynamic Regulomes, Stanford University, Stanford, CA 94305, USA; Howard Hughes Medical Institute, Stanford University, CA 94305, USA; Department of Psychiatry and Behavioral Sciences, Stanford University, CA 94305, USA

## Abstract

How have complex brains evolved from simple circuits? Here we investigated brain region evolution at cell type resolution in the cerebellar nuclei (CN), the output structures of the cerebellum. Using single-nucleus RNA sequencing in mice, chickens, and humans, as well as STARmap spatial transcriptomic analysis and whole-CNS projection tracing in mice, we identified a conserved cell type set containing two classes of region-specific excitatory neurons and three classes of region-invariant inhibitory neurons. This set constitutes an archetypal CN that was repeatedly duplicated to form new regions. Interestingly, the excitatory cell class that preferentially funnels information to lateral frontal cortices in mice becomes predominant in the massively expanded human Lateral CN. Our data provide the first characterization of CN transcriptomic cell types in three species and suggest a model of brain region evolution by duplication and divergence of entire cell type sets.

## Introduction

The brains of extant animals are a product of hundreds of millions of years of evolution. Over time, cell types diversified (Arendt 2008; Arendt et al. 2016, 2019) and new brain regions appeared, giving rise to complex vertebrate brains today. Various models of brain region evolution have been proposed (Tosches 2017; Chakraborty and Jarvis 2015; Grillner and Robertson 2016; Frangeul et al. 2016). These include the duplication of entire regions followed by either divergence (neofunctionalization, supporting new functions) or maintenance (isofunctionalization, supporting more of the same function) of the duplicated products. Brain regions could also arise by splitting previously multifunctional regions into more specialized ones (subfunctionalization), or might evolve from *de novo* generation and combination of cell types. To our knowledge, however, none of these processes have been demonstrated in vertebrate brain evolution at cell type resolution. Doing so requires a comprehensive comparison of cell types across regions (Yao et al. 2020) and species (Hodge et al. 2019; Bakken et al. 2020; Boldog et al. 2018; Hodge et al. 2020; Tosches et al. 2018; Krienen et al. 2019; Peng et al. 2019; Hoang et al. 2019; Norimoto et al. 2020; Khrameeva et al. 2020) in a system that contains different numbers of homologous regions in different species.

The cerebellar nuclei (CN) are ideally suited for investigating brain region evolution. The cerebellum, consisting of the cerebellar cortex and CN, is an ancient hindbrain structure present in all jawed vertebrates (Montgomery, Bodznick, and Yopak 2012). It is classically involved in balance and fine motor control but also contributes to cognitive functions (Buckner 2013; Koziol et al. 2014; Wagner and Luo 2020). The cerebellum sends almost its entire output through the CN to a large number of target regions (Chan-Palay 1977; Teune et al. 2000) (Fig. 1A). Whereas the cerebellar cortex has expanded across evolution while maintaining a constant circuit motif (Yopak, Pakan, and Wylie 2016), the CN have been more plastic. Jawless vertebrates have cerebellum-like structures considered to be precursors to the cerebellar cortex but lack CN (Bell, Han, and Sawtell 2008). By contrast, a single pair of CN can be recognized in cartilaginous fishes and amphibians, two pairs in reptiles and birds, and three pairs in mammals (Yopak, Pakan, and Wylie 2016; Arends and Zeigler 1991). These findings suggest that the last common ancestor of jawed vertebrates had a single pair of CN, and CN numbers have increased in amniotes in the process of expanding the cerebellar output channels (Fig. 1B). Remarkably, the Lateral CN in humans expanded to be 17× larger than each of the other two nuclei (Tellmann et al. 2015), concomitant with the expansion of the prefrontal cortex that preferentially communicates with the lateral cerebellum (Bostan, Dum, and Strick 2013).

**Figure 1:**
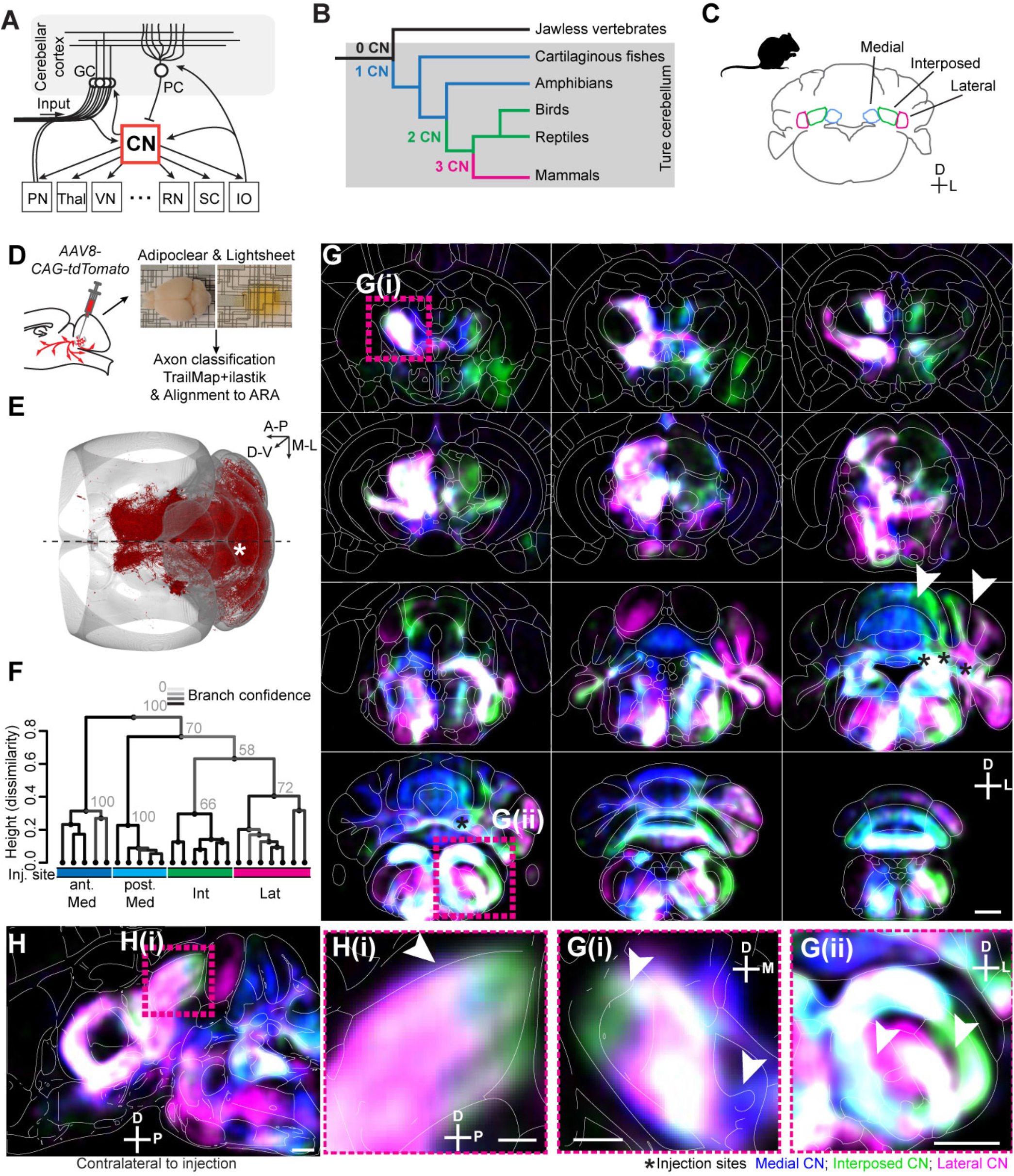
Brain-wide projections of mouse CN. (A) Schematic of the cerebellar circuit. Information enters the cerebellar cortex through mossy fibers from the PN and elsewhere in the brain, and climbing fibers from the IO. Purkinje cells send cerebellar cortex output to the CN, which project to many brain regions. PN, pontine nuclei; Thal, thalamus; VN, vestibular nuclei; RN, red nucleus; SC, superior colliculus; IO, inferior olive; GC, granule cells; PC, Purkinje cells. (B) Vertebrate cladogram, annotated with the number of CN pairs. (C) Schematic of the Medial, Interposed, and Lateral CN in mice. (D) Schematic of experimental workflow. Anterograde tracers were injected into individual nuclei. After expression, brains were cleared and imaged, and images were registered onto Allen CCF. Axons were quantified using a custom classification pipeline based on TrailMap (Friedmann et al. 2020) and ilastik (Berg et al. 2019). (E) Dorsal view of a representative brain volume with detected axons plotted in red. *, tracer injection site. Dashed line denotes the midline. (F) Dendrogram showing hierarchical clustering of 23 brains with injections into anterior Medial, posterior Medial, Interposed, and Lateral CN. Medial CN is most distinct from the other CN. Line color and grey numbers indicate bootstrapping-based branch confidence measured by the Approximately Unbiased p-value (out of 100 (Shimodaira 2005)). Values >40 indicate reasonable support. (G) Heat maps of axonal innervation from the three mouse CN. Coronal sections are shown, spaced 625 μm apart along the A–P axis, with Allen compartments in background. Scale bar = 1 mm. Heat maps were derived from N = 5 anterior Medial CN, 5 posterior Medial CN, 6 Interposed CN, and 7 Lateral CN injections. *, average tracer injection sites. Arrowheads and insets show shifted projections in (G(i)) contralateral thalamus and (G(ii)) ipsilateral brainstem. Scale bar = 500 μm. (H) Sagittal heat map, showing shifted projection patterns in the contralateral superior colliculus. Scale bar, main = 1 mm; inset = 500 μm. In this and all subsequent figures: A, anterior; P, posterior; D, dorsal; V, ventral; M, medial; L, lateral.

Despite their obvious importance in the cerebellar circuit, the CN are poorly understood. Their transcriptomic cell types have not been identified in any species, beyond a basic division into excitatory, GABAergic, and glycinergic neurons in rodents (Uusisaari and Knöpfel 2013). There have not been quantitative brain-wide comparisons of projection patterns of different cerebellar nuclei in any species (but see (Teune et al. 2000; Sugihara and Shinoda 2007; Chan-Palay 1977; Gould 1979; Aumann et al. 1994)), and few CN injections are available in the Allen Connectivity Atlas (Oh et al. 2014). Here, we characterize the transcriptomic cell types, spatial organization and CNS-wide projections of the three mouse CN, and compare these data to transcriptomic cell types in the two CN of chickens and the three CN of humans. We identify an archetypal CN—comprising a deeply conserved, stereotyped cell type set—as the unit of CN organization and evolution.

## Results

### CNS-wide projection mapping reveals shifting projection targets across mouse CN

Mouse CN are classically divided into three regions: Medial (Fastigial), Interposed, and Lateral (Dentate) CN (Fig. 1C; see Table S1 for nomenclature). The Medial CN is considered to be phylogenetically the oldest, and the Lateral CN the youngest (Yopak, Pakan, and Wylie 2016). These three nuclei differ in their axonal projection patterns (Chan-Palay 1977; Teune et al. 2000) and potentially gene expression (Chung, Marzban, and Hawkes 2009). To comprehensively characterize the differences between the individual nuclei, we began by comparing their projection patterns. We performed CNS-wide anterograde tracing of each nucleus using *AAV8-CAG-tdTomato* followed by brain and spinal cord clearing and light-sheet imaging (Chi et al. 2018; Ren et al. 2019; Friedmann et al. 2020) (Figs. 1D–H, S1–S9). We aligned all brain volumes to the Allen Common Coordinate Framework reference brain, detected axons using a custom classification pipeline (see Methods, Figs. 1D, S1) and quantified axonal innervation into 242 and 246 brain regions in the ipsi- and contralateral hemispheres, respectively (Figs. S7–S9; Table S2).

We traced 23 brains from 4 injection sites (anterior Medial, posterior Medial, Interposed, and Lateral CN). All three nuclei projected extensively to both hemispheres (Teune et al. 2000), innervating 125±34 and 140±32 (mean ± SD) brain regions in the ipsi- and contralateral hemispheres, respectively. Medial and Interposed CN also projected primarily to the contralateral cervical spinal cord (Fig. S6) (Asanuma, Thach, and Jones 1983). The brain-wide projection pattern of the Medial CN and in particular of the anterior Medial CN (which only has weak thalamic projections; Fig. S2) were most distinct, whereas projections of the putatively more recently diverged Interposed and Lateral CN were comparatively more similar (Figs. 1F, S7).

Closer inspection of CN projection patterns revealed cases where the three nuclei innervate adjacent brain regions with axons shifted relative to each other (Fig. 1G, H); such shifts were likely an underestimate of actual shift due to the spread of anterograde tracers at injection sites. Shifts were apparent in the ipsilateral cerebellar cortex (Fig. 1G), where Medial, Interposed and Lateral CN innervated the vermis, paravermis, and hemisphere, respectively (Gould 1979), and in the anterior contralateral thalamus (Fig. 1G(i)), where Interposed CN innervates regions shifted dorsolaterally relative to Lateral CN (matching observations in the rat (Aumann et al. 1994)) and Medial CN innervates regions shifted ventromedially (Gao et al. 2018). Other shifts were observed in the ipsilateral brainstem (Fig. 1G(ii)), where the three CN innervated adjacent parasagittal stripes, and in the contralateral superior colliculus (Fig. 1H), where Interposed CN innervated more posterior regions than the Lateral CN.

In summary, with the exception that the Lateral CN does not appear to innervate the spinal cord, all mouse CN innervate large portions of the CNS in both the ipsi- and contralateral hemisphere. Different nuclei innervate grossly similar regions in the thalamus, midbrain, and hindbrain. Their projections, however, are often shifted relative to each other such that different nuclei innervate adjacent volumes within or across brain region boundaries. Putatively more recently diverged Interposed and Lateral CN projections are more similar to each other than to Medial CN projections.

### Mouse CN comprise nucleus-specific excitatory neurons and nucleus-invariant inhibitory neurons

To investigate the molecular basis of the projection differences, we next used single-cell transcriptomics to determine the cell type composition of the CN. We separately dissected Medial, Interposed, and Lateral CN in each experiment and sorted NeuN+ neuronal nuclei into 384-well plates for high-depth, full-length single-nucleus RNA sequencing (snRNAseq; ~1 million aligned reads per cell) using a modified SmartSeq2 protocol (Schaum et al. 2018) (Fig. 2A). This strategy ensured relatively unbiased sampling of CN neuronal cell types and is directly transferable to frozen brain samples from other species due to the conservation of NeuN (Kim, Adelstein, and Kawamoto 2009; Bakken et al. 2018). After quality filtering and excluding contaminating cells, we retained high-quality data from 4605 mouse CN neurons.

**Figure 2:**
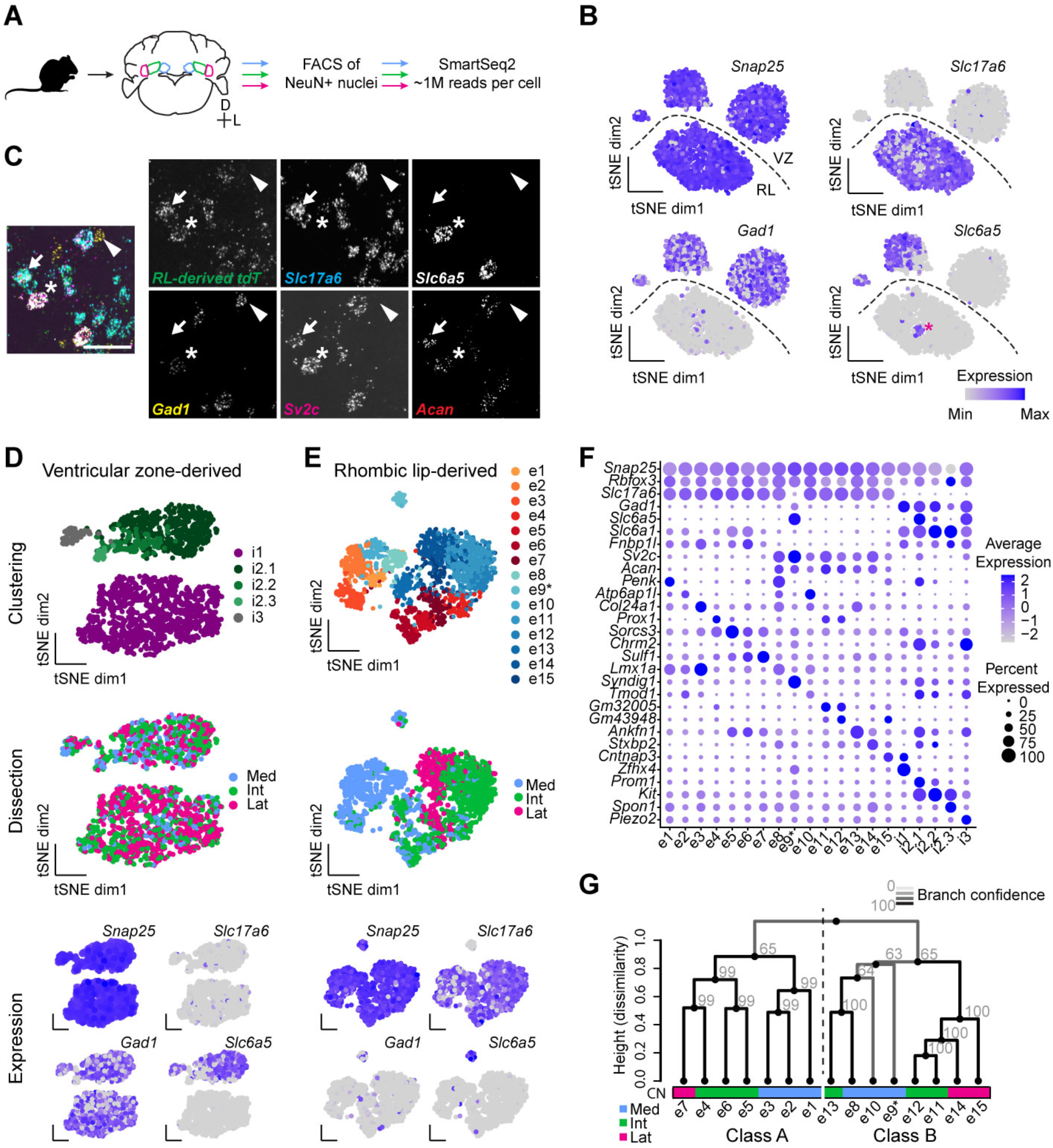
Mouse CN cell types. (A) Workflow of snRNAseq of mouse CN. The three cerebellar nuclei were dissected separately, cell nuclei were liberated, sorted for NeuN expression, and sequenced. (B) Marker gene expression for all CN neurons. The division into rhombic lip (RL)- and ventricular zone (VZ)-derived cells is indicated. N = 6 rounds of FACS using 9 mice each. (C) Representative image of permanently labeled RL-derived cells probed for endogenous marker gene expression. Arrow, excitatory neuron; arrowhead, inhibitory neuron; *, *Slc6a5*+ RL-derived. Scale bar = 50 μm. N = 2. (D, E) Clustering results for VZ- and RL-derived cells, labeled by clustering result (top) and CN dissection (middle), with marker gene expression at the bottom. (F) Marker gene expression for all mouse CN cell types. (G) Hierarchical clustering of excitatory CN cell types in the space of differentially expressed genes, using a correlation-based distance metric. Line color and grey numbers indicate bootstrapping-based branch confidence measured by the Approximately Unbiased p-value (out of 100 (Shimodaira 2005)). Values >40 indicate good support.

Overall, mouse CN neurons separated into 4 broad clusters. Three were *Gad1+* (encoding glutamic acid decarboxylase) inhibitory neurons. The remaining one was largely *Slc17a6*+ (encoding vesicular glutamate transporter 2, Vglut2) excitatory neurons; however, a small group of neurons within the *Slc17a6*+ cluster was *Slc17a6*– but *Slc6a5*+ (encoding glycine transporter 2, Glyt2) and likely glycinergic (Fig. 2B). We speculated that these broad divisions are driven by the developmental origins of excitatory and inhibitory CN neurons from the rhombic lip and ventricular zone, respectively (Elsen et al. 2013; Fink et al. 2006) (Fig. S10A). To test this, we permanently labeled rhombic lip-derived neurons with *tdTomato* using *Atoh1-Cre* (Matei et al. 2005) and performed STARmap *in situ* sequencing (Wang et al. 2018) on adult animals to quantify mRNA of various endogenous marker genes and *tdTomato* (Figs. 2C, S10). We found that all excitatory CN neurons were *tdTomato*+ and therefore derived from the rhombic lip. By contrast, all neurons falling into the three *Gad1*+ clusters were *tdTomato*– and likely ventricular zone-derived (Figs. 2C, S10). The exception were the *Slc6a5*+ neurons within the *Slc17a6*+ transcriptomic cluster (Fig. 2B, asterisk). These putative glycinergic neurons were *tdTomato*+ and therefore rhombic lip-derived (Figs. 2C, S10E). Based on their large size and location in the lateral part of the Medial CN (Fig. S10B), these cells likely correspond to the previously described large glycinergic projection neurons (Bagnall et al. 2009). For simplicity, we hereafter refer to rhombic lip- and ventricular zone-derived cells as “excitatory” and “inhibitory”, respectively.

To understand how neuronal cell types differ across CN, we separately clustered inhibitory and excitatory neurons (Figs. 2D, E). Inhibitory neurons showed relatively low diversity and formed 3 classes (Table S1). Class 1 and 3 each comprised a single transcriptomic cell type (i1, i3; referred to as cell type hereafter), while class 2 comprised one major (i2.1) and two minor (i2.2, i2.3) cell types. All cell types were represented across CN without discernible nucleus-specific changes (Figs. 2D, S11A–C). i1 neurons were *Gad1+Slc6a5–*, and likely corresponds to inferior olive-projecting CN inhibitory neurons (Prekop et al. 2018). i2.1 and i3 were *Slc6a5+* glycinergic neurons. In contrast to the relatively low diversity of inhibitory neurons, excitatory neurons formed 15 distinct cell types, each of which was specific to a single nucleus (Figs. 2E, S11D, E). Medial CN cell types were most distinct, whereas Interposed and Lateral CN cell types were more similar to each other (Fig. 2E), mirroring the projection data (Fig. 1F). While some of these cell types can be tentatively matched to previously described morphologically- or electrophysiologically-defined cell types (Fig. S12; see also (Fujita, Kodama, and Lac 2020)), the diversity uncovered from our study far exceeds previous reports.

In summary, mouse CN contain 5 nucleus-invariant inhibitory cell types in 3 classes and 15 nucleus-specific excitatory cell types, all of which can be distinguished by specific marker genes (Figs. 2F, S13).

### Excitatory neurons belong to two classes in each nucleus

If the three mouse CN arose from a single ancestral nucleus, cell types with a common evolutionary origin might exist in the different nuclei (Arendt 2008; Arendt et al. 2016). Such “sibling cell types” should share gene expression signatures that form an axis of variation independent of nucleus-specific changes. The nucleus-invariant inhibitory cell types found in each CN fulfill these requirements.

To investigate whether sibling cell types for excitatory neurons also exist, we hierarchically clustered all excitatory cell types in the space of differentially expressed genes between them (Fig. 2G). This analysis revealed a split of excitatory cell types into two classes, hereafter termed ‘Class A’ and ‘Class B’ (Table S1). On average, more genes were detected in Class B neurons than Class A neurons, hinting that Class B neurons might be larger than Class A neurons (Fig. S11F, G). Further, a large number of genes were differentially expressed in Class A and B neurons (Figs. S14, S15), including those with cell adhesion (Fig. S14B) and ion channel activity (Fig. S14C) that might contribute to different physiological properties of neurons in the two classes. Importantly, both Class A and Class B neurons were represented in each nucleus with 1–3 types each. We therefore consider the excitatory cell types within each class as putative sibling cell types to each other.

### Each CN subnucleus contains a stereotyped cell type set

The existence of both Class A and Class B sibling cell types in each nucleus indicates that the CN might have evolved through duplication. The finding of more than one Class A or Class B cell type within an individual nucleus, however, suggested that the CN are evolutionarily organized into units smaller than individual nuclei. Indeed, the mouse CN can each be divided into several subnuclei based on their cytoarchitecture (Paxinos and Franklin 2011; Sugihara and Shinoda 2007) (Table S1). To identify the relationship between these subnuclei and the CN cell types, we applied sequential STARmap *in situ* sequencing (Wang et al. 2018) to the CN. We detected up to 20 marker genes (Methods, Table S3) chosen to distinguish all cell types within each nucleus on coronal sections spanning the anterior–posterior axis of the CN (Figs. 3A–F, S16–19). We then classified neurons by cell type based on binarized marker gene expression (Fig. 3A, B) and inspected their location. Strikingly, we found that individual excitatory cell types were largely confined to cytoarchitecturally defined CN subnuclei: Medial CN is split into Med, MedL and MedDL, Interposed CN into IntA and IntP, and Lateral CN remains unsplit (Table S1). Within each subnucleus, Class A and Class B neurons were intermingled, albeit with local density differences (Figs. 3C–F, S16, S17). In addition, most subnuclei only contained a single excitatory cell type per class, and if two cell types from the same class were present in a subnucleus, they were often spatially segregated.

**Figure 3:**
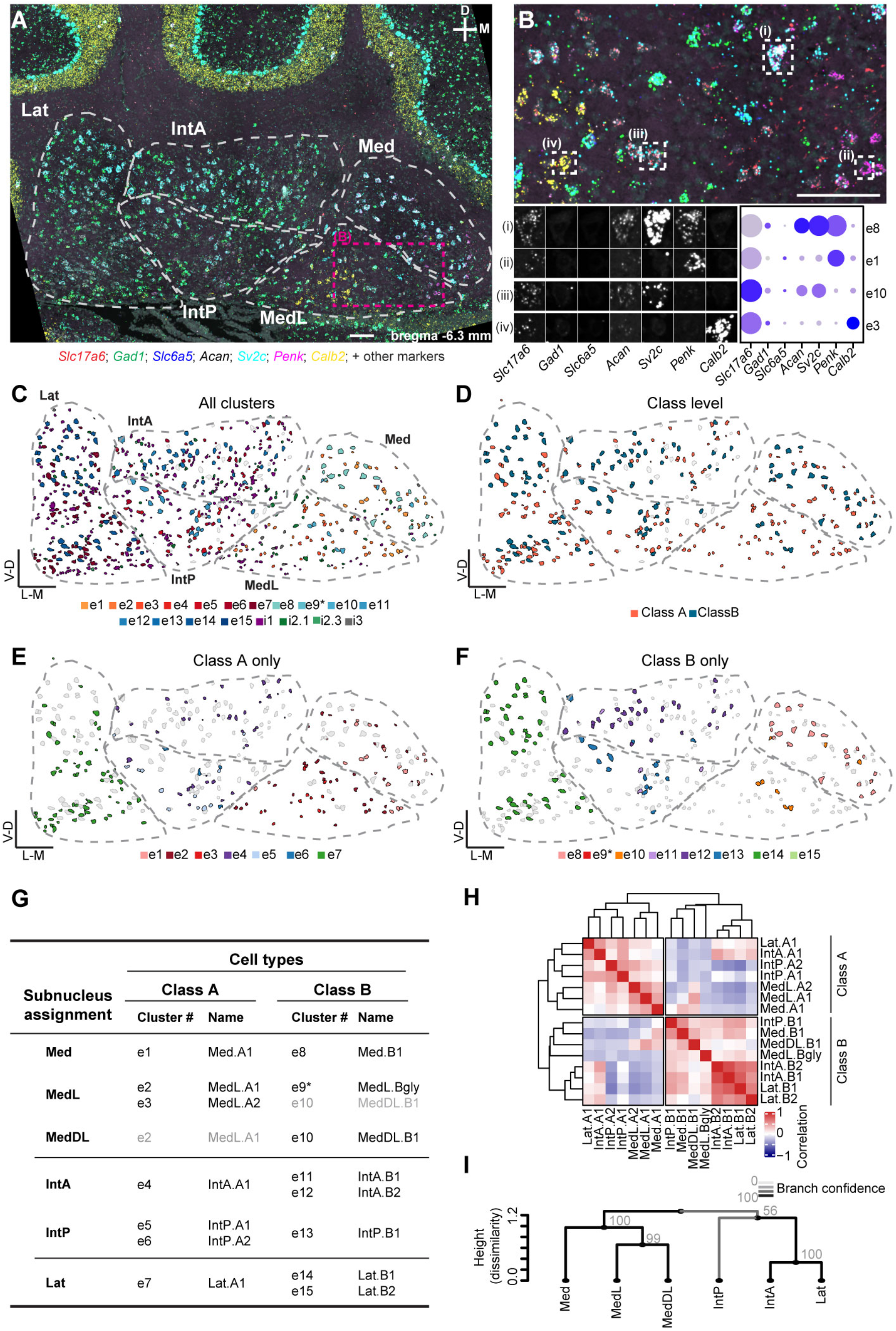
Spatial organization of CN mouse cell types. (A) Representative STARmap coronal section of the CN, showing 7 marker genes for illustration. Cytoarchitectonic subnuclei boundaries are indicated. Scale bar = 100 μm. N = 2, each dataset includes two hemispheres each of 3–6 coronal sections spanning the anterior–posterior axis of the CN. (B) Zoom in on the area marked in (A). Scale bar = 100 μm. Four excitatory cells are marked and decomposed into the 7 illustrated STARmap channels. Comparison to snRNAseq data (dot plot) yields the classification of the cells into transcriptomic cell types. (C–F) Classification results of the same section shown in (A). Colored cells represent all excitatory and inhibitory neurons by their assigned transcriptomic cell type according to the color scheme of Fig. 2 (C); excitatory neurons only colored by class (D); Class A-only (E) and Class B-only (F) excitatory neurons colored by their transcriptomic cluster showing subnuclei specificity. Unassigned neurons in each case are in gray. (G) Summary table of STARmap results for all excitatory cell types, noting the location of each cell type and new cell type names. Grey entries signify minor contributions to the indicated subnuclei. (H) Correlation matrix of all excitatory cell types annotated by subnuclei location. IntA correlates well with Lat in both Class A and Class B, whereas IntP is more similar to Medial CN cell types. (I) Hierarchical clustering of CN subnuclei. Line color and grey numbers indicate bootstrapping-based branch confidence measured by the Approximately Unbiased p-value (out of 100 (Shimodaira 2005)). Values >40 indicate good support.

As an example, consider the Interposed nucleus (Figs. 3C–F, S16, S17). Among Class B cell types, e13 was restricted to IntP, whereas e11 and e12 were both located in IntA only. However, e11 was located only in the most anterior part of IntA, and e12 was located in the more posterior part of IntA. Likewise, among Class A cell types, e5 and e6 were restricted to IntP—with e6 located more laterally than e5—and e4 was only located in IntA. To reflect these findings, we renamed excitatory cell types to indicate both their subnuclear location and class (Fig. 3G).

Both pairwise correlations between excitatory cell types (Fig. 3H) and hierarchical clustering of excitatory neurons grouped by subnuclei (Fig. 3I) revealed consistent relations between subnuclei within and across classes. Medial CN subnuclei grouped with each other. Whereas IntP grouped with the Medial CN in Class B, IntA was more closely related to Lat. Inspection of differentially expressed genes across subnuclei revealed both class-specific, subnucleus-independent (Fig. S14), and class-independent, subnucleus-specific gene sets (Fig. S19B).

In contrast to the subnuclear specificity of CN excitatory neurons, inhibitory neurons were broadly distributed across subnuclei (Fig. S18). The only exception was reduced numbers of i1 neurons and slightly increased numbers of i3 neurons in Medial CN, mirroring our snRNAseq data (Fig. S11E).

In summary, spatial transcriptomic analysis indicated a simple organizing principle for the CN. Subnuclei are the repeating units that form the CN and cerebellar output channels. Each subnucleus contains a stereotyped cell type set: 1–2 types each of Class A and Class B excitatory neurons that are subnucleus-specific, and 3 inhibitory classes that are subnucleus-invariant (Table S1).

### CN subnuclei as units of evolutionary duplication

Our mouse data suggest a model of CN evolution wherein a stereotyped cell type set is duplicated over the course of evolution to form a new CN subnucleus, accompanied by changes of gene expression and shifts of projection targets for the new subnucleus relative to old ones. To test this model, we investigated the transcriptomic cell types of chicken CN.

Chickens are thought to have two pairs of cerebellar nuclei without the equivalent of the mammalian Lateral CN (Fig. 1B) (Feirabend and Voogd 1986; Yopak, Pakan, and Wylie 2016). We dissected the entire chicken CN together for snRNAseq as done for the mouse (Fig. 4A). After quality filtering and removing contaminating cell types, we retained 1238 high-quality CN neurons. These cells split into major groups in a pattern comparable to mouse cells, with one broad group of excitatory neurons and two major groups of inhibitory neurons (Fig. 4B). Intriguingly, we could detect only very limited *SLC6A5* expression, indicating few if any glycinergic cells in the chicken CN.

**Figure 4:**
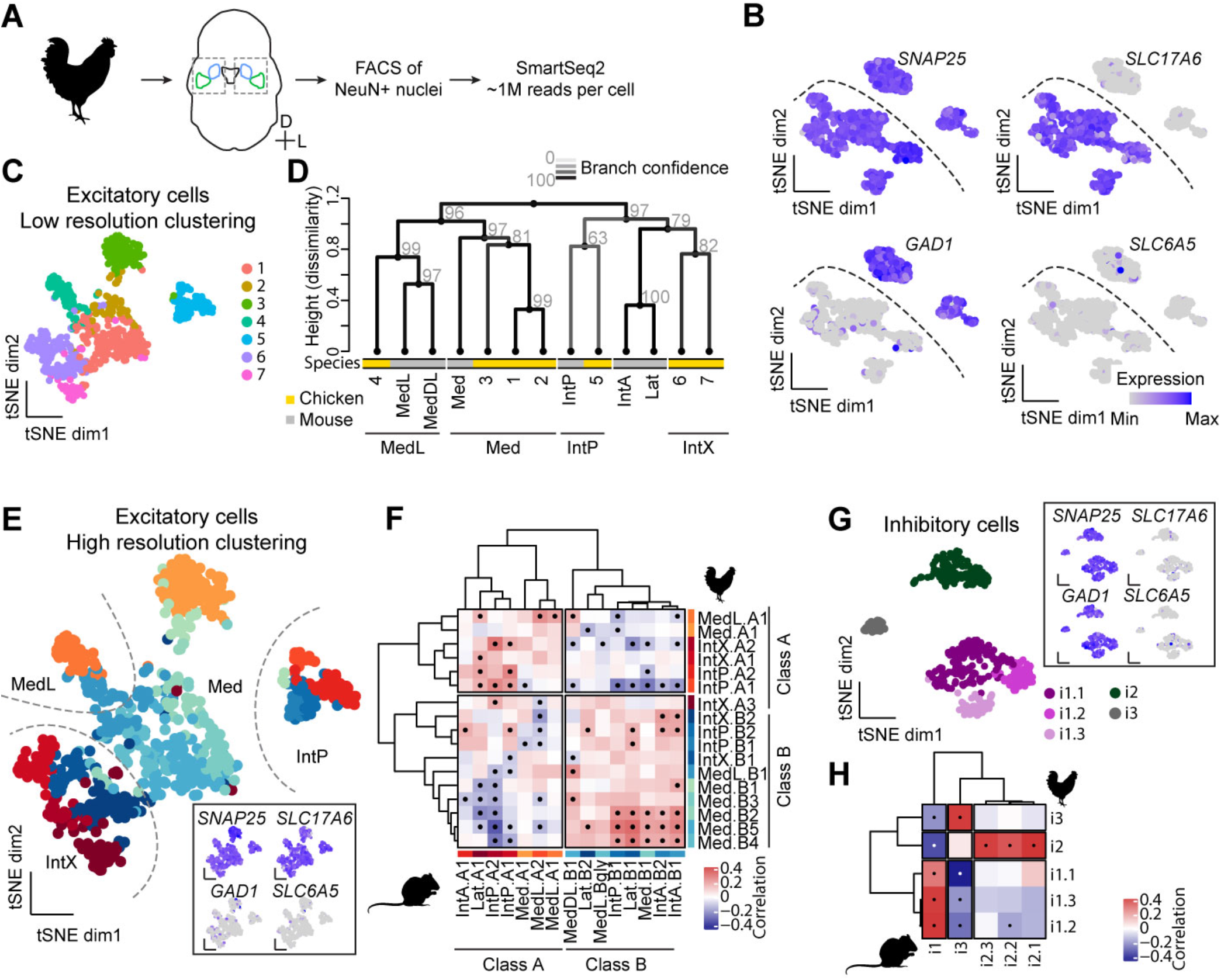
CN cell types in the chicken. (A) Workflow for chicken snRNAseq. The entire CN were dissected together from frozen tissue, cell nuclei were released and sorted for NeuN expression, and sequenced using a SmartSeq2 protocol. (B) Marker gene expression in all chicken CN neurons. The division into excitatory and inhibitory neurons is indicated. N = 3 chickens. (C) Coarse clustering result of all excitatory cells. (D) Dendrogram showing hierarchical clustering of coarse excitatory chicken clusters and mouse excitatory neurons grouped by subnuclei. Line color and grey numbers indicate bootstrapping-based branch confidence measured by the Approximately Unbiased p-value (out of 100 (Shimodaira 2005)). Values >40 indicate good support. (E) High-resolution clustering results of chicken excitatory neurons. Anatomical demarcations are inferred from comparative analysis with mouse subnuclei in panel D. Inset shows marker gene expression. (F) Correlation matrix between mouse and chicken excitatory cell types. A clear division of chicken excitatory cell types into Class A and Class B is apparent. Dots indicate significant correlations. (G) Clustering results of inhibitory neurons. Inset shows marker gene expression. (H) Correlation matrix between mouse and chicken inhibitory neurons. Dots indicate significant correlations.

To understand CN evolution at the level of subnuclei, we first focused on the excitatory chicken neurons and coarsely clustered them (Fig. 4C) based on the observation that mouse excitatory cells clustered coarsely by subnuclei (Figs. 2E, 3H). We then built a joint phylogenetic tree of these coarse chicken clusters and mouse subnuclei in the space of differentially expressed genes shared across species (Fig. 4D; Methods) (Tosches et al. 2018). Mouse subnuclei intermingled with chicken clusters, indicating that chicken CN contained regions homologous to mouse Med, MedL/MedDL, and IntP, but not IntA and Lat. The chicken CN also included an additional region that fell within the same clade as the mouse Interposed/Lateral CN. We term this region IntX. The identification of shared as well as new regions in the chicken and mouse support the notion that the CN number increased by the duplication and divergence of CN subnuclei.

### Excitatory and inhibitory neuron classes conserved across amniotes

Next, we sought to determine if the above model held at the resolution of cell types; specifically, is the distinction between Class A and Class B excitatory neurons in the mouse conserved in the chicken? We clustered the chicken excitatory neurons at a higher resolution, aiming to match clustering resolution between mouse and chicken data (Methods), and compared them to the mouse excitatory cell types (Fig. 4E, F). Correlational analysis between mouse and chicken cell types in the space of shared differentially expressed genes revealed both Class A and Class B excitatory cells in the chicken, with good correspondence to the mouse cell types (Fig. 4F). Importantly, each of the putative chicken CN regions identified above (Fig. 4D) contained representatives of both Class A and Class B neurons. We named the chicken cell types according to the mouse convention to reflect their inferred subnuclei (Fig. 4D) and class membership (Fig. 4F). Chicken Class B cells also had on average more genes detected than Class A cells (Fig. S20D, E), suggesting that Class B cells are larger than Class A cells in chickens as in mice. All chicken excitatory cell types could be robustly distinguished by differentially expressed genes (Fig. S20F, Fig. S21).

Analysis of chicken inhibitory neurons revealed 5 cell types which, like mouse inhibitory neurons, fell into three classes (i1–3, Figs. 4G, S20C). Correlation analysis to the mouse data showed a perfect match between the species at the class level (Fig. 4H). At a finer resolution, our data indicate independent cell type diversification or loss of ancestral diversity in chickens and mice in classes i1 and i2, respectively. Whereas the putatively inferior olive-projecting i1 class comprises three cell types in chickens, only a single cell type is found in mice. Conversely, while class i2 contains three cell types in mice, it contains only a single type in chickens.

Taken together, our chicken data indicate the conservation of the previously identified archetypal CN subnuclei in both excitatory and inhibitory cell classes. Our findings thus support the proposal that amniote CN evolved by repeatedly duplicating an archetypal CN subnucleus composed of a deeply conserved cell type set (Fig. 5I, left).

**Figure 5:**
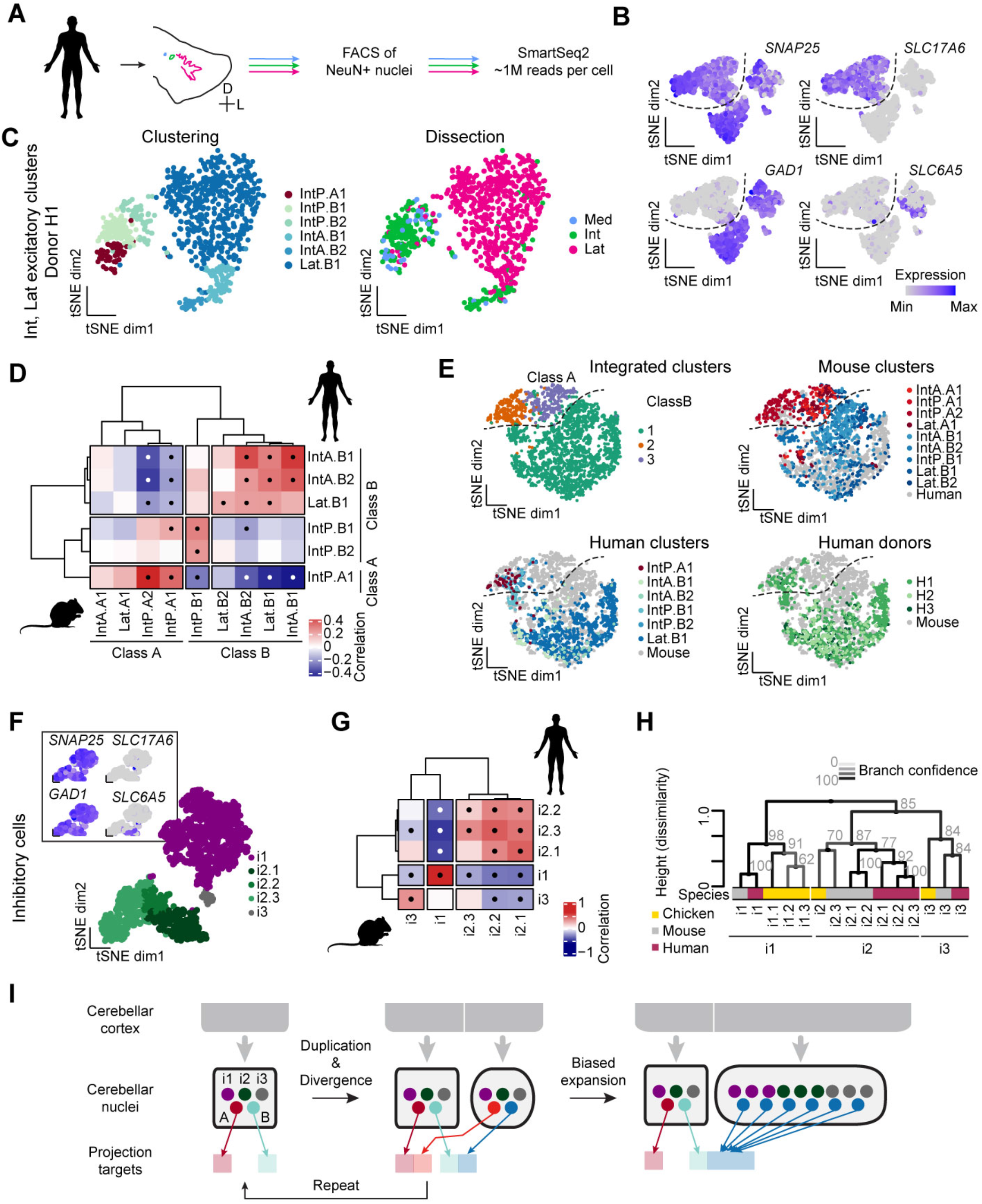
Class B neurons expanded in human Lateral CN. (A) Overview of workflow for human snRNAseq. The three CN are separately dissected from frozen tissue, and processed for snRNAseq of NeuN+ nuclei as done for mouse and chicken CN. (B) Marker gene expression for all human CN neurons. N = 3 donors. The division between excitatory and inhibitory neurons is indicated. (C) Interposed and Lateral CN excitatory clusters from donor H1, in which the entire Lateral CN was dissected and representative cells sequenced. Cells are colored by cluster assignment and dissection label. Some cells dissected with the Medial CN appear in Interposed clusters and likely represent Interposed CN contamination of the Medial CN dissection. (D) Correlation matrix of mouse and human donor H1 Interposed and Lateral CN excitatory cell types. Human Interposed CN contains cell types that correlate with both mouse Class A and Class B cell types. Human Lateral CN only correlates with Class B neurons. Dots indicate significant correlations. (E) Seurat integration of mouse and human Interposed and Lateral CN cells. Human Lateral CN cells fall exclusively into Class B domain of the tSNE plot, whereas Interposed cells fall into both classes. (F) Clustering results of human inhibitory neurons across all donors, integrated by FACS session using Seurat. Cells are colored by cluster assignment. Marker gene expression is indicated in the insert. (G) Correlation matrix of mouse and human inhibitory neurons, showing one-to-one correspondences. Dots indicate significant correlations. (H) Hierarchical clustering of CN inhibitory cell types across all three species (color coded). Conservation of three inhibitory classes across amniotes is apparent. Greyscales of line and numbers indicate bootstrapping-based branch confidence measured by the Approximately Unbiased p-value (out of 100 (Shimodaira 2005)). Values >40 indicate good support. (I) Schematic illustrating the proposed model of subnucleus duplication-and-divergence (left) and biased expansion of Class B excitatory neurons in human Lateral CN (right).

### Class B expanded at the expense of Class A in human Lateral CN

Cerebellar nuclei not only differ in number across vertebrates, but also the size of individual nuclei. The dramatic expansion of the human Lateral nucleus is a prime example. This expansion could be the result of an even increase in neuron numbers across all cell types, the formation of new subnuclei within the Lateral CN by duplication-and-divergence, or the formation of many *de novo* subnuclei within the Lateral CN. To test which model applies, we set out to determine the transcriptomic cell types of the human Medial, Interposed, and Lateral CN. We separately dissected the three nuclei from postmortem human cerebella and processed them for snRNAseq as done for mice and chickens (Fig. 5A). Importantly, in one donor (H1) we dissected and dissociated the entire Lateral CN to evenly sample its neuronal diversity to rule out biased cell recovery due to spatial heterogeneity. We obtained 3050 high-quality CN neurons, which clustered into 4 major groups, as in the mouse (Figs. 5B, S22, S23).

Human CN excitatory neurons readily separated by dissection labels (Fig. S22A), mirroring the nucleus-specificity observed in the mouse. Due to dissection difficulties of the small Medial CN, we collected only a small number of excitatory neurons from this nucleus. We therefore focused our analysis on Interposed and Lateral CN (Fig. 5C). We detected five distinct cell types in the Interposed CN. Surprisingly, Lateral CN neurons, although by far the largest population collected (206–535 cells per donor), formed only a single cluster. We then compared Interposed and Lateral CN excitatory cell types from mice and humans using correlation analysis (Tosches et al. 2018) (Fig. 5D) and Seurat data integration (Stuart et al. 2019) (Fig. 5E). Both analyses showed that while the human Interposed CN contained Class A and Class B neurons, all human Lateral CN excitatory neurons were of Class B, with few if any Class A neurons. Importantly, Lateral CN neurons from all donors gave the same result (Fig. 5E).

Clustering the inhibitory neurons revealed 5 CN-invariant cell types in 3 classes (Figs. 5F, S22B) with perfect correspondence to the mouse inhibitory classes (Fig. 5G). Intriguingly, the *Slc6a5*– i2.3 cell type, which is rare in mice, is much more abundant in humans, reducing the overall abundance of *Slc6a5*+ cells in human CN. Taken together with the absence of *Slc6a5*+ neurons in the chicken, this suggests that glycinergic neurons became abundant in the clade leading to the mouse after the divergence of rodents and primates. Hierarchical clustering of all inhibitory cell types in chickens, mice, and humans confirms the results of the pairwise comparisons (Figs. 4H, 5G), and supports the classification of CN inhibitory neurons into 3 conserved classes (Fig. 5H).

In summary, the human Interposed CN follows the cell type composition of the archetypal CN. However, in the human Lateral CN, Class B neurons are expanded at the expense of Class A neurons, suggesting that evolution tuned relative abundance of cell types within the framework of duplicating a stereotyped cell type set (Fig. 5I, right).

### Lateral nucleus Class A and Class B excitatory neurons preferentially connect via thalamus to medial and lateral frontal cortex, respectively

To investigate the relevance of the selective expansion of Class B neurons in human Lateral CN, we sought to determine how Class A and Class B neurons differ in their brain-wide projection patterns. As cell type-specific tracing is currently impossible in humans, we performed this analysis in mice, where both Class A and B neurons are abundant in the Lateral CN and high-resolution tracing is possible. Even in mice, however, no Cre lines faithfully distinguishing between the classes are available. We therefore used double retrograde tracing combined with STARmap *in situ* sequencing to identify projection targets of either class (Fig. 6A). We found that most target regions labeled both Class A and B neurons roughly equally (data not shown), which agrees with collateralization mapping experiments that indicate extremely broad projection patterns of CN neurons (Fig. S24). However, contralateral zona incerta (ZI) injections preferentially labeled Class A neurons in the Lateral CN, whereas contralateral brainstem reticular nucleus (Ret) injections primarily labeled Class B neurons (Fig. 6B).

**Figure 6:**
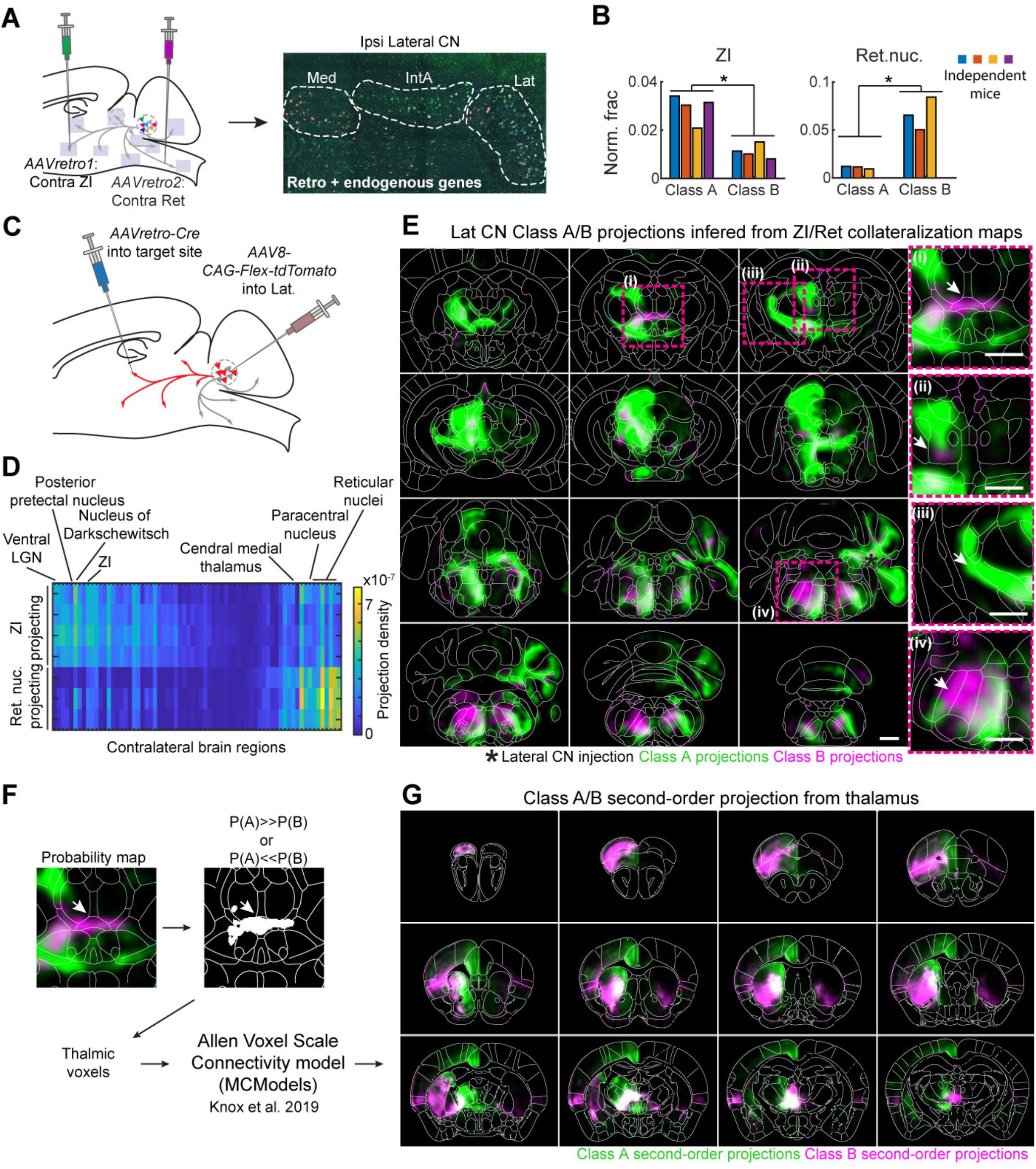
Differential projections of Lateral CN Class A and B neurons in mice. (A) Schematic of retrograde tracing and STARmap identification of Class A and B neurons in the Lateral CN. Contralateral zona incerta (ZI) and contralateral parvocellular reticular nucleus were injected with different *AAVretro* tracers. Gene expression was then measured by STARmap in the ipsilateral Lateral CN. (B) Quantification of retrograde tracing results across N = 4 independent mice for the Lateral CN at class resolution. *, p<0.05, paired t-test without corrections for multiple comparisons. (C) Schematic of collateralization mapping experiments. *AAVretro-Ef1a-Cre* was injected into either contralateral ZI or contralateral Ret, and *AAV-CAGs-FLEX-tdTomato* was injected into ipsilateral Lateral CN. After allowing marker expression, brains were cleared, volumetrically imaged, and aligned to Allen CCF. (D) Heat map showing all differentially innervated contralateral regions (p < 0.05, no multiple comparison correction) from Ret-projecting (N = 3) and ZI-projecting (N = 4) Lateral CN cells. Brain regions are sorted by mean innervation difference. (E) Probability maps of Class A and Class B projection patterns as computed from ZI and Ret collateralization patterns. Scale bar = 1 mm. Shown are coronal sections 625 μm apart along the A–P axis. Regions of differential intralaminar thalamus innervation ((i) and (ii)) and of the *AAVretro-Cre* injection sites ((iii) and (iv)) by Class A and Class B neurons are highlighted. Scale bar = 1 mm. (F) Workflow for *in silico* tracing of second-order projections from preferentially Class A- or B-innervated thalamic voxels. Starting voxels are identified and fed into a voxel scale connectivity model derived from Allen connectivity atlas injections (Knox et al. 2019). (G) Coronal sections (500 μm apart along the A–P axis) showing brain-wide normalized projection probabilities from thalamic voxels preferentially innervated by Class A (green) or Class B (magenta). Scale bar = 1 mm.

To investigate which other brain regions are differentially innervated by Class A and Class B neurons, we performed whole-brain collateralization mapping initiated at ZI and Ret (Schwarz et al. 2015). We injected *AAVretro-Ef1a-Cre* into either contralateral ZI or Ret and *AAV8-CAG-FLEX-tdTomato* into the ipsilateral Lateral CN (Fig. 6C), which would specifically label the brain-wide projections of Lateral CN neurons projecting to ZI and Ret, respectively. Ret injections labeled a smaller set of Lateral CN neurons, with a more restricted projection pattern than ZI-projecting neurons. In many brain regions, projections of the Ret-projecting neurons overlapped with those of ZI-projecting neurons (Figs. S25–S27, Table S4). However, several regions of the contralateral intralaminar nuclei of the thalamus—including paracentral nucleus and central medial nucleus—were more innervated by Ret-projecting neurons than ZI-projecting neurons (Figs. 6D, S25C, D).

As apparent from our retrograde tracing data (Fig. 6B), ZI- and Ret-projecting neurons do not perfectly correspond to Class A and B, respectively. Having obtained brain-wide projection probability maps for these two populations and knowing the ratio of Class A : Class B labeling from retrograde tracing, we could estimate the underlying projection probability maps for Class A and Class B neurons (Methods). The resulting computed maps reinforced the previous results of Class B projections to the intralaminar thalamus (Fig. 6E), but also highlighted intralaminar regions innervated by non-overlapping projections of both Class A and Class B neurons (Fig. 6E(ii)).

To investigate the relevance of these finer differences in Class A and B projections patterns, we first identified the thalamic voxels much more likely to be innervated by Class A than Class B Lateral CN neurons, and vice versa. We then used these voxels as starting points for *in silico* anterograde tracing using the recently published Allen Atlas voxel scale connectivity model (Knox et al. 2019), which is essentially a brain-wide connectivity matrix at 100 μm voxel resolution inferred from thousands of individual Allen Connectivity Atlas injections (Fig. 6F). The resulting projection probability maps revealed specific projections from primarily Class B innervated thalamic voxels to a lateral network of frontal association, ventral orbital, and insular cortices (Paxinos and Franklin 2011), as well as ventrolateral striatum (Fig. 6G, S28C, D). Conversely, *in silico* tracing from primarily Class A neuron innervated thalamic voxels revealed relatively broader projections to frontal cortical regions, but with a strong bias towards a medial network, including medial prefrontal cortex and anterior cingulate cortex, as well as dorsomedial striatum (Figs. 6G, S28A, B). We obtained similar results when we performed the same analysis based directly on ZI- and Ret-initiated collateralization maps rather than computed Class A/B projection maps (Fig. S28E– H), indicating that our results are not an artifact of our inferred Class level projection maps. Class A and B CN neurons, therefore, funnel information through the thalamus to different prefrontal networks in the mouse. Taken together with the expansion of Class B in the human Lateral CN, and assuming conservation of the discovered projection networks, these results suggest that cerebellar connectivity to the lateral prefrontal network is preferentially expanded in humans.

## Discussion

Here we present the first comprehensive dataset describing CN transcriptomic cell types and brain-wide projections in mice, as well as transcriptomic CN cell types in chickens and humans. These data reveal a conserved cell type set that makes up an archetypal CN unit, which we propose is effectively duplicated during evolution to increase the number of CN units—which in mice correspond to cytoarchitecturally defined subnuclei—and thus the number of cerebellar output channels. In addition, the predominance of Class B excitatory neurons in human Lateral CN indicates that the archetypal CN composition can be modified by varying the relative abundance of constituent cell types (Fig. 5I). By addressing how brain regions evolve at cell type resolution, our data extend previous studies of the evolution of brain regions and cell types.

### Subnuclei are the repeating units of the CN

At the outset of this study, we took advantage of the variations of the number of cerebellar nuclei in different species to investigate brain region evolution (Fig. 1B). We discovered instead that the fundamental repeating units in the mouse cerebellar nuclei are the subnuclei, each of which is formed by the same stereotyped cell type set (Fig. 3). This set contains 1–2 subnucleus-specific excitatory cell types each of Class A and Class B neurons, and the 3 classes of subnucleus-invariant inhibitory cell types.

Interestingly, comparisons of excitatory and inhibitory neurons across neocortical regions also suggest a region-specific set of excitatory cell types accompanied by a region-invariant set of inhibitory cell types (Yao et al. 2020; Tasic et al. 2018). Developmentally, neocortical excitatory neurons derive from the ventricular zone through local radial migration, whereas inhibitory neurons originate from the ventral forebrain through long-distance tangential migration (Marín and Rubenstein 2003). Thus, despite the opposite migratory paths giving rise to excitatory and inhibitory neurons, the CN and neocortex share a similar feature: region-specific excitatory cell types and region-invariant inhibitory cell types.

### Brain region evolution by duplication and divergence

Comparison between mice and chickens revealed that the stereotyped cell type set in subnuclei is deeply conserved across amniotes (Fig. 4), and thus likely describes an archetypic cell type composition of the CN in the last common ancestor of birds and mammals 320 million years ago. Our data suggest a model wherein CN subnuclei increased in number by repeatedly duplicating the entire cell type set—likely achieved by a coordinated expansion of cell numbers within all cell types followed by anatomical regionalization. Such duplication events were accompanied by divergence in gene expression in the excitatory but not inhibitory neurons (Figs. 2–5), and in projection patterns (Fig. 1). On the whole, CN evolution is therefore best described as region level duplication and divergence (Fig. 5I, left). At finer resolution, however, duplication-and-divergence or neofunctionalization is restricted to rhombic-lip derived excitatory neurons, and duplication-and-maintenance or isofunctionalization appears to govern the evolution of ventricular zone-derived inhibitory neurons.

We note that the developmental implementation of such regional “duplications” of a cell type set could take a multitude of paths. These include duplication of an early multipotent progenitor (Arendt et al. 2016) or establishment of a new region-defining morphogen gradient (O’Leary, Chou, and Sahara 2007; Green and Wingate 2014). Analysis of the CN in more species and detailed developmental investigations are needed to distinguish these possibilities. We expect that differences in the two developmental sources of CN neurons will explain the divergence vs. maintenance of transcriptomic state observed for excitatory and inhibitory neurons, respectively.

At a functional level, the duplication-and-divergence model of CN evolution implies that each CN subnucleus should be considered as a modular output node of the cerebellum. Taken together with evidence of topographic projections from Purkinje cells to the CN (Buisseret-Delmas and Angaut 1993; Sugihara and Shinoda 2007), and the extensively analyzed crystalline cerebellar motif (Ito 2006), this finding supports a model wherein specific regions of the cerebellar cortex and their connected CN subnuclei act as a functional module in parallel with other such modules (Herzfeld et al. 2017; Ekerot, Jörntell, and Garwicz 1995). Increased functionality of the cerebellum across evolution might then be implemented by the addition of such cerebellar cortex–CN modules to brain-wide circuits. The recently reported control of cerebellar cortex size by excitatory CN neurons (Willett et al. 2019; Fleming and Chiang 2015) and control of cerebellar cortex folding by mechanical constraints (Lawton et al. 2019) provide a simple mechanism for coordinated evolutionary expansion of CN and cerebellar cortex.

### Variations within the duplication-and-divergence framework

There is considerable variation in the brain region duplication-and-divergence framework proposed above. The existence of several representatives of Class A or B cell types in individual CN subnuclei suggests within-subnucleus cell type diversification (Fig. 3G). Conversely, varying numbers of cell types per inhibitory cell class i1 and i2 in mammals and chickens (Figs. 4, 5H) highlight the possibility of gain of new diversity or loss of ancestral diversity that uniformly affects all regions. Individuation of cell types after region-level duplication, moreover, can be dramatic, as illustrated by the apparent neurotransmitter switch in the rhombic-lip derived, *Slc17a6–*/*Slc6a5+* MedL.Bgly cell type of mouse Medial CN (Fig. 2C). Finally, the biased expansion of human Lateral CN illustrates the possibility of drastic changes in relative cell type abundance within the archetypal set. In the mouse, Class B neurons of the Lateral CN preferentially funnel information into frontal association cortex and lateral orbital and insular regions via the thalamus, whereas Class A neurons selectively access a medial network including medial prefrontal and anterior cingulate cortex (Fig. 6). The human lateral CN is greatly expanded relative to the other CN but it seems to have largely lost the Class A type neurons (Fig. 5). It is tempting to speculate that the expansion of human Lateral nucleus in general, and Class B type neurons within it, occurred in concert with the expansion of the human frontal cortical regions. We would therefore predict that the homolog to mouse lateral frontal cortex is expanded in the human. The precise evolutionary relationships between mouse frontal cortical regions and the human frontal cortex, however, are currently unclear (Carlén 2017; Laubach et al. 2018). Future comparative transcriptomic and connectomic work on mammalian frontal cortex evolution will shed more light on this important question.

In conclusion, our studies of the cerebellar nuclei evolution provide strong support for a duplication-and-divergence framework for brain region evolution at cell type resolution. Investigations of other brain regions using approaches similar to what is outlined here may provide insight into how generalizable this framework is, and will deepen our understanding of how brains changed over the course of evolution.

## Materials and Methods

Animal procedures were approved by the Stanford University or the University of California Davis Animal Care and Use Committee and were carried out in accordance with NIH standards. We used 8–12 week old male C57BL/6J mice (Jackson Labs, #000664) for all mouse experiments, except for STARmap sequencing of rhombic lip-derived cells (Figs. 2C, S10). For this experiment, we used 8–12 week old male *Atoh1-cre* (Jackson Labs, #011104) × Ai14 (Jackson Labs, #007914) animals. Chicken snRNAseq was performed on adult (~20-week-old) male chickens that were the F1 progeny of a Line 6 × Line 7 cross from the Avian Disease and Oncology Laboratory (ADOL). Human samples were obtained from Donor Network West and were deemed exempt from IRB regulations by Stanford University. Both donor H1 and donor H2 were 65-year-old white males, and donor H3 was a 39-year-old black female. All died of cancer, with no brain involvement.

### Sample processing and CN dissection

#### Mouse

To dissect mouse CN for snRNAseq, we cut acute coronal slices of the cerebellum according to previously described methods (Ren et al. 2011). Briefly, we deeply anesthetized the animals with intraperitoneal injection of avertin (300 mg/kg) and transcardially perfused them with 12 mL of ice-cold oxygenated perfusion solution (225 mM sucrose, 119 mM NaCl, 2.5 mM KCl, 1 mM NaH_2_PO_4_, 4.9 mM MgCl_2_, 0.1 mM CaCl_2_, 26.2 mM NaHCO_3_, 1.25 mM glucose, 3 mM kynurenic acid, 1 mM Na-ascorbate; all from Sigma). We then rapidly decapitated the mice and dissected the brains into ice-cold oxygenated slicing solution (110 mM choline chloride, 2.5 mM KCl, 0.5 mM CaCl_2_, 7 mM MgCl_2_, 1.3 mM NaH_2_PO_4_, 1.3 mM Na-ascorbate, 0.6 mM Na-pyruvate, 20 mM glucose, 25 mM NaHCO_3_ saturated with 95% O_2_ and 5% CO_2_; all from Sigma). We cut 300 μm thick coronal slices of the cerebellum on a vibratome (VT1000s, Leica), and collected them in ice-cold slicing solution or room temperature oxygenated artificial cerebrospinal fluid (125 mM NaCl, 2.5 mM KCl, 2 mM CaCl_2_, 1.3 mM MgCl_2_, 1.3 mM NaH_2_PO_4_, 1.3 mM Na-ascorbate, 0.6 mM Na pyruvate, 20 mM glucose, 25 mM NaHCO_3_, 50 μM APV, 20 μM DNQX, 100 nM TTX) (Hempel, Sugino, and Nelson 2007). We then rapidly dissected the individual nuclei from all sections containing the CN under a dissection microscope and flash froze dissected tissue in a dry ice/ethanol slurry, before storing it at –80 °C.

#### Chicken

We killed the adult male chickens by CO_2_ asphyxiation. We then rapidly dissected the cerebellum and flash froze the tissue in liquid nitrogen and stored it at –80 °C. For sectioning, we mounted each cerebellum in Optimal Cutting Temperature (OCT, Tissue Tek), and cut roughly coronal sections of 100 μm thickness on a cryostat (Leica). We then melted these sections onto clean microscope slides and rapidly froze the sections again on a metal plate resting on dry ice. We dissected the CN from the frozen sections using cold scalpel blades. To achieve dissection without shattering of the tissue, we rested the sections on a metal block cooled to approximated –20 °C by freezing 2.25 M CaCl_2_ (Bryan and Byrne 1970). Dissected tissue was stored at –80 °C before further processing.

#### Human

We obtained intact frozen human cerebella from tissue donors through Donor Network West on dry ice. After warming the tissue to –20 °C, we sectioned each cerebellum into 1–2 mm thick coronal sections using a manual frozen meat slicer (Garne-T), melted the sections onto large glass slides (Ted Pella), and rapidly froze them again on dry ice. We then separately dissected the individual nuclei using cold scalpel blades while resting the glass slide on a metal block cooled to –20 °C by freezing 2.25M CaCl_2_ (Bryan and Byrne 1970). Dissected samples were stored at –80 °C before further processing. For all donors, we dissected as much as possible of the Medial and Interposed CN. For Donor H1 we dissected the entire Lateral CN, separating dorsal and ventral Lateral CN of one hemisphere, and separately dissected small regions of dorsal and ventral Lateral CN from the second hemisphere to test for region-level differences in cell type composition. For Donors H2 and H3, we dissected the Lateral CN contained in one coronal section from the center of the CN.

### Tissue processing, FACS, and snRNAseq

CN samples from the three species were processed largely identically during the production of single nucleus suspensions, anti-NeuN staining, fluorescence-activated cell sorting (FACS), and subsequent sequencing library generation. We produced single nucleus suspensions and stained them for NeuN largely as previously described (Tasic et al. 2018). Briefly, we homogenized the dissected samples using an appropriately sized Dounce homogenizer on ice in >1 mL ice-cold homogenization solution [10 mM Tris pH 8.0 (Thermo Fisher), 250 mM sucrose (Sigma), 25 mM KCl (Sigma), 5 mM MgCl_2_ (Sigma), 0.1 % Triton-X 100 (Sigma), 0.5 % RNasin Plus RNase Inhibitor (Promega), 0.5 % SUPERase-In (Thermo Fisher), 1× Protease inhibitor (Promega), 0.1 mM DTT (Thermo Fisher)] per 100 mg of tissue with 15 strokes of the loose and 15–20 strokes of the tight pestle. The entire dissected sample was homogenized together, to avoid biased cell recovery from different regions of the sample. We then gravity filtered the suspension through either 30 μm (mouse) or 70 μm (chicken and human) filters and spun it down at 900×g, 4 °C for 10 minutes. Aliquots of the resulting pellet were flash-frozen in liquid nitrogen and store at –80 °C for future processing or used directly for staining. We resuspended pellets in staining solution [1× PBS (Gibco, pH 7.4), 0.8 % BSA, 0.5% RNasin Plus RNase Inhibitor (Promega), 0.5% SUPERase-In (Thermo Fisher)] and nutated them at 4 °C for 15 minutes. We then added mouse anti-NeuN primary antibody (Millipore MAB377) to each sample at 1:1000 (mouse) or 1:500 (chicken, human) and incubated the samples with agitation at 4 °C for 30 minutes. We pelleted the samples at 900×g, 4 °C for 10 minutes, resuspended them in staining solution containing 1:200 PE goat anti-mouse IgG secondary antibody (#405307, BioLegend) and 1:1000 Hoechst 33342 (Thermo Fisher) and incubated with agitation at 4 °C for 30 minutes. We then pelleted the sample again (900×g, 4 °C, 10 minutes) and resuspended in an appropriate volume of staining solution for FACS.

We used a Sony SH800S FACS machine to select for Hoechst+ NeuN+ neuronal nuclei and sorted them into 384-well lysis plates at the “ultrapurity” sort setting. Using index sorting information, we later refined the sorting gates to limit the selection to high backscatter events which selectively enriched for CN neuronal nuclei of all excitatory and inhibitory classes while removing the much smaller, but abundant, contaminating granule cell nuclei in all species. We either sorted positive events directly into wells or performed an enriching pre-sort at lower purity settings before resorting these events into wells. To avoid batch effects between different cerebellar nuclei of the same species, every mouse and chicken plate contained neuronal nuclei from every cerebellar nucleus. Most human H1 plates contained neuronal nuclei from every analyzed cerebellar nucleus, except for some plates that contained nuclei only from Interposed or Lateral CN. Donor H1 and H2 plates contained largely only Lateral CN neuronal nuclei.

Lysis plates and library preparation were performed as previously described (Schaum et al. 2018). Briefly, each well of the lysis plates contained 0.4 μL lysis buffer [0.5 U Recombinant RNase Inhibitor (Takara Bio, 2313B), 0.0625% Triton X-100 (Sigma, 93443-100ML), 3.125 mM dNTP mix (Thermo Fisher, R0193), 3.125 μM Oligo-dT30VN (Integrated DNA Technologies, 5′AAGCAGTGGTATCAACGCAGAGTACT30VN-3′), 1:600,000 ERCC RNA spike-in mix (Thermo Fisher, 4456740)]. After sorting, plates were spun down and frozen on dry ice before further processing. We performed cDNA synthesis using the Smart-seq2 protocol (Picelli, Faridani, et al. 2014). Plates were heated to 70 °C for 3 minutes and cooled to 10 °C to anneal the primers to the mRNA. We then added 0.6 μL RT-mix [16.7 U μL^−1^ SMARTScribe Reverse Transcriptase (Takara Bio, 639538), 1.67 U μl^−1^ Recombinant RNase Inhibitor (Takara Bio, 2313B), 1.67× First-Strand Buffer (Takara Bio, 639538), 1.67 μM TSO (Exiqon, 5′-AAGCAGTGGTATCAACGCAGAGTGAATrGrGrG-3′), 8.33 mM DTT (Thermo Fisher), 1.67 M Betaine (Sigma, B0300-5VL) and 10 mM MgCl_2_ (Sigma, M1028-10X1ML)] to each well using a Mantis liquid handler (Formulatrix). Reverse transcription was carried out at 42 °C for 90 minutes and stopped by heating to 70 °C for 5 minutes. We then added 1.5 μL PCR mix [1.67× KAPA HiFi HotStart ReadyMix (Kapa Biosystems, KK2602), 0.17 μM IS PCR primer (IDT, 5′-AAGCAGTGGTAT CAACGCAGAGT-3′), and 0.038 U μL^−1^ Lambda Exonuclease (NEB, M0262L)] to each well using the Mantis liquid handler and preamplified the cDNA using the following protocol; 1) 37 °C for 30 minutes, 2) 95 °C for 3 minutes, 3) N cycles of 98 °C for 20 seconds, 67 °C for 15 seconds and 72 °C for 4 minutes, and 4) 72 °C for 5 minutes. Mouse nuclei where subjected to N = 25 cycles, and human and chicken nuclei to N = 26 or N = 27 cycles.

We diluted the preamplified cDNA with 10 μL EB buffer (Qiagen) per well and quantified cDNA concentration using the Quant-iT PicoGreen dsDNA Assay (Thermo Fisher) on a fluorescent microplate reader according to the manufactures instructions in a 25 or 50 μL volume per well. We then diluted the amplified cDNA with EB buffer into a new 384-well plate at a concentration of 0.32 ng/μL and a volume of 0.4 μL using a Mantis fluid handler and a Mosquito HTS pipetting robot (TTP Labtech).

We produced Illumina sequencing libraries as previously described (Picelli, Björklund, et al. 2014). Briefly, we added 1.2 μL of tagmentation buffer (1.33× TAPS buffer (pH 8.5, 5 mM final MgCl_2_ concentration, Sigma), 10.67 % m/v PEG8000 (Promega), titrated amount of home-made Tn5 enzyme), and incubated at 55 °C for 5 minutes. We neutralized the reaction by adding 0.4 μL of 0.1 % SDS (Sigma) to each well. We then amplified and indexed the tagmented samples using Kapa HiFi (not hot start) polymerase (KK2102, Kapa Biosystems) and i5 and i7 indexing primers according to the manufacturer’s instructions using the following protocol; 1) 72 °C for 3 minutes, 2) 95 °C for 30 seconds, 3) 10 cycles of 95 °C for 10 seconds, 55 °C for 30 seconds, 72 °C for 1 minute, 4) 72 °C for 5 minutes. After PCR, we evenly pooled all wells of any given plate and cleaned up the library using SPRI beads (Beckman Coulter) using a dual purification of 0.8× beads followed by 0.8× or 0.7× beads. We quantified the libraries on a BioAnalyser (Agilent) and sequenced them using PE100 reads and 2 × 8 bp index reads on a Novaseq 6000 Sequencing System (Illumina) aiming for 1 million reads per cell.

### snRNAseq data processing and clustering

We aligned demultiplexed sequencing reads to both exons and introns of the relevant genome using STAR version 2.5.4 (Dobin et al. 2013) to maximize information per cell (Bakken et al. 2018). Specifically, for mice we used the Ensembl 92 annotation of mouse genome GRCm38 to produce a “pre-mRNA” annotation file, in which we reannotated ‘introns’ as ‘exons’ and used the STAR --quantMode geneCounts flag to count up reads per gene. For chickens, we used the Ensembl 99 annotation of chicken genome GRCg6a to produce a separate annotation file containing only introns following the procedures previously used in CRIES (https://github.com/csglab/CRIES). We aligned all reads using STAR to the chicken genome, using the exonic gtf file as guidance, and then counted reads in intronic and exonic regions using HTseq 0.10.0 using the --union and --intersection-strict flags with the intronic and exonic gtf files, respectively. For analysis, we combined the two read matrices. For human data we followed the same procedure as for chickens, using the Ensembl 94 annotation of GRCh38. The different alignment strategies gave comparable results with slightly increased numbers of genes detected in the separate alignment case. For snRNAseq data analysis, we largely followed standard procedures for filtering, variable gene selection, dimensionality reduction and clustering using Seurat v3 (v3.0.0 for mouse snRNAseq clustering; v3.1.5 for chicken and human, and all comparisons) (Stuart et al. 2019). Briefly, we read in the read × cell matrices obtained from HTseq into Seurat v3, filtered out cells with <500 genes (mice) or <2000 genes (human) detected, selected the 2000 most variable genes using the ‘vst’ method, calculated principal components and performed graph-based clustering. At this stage, we removed clusters of bad quality cells (low reads), non-neuronal clusters (low *Snap25*/*SNAP25* expression), and doublet clusters. We removed clusters of contaminating neuronal cell types by known marker expression (e.g. *Etv1*/*ETV1* expression in granule cells) and via Seurat v3 alignment to the recently published cell type atlas of mouse cerebellar cortex (Kozareva et al. 2020). We also removed very small, very distinct clusters (< 20 cells). In the chicken inhibitory cells, we removed two outlier clusters based on their lack of GRM1 expression, which is universal to all CN inhibitory clusters in the mouse, and all other chicken inhibitory clusters.

We then separately analyzed and clustered each of the major groups of CN neurons in each of the species. Specifically, in the mouse, we separately clustered excitatory cells, i1 cells, and glycinergic cells (i2.1, i2.2, i2.3, i3, e9* /MedL.Bgly). In the chicken, we separated inhibitory and excitatory cells. For low-resolution clustering (Fig. 4C) of the excitatory cells, we considered them all together. For higher resolution clustering (Fig. 4E), we separately clustered low-resolution clusters 3, 5, and the remaining clusters. Each round of clustering proceeded in the same way. Briefly, we selected the 2000 most variable genes, regressed out FACS round and the number of detected genes, and calculated a set of truncated PCs, which are composed of only the top 40 or 60 genes (Su et al. 2018). We selected the set of relevant PCs based on Elbow plots, JackStraw procedure, and manual inspection of PC loadings. We then intentionally over-clustered the cells using the Seurat Louvain clustering algorithm (resolution of 2 for mouse and human, resolution of 3 for chicken) and joined similar clusters using the Allen Institute’s scratch.hicatt package ‘merge_cl’ function using the following parameters; padj.th = 0.05, lfc.th = 1, low.th = 1, q1.th = 0.4, q.diff.th = 0.6, de.score.th = 40. This was necessary to allow for both small and large clusters in the same cell grouping, as pure Louvain clustering tended to merge distinct but much smaller clusters to neighboring larger clusters or split homogenous larger clusters into very similar small clusters. Differential gene expression was calculated using the default Wilcox rank-sum test.

Donor batch effects in the human dataset forced slight variations in this procedure. For human inhibitory cells, we used the Seurat data integration feature to integrate across FACS sessions (and the correlated donor structure), before proceeding with the pipeline described above. We note here that analysis of individual donors gave similar results as the integrated analysis. For excitatory cells, we largely analyzed each donor separately as described above, but integrated across them and mice using Seurat v3 for Fig. 5E.

### Hierarchical clustering of cell types and cross-species correlation analysis

For hierarchical clustering analysis of cell types and cross-species correlation analysis, we broadly followed procedures established in (Tosches et al. 2018). For hierarchical clustering within a species, we proceeded as follows. We first identified the set of differentially expressed genes between all cell types using a Wilcox Rank Sum test, requiring a minimum fold change of 2 and an adjusted p-value < 0.01. We calculated the average expression of these genes for every cell type and normalized expression of each gene by its average expression across cell types. We then hierarchically clustered the cell types using a Spearman correlation distance metric and average linkage criterion. Multilevel bootstrapping was performed as implemented in the R package *pvclust* with 10,000 steps (Suzuki R and Shimodaira H 2006). Bootstrapping confidence was returned as Approximately Unbiased p-value × 100, which gives a corrected estimate of how many trees out of 100 bootstrapped trees contained the same leaves below the respective node.

For cross-species analysis, we considered only one-to-one orthologous genes as defined by Ensembl. Within each species, we identified and normalized differentially expressed genes as above and then took the intersection of differentially expressed genes across species as a basis for further analysis. We either hierarchically clustered the cell types using a Spearman correlation distance metric as above (e.g. Figs 4D, 5H), or calculated Spearman correlation coefficients for all cross-species cell type pairs (e.g. Figs. 4F, 4H, 5D, 5G), and sorted the rows and columns of this matrix by hierarchical clustering using a Spearman correlation distance metric. We assessed the significance of cross-species correlations by shuffling the expression value of every gene between analyzed cell types 10,000 times, and computing the likelihood of obtaining a correlation coefficient as extreme or more extreme than the observed one. Correlations with p < 0.05 were labeled with a dot in the correlation matrix.

We note here that the clustering resolution/criterion in one species will to some extent influence the meaning of the resulting hierarchical clustering or correlation matrix by defining the gene space in which clustering or correlations are calculated. Consider the largely independent axes of variation of excitatory cell types by class and CN subnuclei. Mouse excitatory cell types hierarchically cluster first by class then by CN subnuclei (Fig. 2G). In contrast, chicken excitatory cell types cluster first by subnuclei, and then by class (Fig. S20G). Nevertheless, the correlation matrix between mouse and chicken excitatory cell types is dominated by the class level split, with CN subnuclei level correlations discernible at finer clustering resolutions. In this manuscript, we use this property to our advantage when we know more about one species than the other, like when comparing the coarse clustering of chicken excitatory neurons to mouse CN subnuclei (Fig. 4D). By clustering mouse excitatory cells by their STARmap defined CN subnuclei of origin, we shape the space of comparison towards genes that distinguish subnuclei (rather than by classes), helping us to define CN subnuclei in the chicken.

### STARmap *in situ* sequencing

We performed STARmap *in situ* sequencing largely as described in (Wang et al. 2018) using the thin section protocol, combined with the sequential gene readout presented in (Wang et al. 2018) for thick sections (i.e. every base in every sequencing round encodes one gene).

#### Gene sets

We manually selected various gene sets based on our snRNAseq data to distinguish CN cell types (Table S3). Taking into account cytoarchitectonic divisions between Medial, Interposed and Lateral CN, a minimum core set of 12 genes (*Slc17a6*, *Gad1*, *Slc6a5*, *Acan*, *Sv2c*, *Slc6a1*, *Ankfn1*, *Penk*, *Stac2*, *Calb2*, *Kitl*, *Sez6*; Fig. S9) was needed to distinguish all mouse CN cell types (except for i2.2). We designed gene-specific snail probes exactly as previously described (Wang et al. 2018). Probes were ordered as oPools oligo pools from IDT. For sequencing, we used seven 11 nucleotide orthogonal reading probes (OR1–7) and four 1-base fluorescent probes (1base_F1 to 1base_F4) labeled with Alexa 488, 594, 674 and 750, respectively (IDT). All sequences can be found in Table S3.

#### Sequential STARmap protocol

We performed STARmap library preparation as previously described (Wang et al. 2018) with minor modifications.

##### Tissue prep

Briefly, we deeply anesthetized a mouse with Isofluorane, then rapidly decapitated it, dissected out the brain, and froze it embedded in OCT (Tissue Tek) on dry ice. We then cut 16-μm sections on a cryostat and mounted them in 12- or 24-well glass-bottom dishes (P12G-1.5-14-F or P24G-1.5-13-F, Mattek) that were previously coated with Bind-Silane (GE17-1330-01, Sigma) followed by Poly-L-Lysine (P8920-100ML, Sigma) according to the manufacturer’s instructions. We fixed the sections in 4 % Paraformaldehyde (Electron Microscopy Sciences) for 10 minutes at room temperature, washed them 3 times in PBS, and permeabilized them in methanol precooled to –20 °C at 4 °C for 15 minutes and at – 80 °C for a minimum of 20 minutes, and for long term storage.

##### Library prep

For library preparation, we heated snail probes dissolved at 100 μM in RNase free H_2_O to 90 °C for 3 minutes, and let them cool to room temperature for approximately 10 minutes. We removed the samples from –80 °C freezer, and let them come to room temperature, before rehydrating them in PBSTR [1× PBS (Gibco), 0.1 % Tween-20 (Calbiochem), 0.1 U/μL SUPERase-In (Thermo Fisher)] for 2 minutes. Samples were then incubated in hybridization mix [10 nM of each oligo, 2×SSC (Sigma), 10 % formamide (Calbiochem), 1 % Tween-20, 20 mM RVC (NEB), 0.1 mg/mL salmon sperm DNA (Thermo Fisher)] with agitation at 40 °C overnight. We then washed the samples in PBSTV (1× PBS, 0.1 % Tween-20, 2 mM RVC) at room temperature for 2 × 20 minutes, and in 4× SSC in PBSTR at 37 °C for 20 minutes. We rinsed the sample once in PBST (1x PBS, 0.1 % Tween-20) and incubated it in T4 DNA ligation mixture [0.1 U/μL T4 DNA ligase (EL0011, Thermo Fisher), 1× ligase buffer, 0.1 mg/mL BSA, 0.2 U/μL SUPERase-In] with agitation at room temperature for 2 hours. After washing the samples in PBSTR at room temperature for 2 × 20 minutes, we incubated the samples in RCA mixture [2 U/μL Phi29 (EP0094, Thermo Fisher), 1× Phi29 buffer, 250 μM dNTP, 0.1 mg/mL BSA, 0.2 U/μL of SUPERase-In, 20 μM 5-(3-aminoallyl)-dUTP] at 30 °C for 4 hours. We then washed the samples in PBST at room temperature for 2 × 20 minutes and treated them with 20 mM acrylic acid NHS ester (Sigma; from fresh 0.5 M stock in DMSO) in PBST at room temperature for 2 hours. Samples were then stored overnight in PBSTR at 4 °C. We washed the sample in PBST briefly and incubated it in monomer buffer [4 % acrylamide (BioRad), 0.2 % BIS-acrylamide (BioRad), 2× SSC] at room temperature for 30 minutes. We removed the monomer buffer and added 12 μL polymerization solution (0.2 % ammonium persulfate, 0.2 % tetramethylethylenediamine in monomer buffer) to the center of the section, and rapidly covered it with a GelSlick (Lonza) coated 12 mm coverslip, taking care to remove any bubbles under the coverslip. We allowed the sample to polymerize for 1 hour at room temperature and then washed the sample in PBST 2 × 5 minutes, removing the GelSlick coverslip with forceps without disturbing the sample. We then digested the sample in digestion mix [0.8 mg/mL ProteinaseK (Thermo), 2× SSC, 1 % SDS] with agitation at 37 °C for 1 hour to overnight and finally washed the sample in PBST for 3 × 5 minutes.

##### Sequencing

We incubated the sample in sequencing mixture (0.1 U/μL T4 DNA ligase, 1× ligase buffer, 0.1 mg/mL BSA, 5 μM orthogonal reading probe, 0.25 μM 1base_F1 through F4 each) with agitation at room temperature for 3 hours to overnight, followed by washing in Washing&Imaging buffer (2× SSC, 10 % formamide) for 3 × 10 minutes. During the second wash, we added 1:1000 DAPI (Thermo). We imaged the samples using a 20× air objective on an Andor Dragonfly 500 spinning disk confocal, recording DAPI, 488, 594, 647, and 750 channels in z-stacks with 3 μm spacing. Samples were tiled and automatically stitched together with a 10 % overlap between tiles. After imaging, we stripped the samples by incubation in stripping buffer (80 % formamide, 0.1 % triton-X100) with agitation at room temperature for 3 to 4 × 10 minutes, followed by three 5 minute washes in PBST before the next sequencing round.

After the last round of sequencing and stripping, we incubated the samples with NeuroTrace 530/615 Red Fluorescent Nissl Stain (Thermo Fisher) at a 1:100 dilution in PBST at room temperature for 1 hour and washed 3 × 10 minutes in Washing&Imaging buffer, before acquiring images of the Nissl and DAPI channels for cell segmentation.

#### Data processing

We began data processing by producing maximum z-projections of each acquired dataset, which was made feasible due to the low density of neurons in the CN. We next registered data sets across sequencing and Nissl rounds using the DAPI channel and the Fiji “Register Virtual Stack Slices” plug-in and cropped images to the union of all rounds using custom Fiji scripts. We fed cropped images of the sequencing rounds into the spacetx starfish pipeline (https://github.com/spacetx/starfish; pulled from Github at version alpha 2) for spot/rolony calling using DetectSpots.LacalMaxPeakFinder and decode_per_round_max (min_distance = 2, stringency = 0, minimum object area = 4, maximum object area; code can be found on https://github.com/nbingo/starmap-spacetx). In parallel, we trained a pixel-level classifier in ilastik (Berg et al. 2019) on the Nissl images to classify pixels into “Nissl” and “background”, and applied this classifier to all Nissl images. We fed the resulting ilastik “Nissl” probability maps into a custom Fiji macro, which smoothed the input, applied a threshold, filled holes, and applied a watershed algorithm to segment individual cells. Finally, we combined the segmented cells and starfish identified rolonies in MatLab (R2018b, Mathworks) to generate a cell × gene counts matrix. To avoid noise from incomplete stripping or cross-reactivity between sequencing rounds, we set a minimum signal intensity which a rolony had to exceed to be counted. Note that this cutoff is channel-dependent, as e.g. the 730 channel is weaker than the 647 channel independent of the gene probed for.

#### Cell type calling

Using the Nissl images as a guide, we defined Medial, Interposed, and Lateral CN in every section. To call cell types within each thus defined nucleus we binarized gene expression into “on” and “off” using a general threshold across all genes but fine-tuned this threshold for individual very highly or lowly expressed genes. We then defined cell types by logical combinations of binarized marker genes, guided by snRNAseq data. Generally, we first divided the population of excitatory cells into Class A or Class B cells and then identified individual cell types within these classes using additional marker genes. Cells that expressed either *Slc17a6* or *Gad1*/*Slc6a6* but did not match a specific cell type were labeled as unassigned.

### Viral injections

We performed stereotaxic surgeries and viral injections using standard procedures. Briefly, we anesthetized mice using 1–2 % isoflurane and placed them in a stereotaxic apparatus (Kopf Instruments). We pressure injected AAV virus into specific brain regions at a rate of 3–5 nL/sec using a UMP3 UltraMircoPump (World Surgical Instruments).

For anterograde tracing (Fig. 1) we injected 150 nL *AAV8-CAG-tdTomato* (UNC gene therapy stock vector, Boyden lab Control Vector) into a single site per mouse. We used the following coordinates in the right hemisphere (all relative to lambda); anterior Medial CN: –1.83 mm AP, 0.6 mm ML, 3.3 mm DV (N = 5 mice); posterior Medial CN: –2.18 mm AP, 1.0 mm ML, 3.2 mm DV (N = 5 mice); Interposed CN: –1.83 mm AP, 1.0 mm ML, 3.4 DV (N = 6 mice); Lateral CN: –1.4 mm AP, 2.45 mm ML, 3.7mm DV (N = 7 mice).

For collateralization mapping (Fig. 6C, D) we injected 150 nL of *AAV8-CAG-FLEX-tdTomato* (UNC gene therapy stock vector, Boyden lab Control Vector) into the right Lateral CN (–1.4 mm AP, 2.45 mm ML, 3.7mm DV from lambda) and 150 nL *AAVretro-Ef1a-Cre* (Salk vector core, Addgene #55637) into either left zona incerta (–2.03 mm AP, 2.0 mm ML, 4.0 mm DV from bregma, N = 3 mice) or left brainstem reticular nuclei (parvocellular reticular nucleus, –2.18 mm AP, 1.3 mm ML, 5.5 mm DV from lambda, N = 2 mice; –2.78 mm AP, 1.3 mm ML, 5.5 mm DV from lambda, N = 1 mouse). The center of each retrograde injection site was permanently labeled with co-injected red retrobeads (Lumafluor).

For collateralization mapping as shown in Fig. S24 we injected 5 × 200 nL of *AAV8-CAG-FLEX-tdTomato* (UNC gene therapy stock vector, Boyden lab Control Vector) into the CN at 5 locations (–1.83 mm AP, 0.6 mm ML, 3.3 mm DV; –1.83 mm AP, 1.5 mm ML, 3.4 mm DV; –1.83 mm AP, 2.0 mm ML, 3.4 mm DV; –1.4 mm AP, 2.45 mm ML, 3.7 mm DV; –2.18 mm AP, 1.0 mm ML, 3.2 mm DV; all from lambda). Per animal, we then injected ~200 nL *AAVretro-Ef1a-Cre* (Salk vector core, Addgene #55637) into one site contralateral to the CN injections. Coordinates for these injections are as follows. VL thalamus: –1.07 mm AP, 1.25 mm ML, 3.5 mm DV from bregma; CM thalamus: –1.43 mm AP, 0 mm ML, 3.6 mm DV from bregma; Superior colliculus: –3.27 mm AP, 0.5 mm ML, 1.5 mm DV from bregma; Red nucleus: –3.5 mm AP, 0.6 mm ML, 3.6 mm DV from bregma; Pontine nuclei: –3.9 mm AP, 0.6 mm ML, 5.6 mm from bregma; Vestibular nuclei: –1.5 mm AP, 1 mm ML, 4 mm DV from lambda; Vermis: –1.5 mm AP, 0 mm ML, 1.25 mm DV from lambda; Crus 1: –1.5 mm AP, 3 mm ML, 2.5 mm DV from lambda; Spinal cord: injection between C1 and C2, at 0.8 mm and 0.25 mm depth.

For retrograde tracing followed by STARmap (Fig. 6A, B) we injected 150 nL of *AAVretro-Ef1a-FlpO* (Salk vector core, Addgene #55637) or *AAVretro-Ef1a-Cre* (Salk vector core, Addgene #55636) into right zona incerta (–2.03 mm AP, 2.0 mm ML, 4.0 mm DV from bregma, N = 2 mice per virus) and 150 nL of an *AAVretro:Ef1a:mycH2B* virus (packaged by Stanford Virus Core) with one of two variable 3’UTRs into the right parvocellular reticular nucleus (–2.18 mm AP, 1.3 mm ML, 5.5 mm DV from lambda, N = 2 mice per virus), resulting in an N = 4 independent mice with dual injections into zona incerta and parvocellular reticular nucleus.

All viruses were allowed to express for a minimum of three weeks before proceeding with experiments.

### Brain clearing, whole-brain imaging, and quantification

#### Brain clearing and imaging

For all whole-brain tracing experiments, we transcardially perfused mice with 20 mL 1× PBS (Thermo) containing 10 μg/mL Heparin (Sigma Aldrich), followed by 20 mL 4% paraformaldehyde (Electron Microscopy Sciences) before removing each intact brain, as well as (for Figs. 1, S1-S9, S24) the spinal cord (treated as a brain in what follows) and postfixing at 4 °C overnight. The clearing protocol was largely performed as previously described (Chi et al. 2018; Ren et al. 2019). Briefly, we washed postfixed brains at room temperature in PBS for 3 × 1 hour and then dehydrated them in an ascending methanol gradient [20/40/60/80/100/100 % methanol in B1n buffer (0.1 % Triton-X100, 0.3 M glycine, 0.001 % 10N NaOH)] for 1 hour per step. We delipidated the brains by overnight incubation in 2 : 1 mixture of dichloromethane (Sigma) and methanol, followed by a 1 hour incubation in 100 % dichloromethane. After three washes in 100 % methanol (30 minutes, 45 minutes, and 1 hour long) we bleached the brains in a 5 : 1 mixture of methanol : 30 % H_2_O_2_ (Sigma) for 4 hours at room temperature, and rehydrated the brains in a reverse methanol : B1n gradient (80/60/40/20 %, 30 minutes each). We then washed the brains in B1n buffer for 1 hour and permeabilized them with two washes in PTxwH buffer (1× PBS, 0.1 % Triton-X100, 0.05 % Tween-20, 2 μg/mL heparin, 0.02% NaN_3_) containing 5 % DMSO and 0.3 M glycine for 1 and 2 hours. We washed the brains in PTxwH overnight and performed primary antibody labeling using a rabbit polyclonal anti-RFP antibody (Rockland #600-401-379) at 1:500 in PTwxH at 37 °C for 10 days. We then washed the brains in PTxwH at 37 °C for three days with regular buffer changes and performed secondary antibody labeling using AlexaFluor647 donkey anti-rabbit polyclonal antibody (A-31573, Thermo Fisher) at 1:1000 in PTwxH at 37 °C for 8 days. We then washed the brains in PTwxH at 37 °C for 2.5 days, followed by 2.5 days at 37 °C in PBS. After this washing, we embedded the spinal cords in 2 % low melting point agarose (Sigma) and proceeded with clearing the brains in an ascending methanol : H_2_O gradient (20/40/60/80/100 %, 30 minutes each) at room temperature followed by two additional washes in 100 % methanol (1 hour and 1.5 hours each). We then incubated the brains in a 2 : 1 dichloromethane : methanol mix overnight, followed by three washes in 100 % dichloromethane (30 minutes, 1 hour, 1.5 hours). Finally, we cleared the brains in 100 % dibenzylether, switching the brains to fresh dibenzylether for long term storage after 4 hours.

We imaged brains and spinal cords at least 24 hours after clearing on a LaVision Utramicroscope II light-sheet using a 2× objective and a 3 μm step size taking horizontal optical sections through the brain, with the ventral side facing up. We physically trimmed off the olfactory bulbs and frontal cortex to fit the brain into the field of view of the microscope and imaged the entire remaining brain volume except for the most dorsal and lateral regions of the cerebral cortex. Note that the CN do not project to the trimmed regions. This procedure, therefore, did not result in a loss of projection data. Antibody signal was collected in the 647 channel with 28 horizontal focusing steps to homogenize z-resolution across the field. The autofluorescence signal and retrobead injection site signal were collected in the 488 and 561 channel, respectively, without horizontal focusing.

#### Axon quantification

We classified axons in our whole-brain imaging datasets using a hybrid strategy using the 3D U-Net convolutional network TrailMap (Friedmann et al. 2020) and an ilastik pixel-level classifier (Berg et al. 2019). After adjusting pixel values by multiplication with a scalar to match the background levels seen in the TrailMap training dataset, TrailMap was very good at detecting most axons but missed axons and fiber bundles with the highest signal-to-noise ratio. We, therefore, trained a simple ilastik pixel-level classifier to detect these very bright axons and combined the TrailMap and ilastik probability maps by a maximum operation. The resulting hybrid probability map faithfully captured CN axons in our data sets.

We then aligned the autofluorescence channel of our brains to the Allen Institute’s Common Coordinate framework (Renier et al. 2016) using elastix (Klein et al. 2009). As reference brain, we used a custom version of the Allen STP reference brain at 25 μm resolution in which we changed the pixel values of all fiber tract annotated regions to be bright (as they are in Adipocleared brains), rather than dark (as they are in serial 2-photon tomography), which greatly improved alignment in the brainstem and where ever axon bundles closely abutted the (always dark) regions outside the brain. We applied the same transformation to the detected axon volumes returned by the TrailMap/ilastik pipeline, thresholded the probability maps, and quantified the number and density of axonal voxels in each annotated brain region in Matlab normalizing each brain to the total number of axon containing voxels (R2018b, Mathworks). In this analysis, we ignored the incompletely imaged isocortex and striatum, as well as any voxels in the ‘root’, fiber tract, corpus callosum, and ventricle annotation. In some brains, we observed spurious, sparse innervation of the ipsilateral cortex and deep layers of the contralateral cortex, as well as innervation of the claustrum. These projections are likely caused by virus leak along the cerebellar peduncle out of the CN. As a result, we removed the spuriously innervated claustrum from our analysis. In the ipsilateral hemisphere, we further removed the CN as the injection sites. In total, we quantified 242 and 246 brain regions in the right and left hemispheres, respectively.

To produce brain-wide projection heat maps, we thresholded the aligned TrailMap/ilastik output at 25 μm resolution from all replicate brains, and normalized each brain by the total number of axonal voxels. We then summed all brain volumes and divided each voxel by the number of summed brains. Finally, we smoothed the output by a 9 voxel box filter. No voxels except for ventricles (which contained distracting background signal) were blanked.

To derive the projection probability maps of Lateral CN Class A and Class B neurons, we first derived probability maps of zona incerta and reticular nucleus projecting Lateral CN neurons. We summed the aligned and thresholded TrailMap/ilastik output at 25 μm resolution from all replicate brains, divided voxel values by the number of summed brains, and smoothed the output by a 9 voxel box filter. No voxels except for ventricles (which contained distracting background signal) were blanked. We then assumed that *p*(*ZI projecting*) = *xp*(*ClassA*) + (1 − *x*)*p*(*ClassB*) and conversely, *p*(*Ret projecting*) = *yp*(*ClassA*) + (1 − *y*)*p*(*ClassB*), where 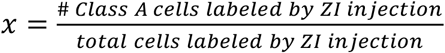 and 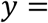 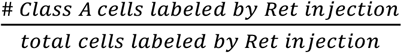. Importantly, we can determine *x* and *y* from the retrograde STARmap experiments. Deriving *p*(*A*) and *p*(*y*) is then simply a question of solving a quadratic equation. This derivation assumes homogeneous projection probabilities to zona incerta and brainstem reticular nuclei within the population of lateral CN Class A and Class B neurons.

### Second-order projection tracing *in silico*

To determine the brain-wide projections of thalamic voxels innervated primarily by Class A or Class B Lateral CN neurons, we used the Allen voxel scale connectivity model (Knox et al. 2019) (https://github.com/nbingo/Mouse-voxel-connectivity-simplified). First, we determined which voxels in the thalamus are primarily innervated by Lateral CN Class A or Class B neurons by selecting voxels in the 25 μm resolution space for which P(A) – P(B) > 0.3 or P(B) – P(A) > 0.3, respectively. Similarly, we selected voxels primarily innervated by ZI-projecting or Ret-projecting Lateral CN neurons by performing pixel-wise two-sample t-tests and selecting pixels for which p < 0.01. We then downsampled the binarized brain with these voxels labeled to 100 μm resolution and used positive thalamic 100 μm voxels as “injection” voxels in the Allen connectivity model. We finally normalized the resulting projection probability maps by the number of “injection” voxels, as the final probability map is the sum of each “injection” voxel’s projection probability map. For region-level quantification, we summed up all voxel values in each region and divided them by the number of voxels in the region. In total, we quantified 261 regions per hemisphere covering the entire brain except for thalamus, which we removed as the *in silico* equivalent of the injection site.

## Acknowledgments

We thank Lauren O’Connell, Hunter Fraser, Jan Lui, Colleen McLaughlin, Jiefu Li, and Hongjie Li for comments on the manuscript; all members of the Luo lab for technical support, advice, and comments on this study; as well as Fabio Zanini, Felix Horns, Geoff Stanley, Xiao Wang, and Jessica Chang for technical advice. We express our thanks for the cooperation of Donor Network West and all of the organ and tissue donors and their families, for giving the gift of life and the gift of knowledge, by their generous donation.

## Author contributions

J.M.K. and L.L. designed the study; J.M.K. performed most of the experiments and data analyses; N.R. assisted in computational analysis; E.B.R. and W.E.A assisted in STARmap experiments with support from K.D.; E.B.R. also assisted in retrograde STARmap experiments; D.F. assisted in whole-brain axon mapping experiments; S.S.K. and R.C.J. assisted in single-nucleus RNAseq experiments with support from S.R.Q.; Y.W. and H.Z. contributed chicken samples; S.W.C. and H.Y.C. contributed RNAseq reagents; J.M.K. and L.L. wrote the paper with feedbacks from all authors. L.L. supervised the project.

## Data availability

The sequencing datasets generated in this study will be available in the NCBI Gene Expression Omnibus. Custom analysis code is available at https://github.com/justuskebschull/CN_code unless otherwise indicated. All other data are available via reasonable requests from the corresponding authors.

