## Supplemental Figures and Table S1 for "Cerebellar nuclei evolved by repeatedly duplicating a conserved cell type set"

| Term | | Definition |
|---|---|---|
| 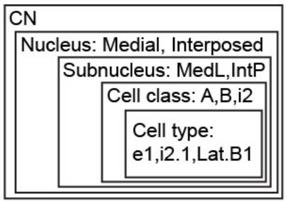 | | Hierarchical structure of terms used in this study, with examples |
| **CN** | | Cerbellar nuclei, used to refer to the complete system |
| **Nucleus** | | Medial, Interposed, or Lateral CN |
| | Medial CN | Most medial cerebellar nucleus; considered phyogenetically oldest |
| | Interposed CN | Cerebellar nucleus located in between Medial and Lateral cerebellar nulcei |
| | Lateral CN | Most lateral cerebellar nucleus; considered phylogenetically youngest |
| **Subnucleus** | | Cytoarchitectually separable part of a nucleus |
| | Med | Central part of the Medial CN |
| | MedL | Lateral part of the Medial CN |
| | MedDL | Dorsolateral protrusion of the Medial CN |
| | IntP | Posterior part of the Interposed CN |
| | IntA | Anterior part of the Interposed CN |
| | Lat | "Subnucleus" that covers the entire Lateral CN |
| **Cell class** | | A group of transcriptomically similar cell types |
| | i1 | Class 1 of inhibitory neurons; putatively IO projecting |
| | i2 | Class 2 of inhibitory neurons; small, likely glycinergic in mouse |
| | i3 | Class 3 of inhibitory neurons; larger, likely glycinergic in mouse |
| | Class A | One of the two classes of excitatory neurons |
| | Class B | The other excitatory neuron class; more detected genes per cell, and likely larger, than Class A neurons |
| **Cell type** | | A group of transcriptomically similar cells |
| | i1.1,i2.1, etc. | Inhibitory cell types belonging to class 1, 2, etc. |
| | e1, etc. | Temporary names for excitatory mouse cell types before we define their location |
| | Lat.B1, etc. | Excitatory cell type of ClassB that is located in the Lat subnucleus, etc. |
| **Sibling cell types** | | Cell types with a common evolutionary origin; examples are the cell types in Class A, or the cell types in ClassB |
| **Stereotyped cell type set** | | A group of cell types comprising members of inhibitory classes i1, i2, i3, and excitatory classes A and B |
| **Archetypal CN** | | A subnucleus that is formed by the stereotyped cell type set;<br>Proposed unit for CN organization and duplication during evolution |



**Table S1:** Key nomenclature used in this study and their relationship.

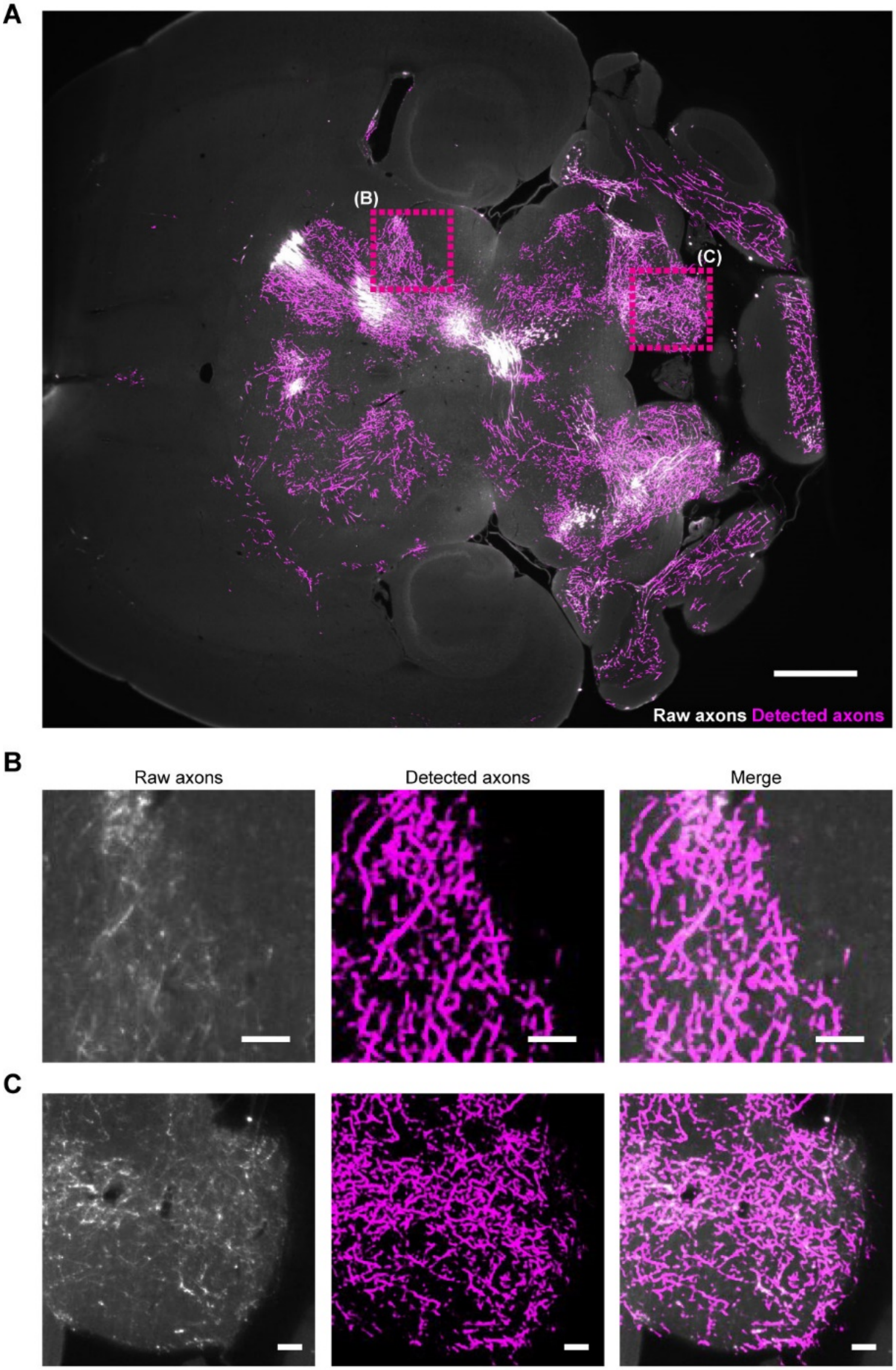


**Figure S1: Axon detection by a hybrid TrailMap-ilastik pipeline.** (A) Representative horizontal section of raw data overlaid with axon probability maps returned by the classification pipeline. Scale bar = 1 mm. (B, C) Zoom in on indicated areas, showing raw data, detected axon probability maps, and merged images. Scale bar = 100 μm.

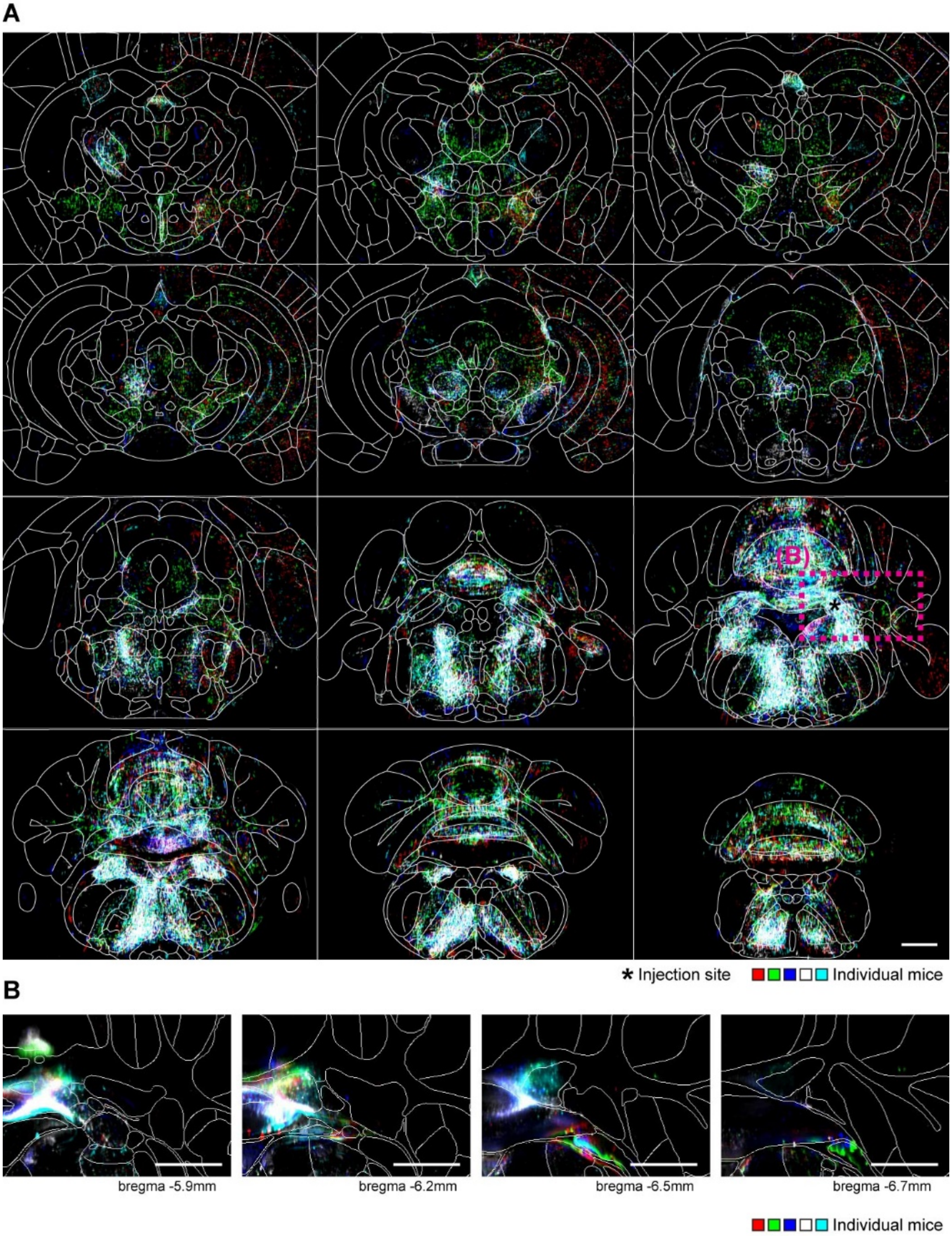


**Figure S2: Brain-wide projections from anterior Medial CN.** (A) Overlay of detected axons for N = 5 independent injections. Coronal sections are shown, spaced 625 μm apart, along the A–P axis, with Allen compartments overlaid in white. Scale bar = 1 mm. (B) Overlay of raw data for these 5 brains, showing the injection sites in coronal sections at the indicated A–P positions. Scale bar = 1 mm. Each mouse is shown in a unique color that matches across panels.

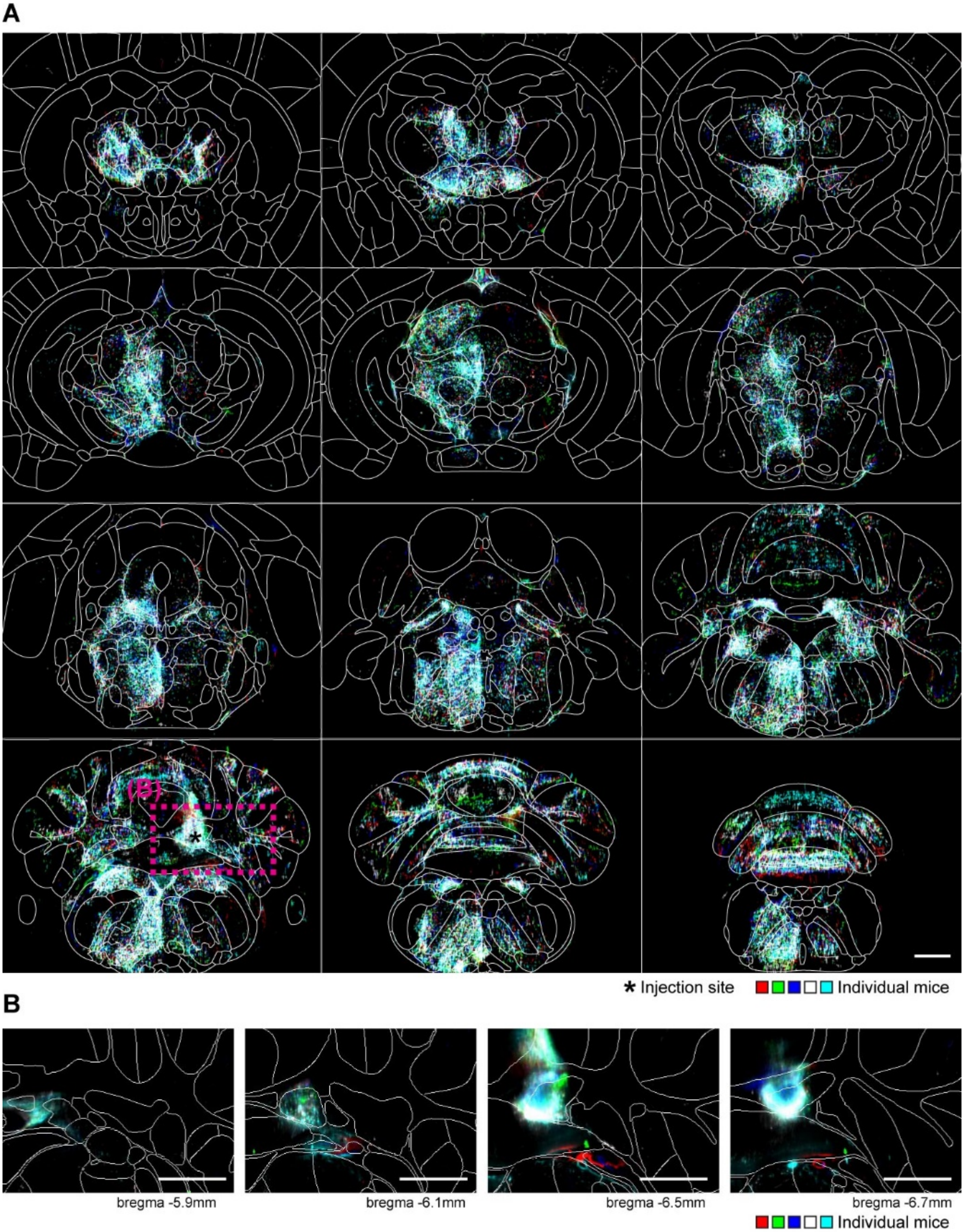


**Figure S3: Brain-wide projection from posterior Medial CN.** See caption for Fig. S2. N = 5.

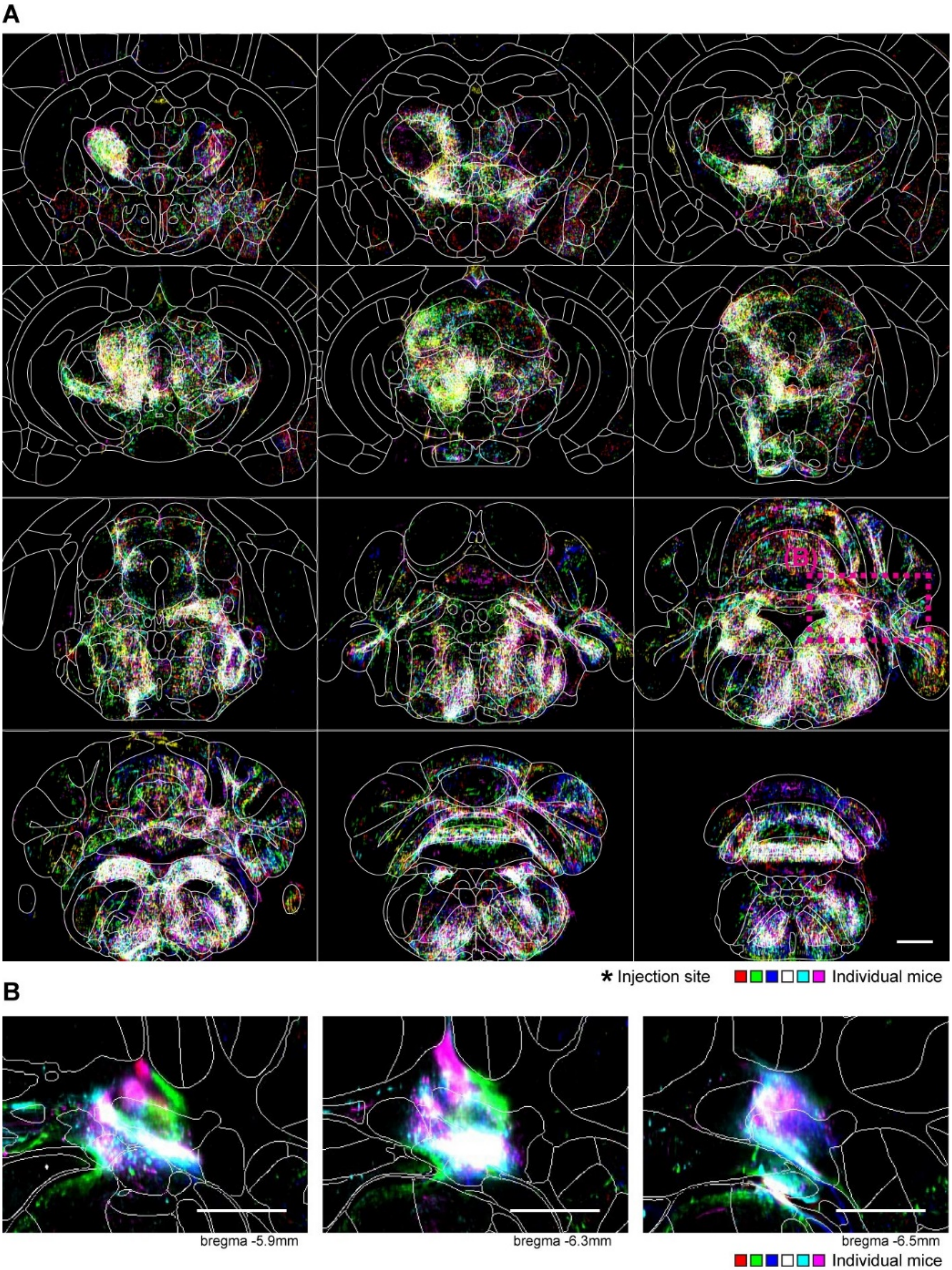


**Figure S4: Brain-wide projection from Interposed CN.** See caption for Fig. S2. N = 6.

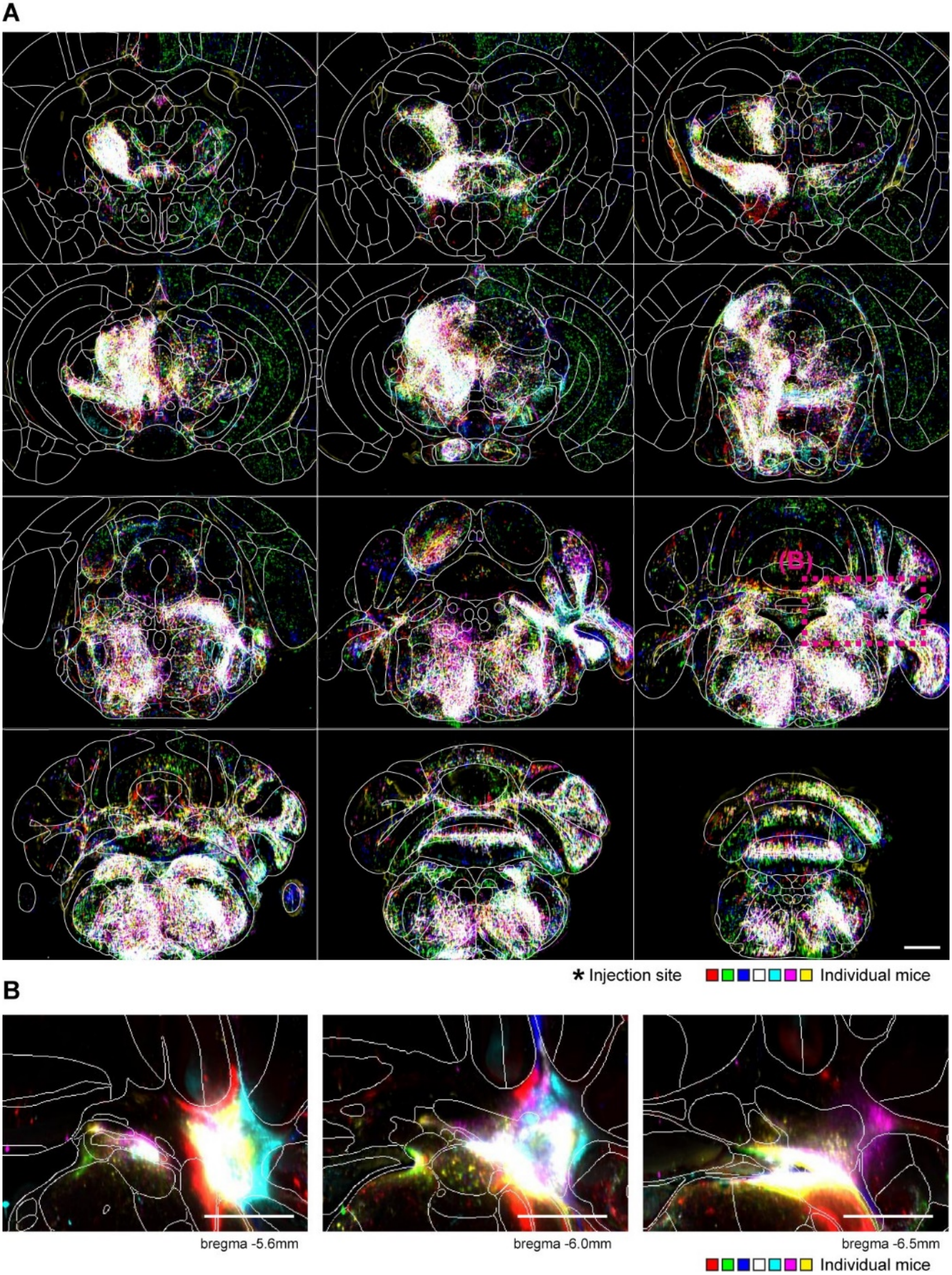


**Figure S5: Brain-wide projection from Lateral CN.** See caption for Fig. S2. N = 7.

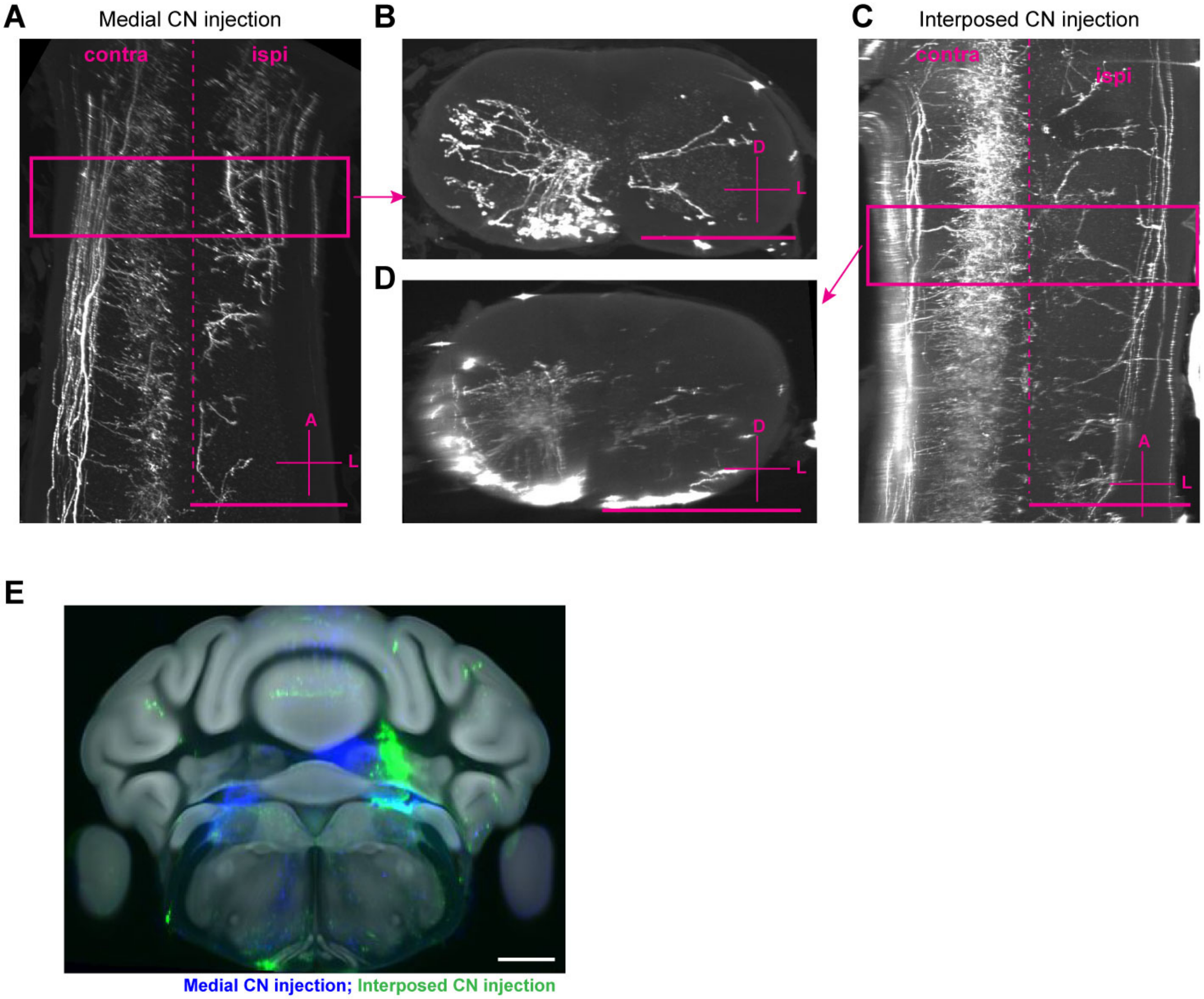


**Figure S6: Spinal cord projections of mouse CN.** (A) Representative horizontal section (maximum z-projection over 300 µm) through the cervical spinal cord of a Medial CN injected animal. Scale bar = 1 mm. (B) Coronal z-projection over 300 µm as indicated by the box in (A). Innervation of the contralateral ventral horn is apparent. Scale bar = 1 mm. (C, D) Innervation of the cervical spinal cord by Interposed CN, as in (A,B). For both Medial and Interposed CN, a few bright fibers continued past the cervical spinal cord but were never seen to innervate the spinal cord gray matter at any level. Lateral CN injections result in no spinal cord labeling (data not shown). (E) Injection sites for the two brains shown in (A–D). Medial and Interposed CN injections are well separated. Scale bar = 1 mm.

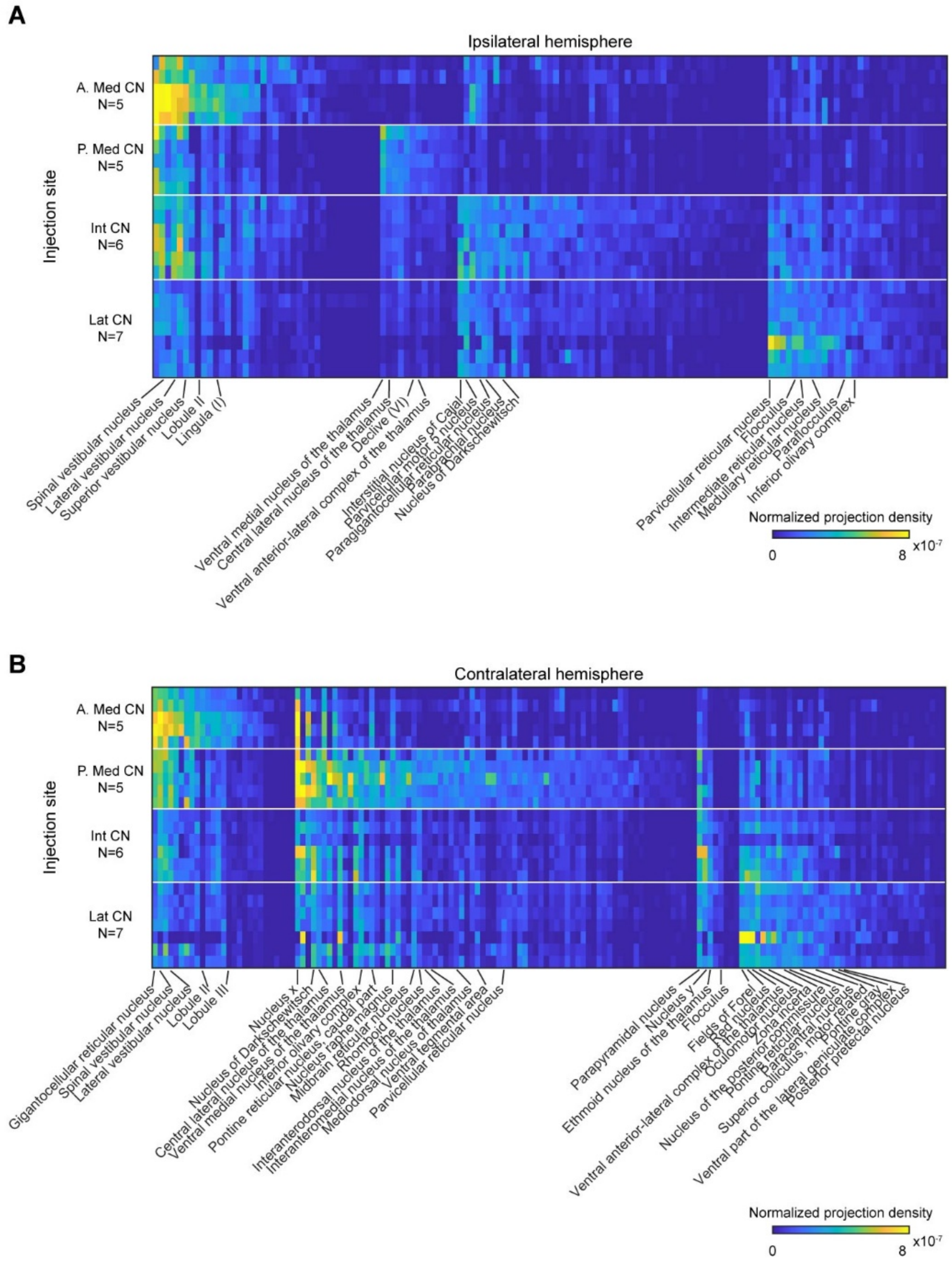


**Figure S7: Brain-wide differences in the projections of the moues CN.** (A,B) Heat map of all brain regions significantly differently innervated by anterior Medial, posterior Medial, Interposed or Lateral CN ($p < 0.05$, 1 way ANOVA) on the (A) ipsi- or (B) contralateral side. Brain regions are sorted by their mean projection within each injection site. Medial CN have a more distinct projection pattern than Interposed and Lateral CN. Intensity values indicate normalized innervation densities per animal.

A

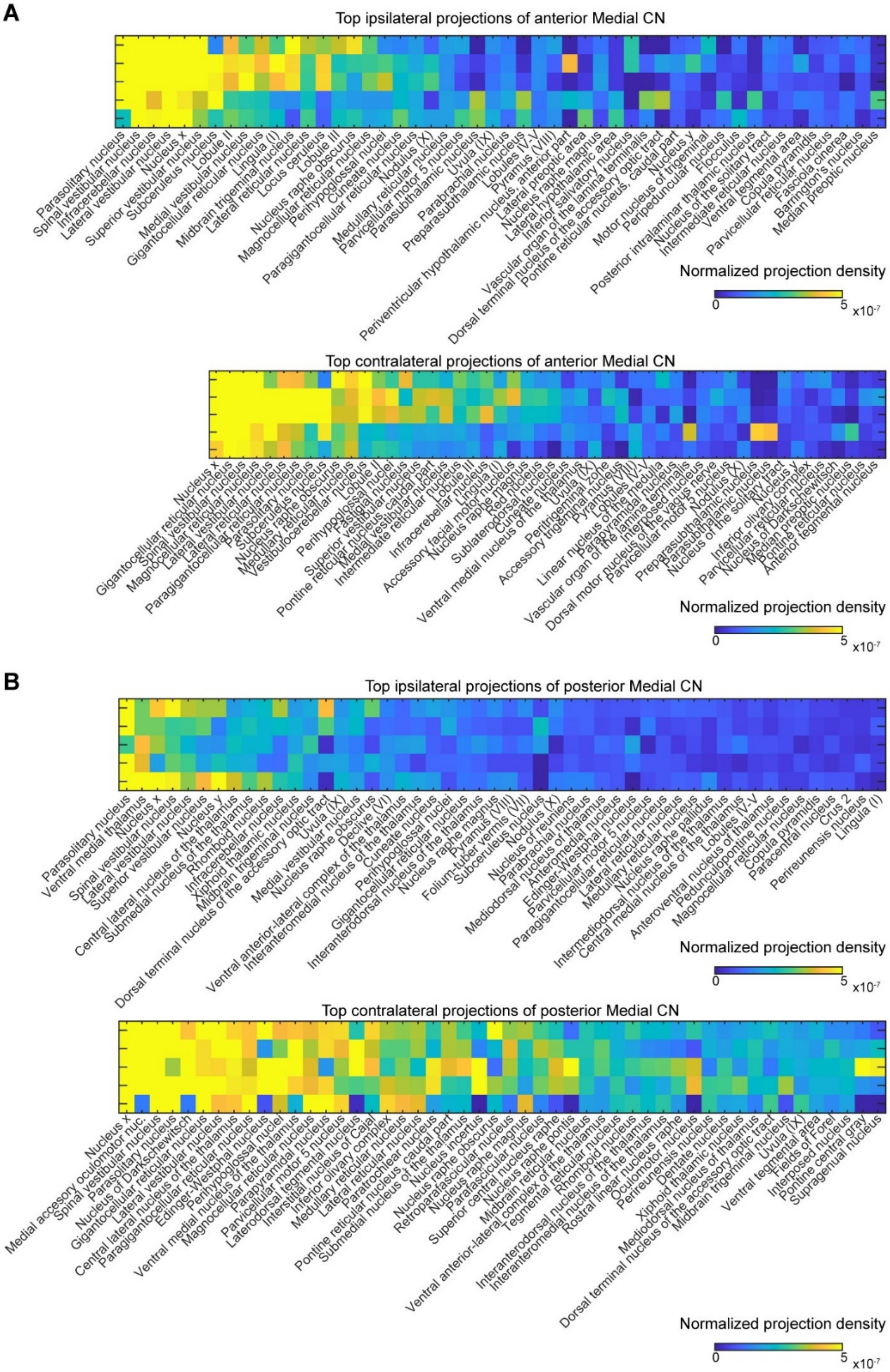


**Figure S8: Quantification of Medial CN projection targets.** (A, B) Ipsi- (top) and contralateral (bottom) normalized innervation densities into the top 50 target regions of anterior Medial (A) and (B) posterior Medial CN (B).

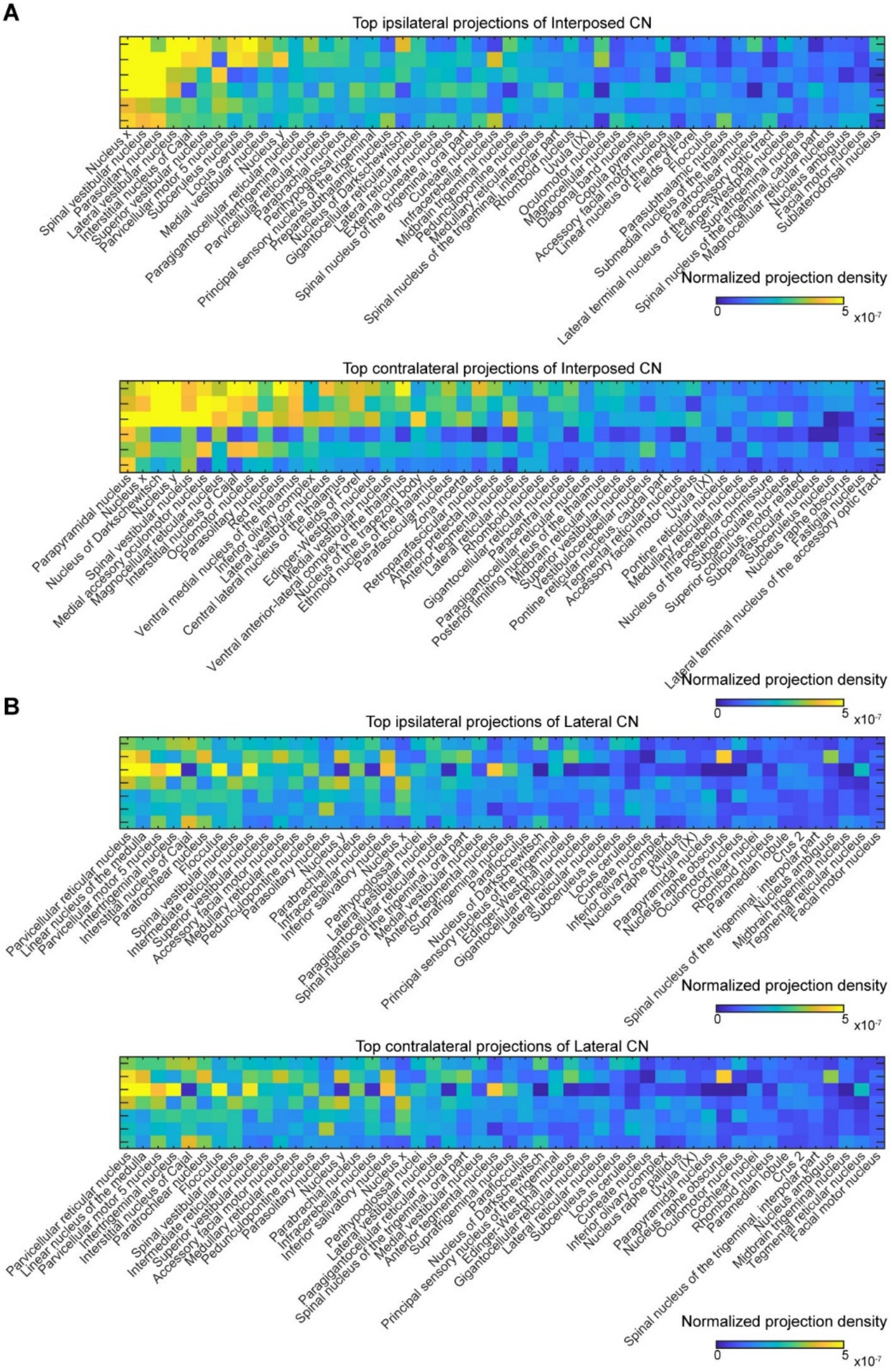


**Figure S9: Quantification of Interposed and Lateral CN projection targets.** (A, B) Ipsi- (top) and contralateral (bottom) innervation densities into the top 50 target regions of Interposed (A) and Lateral CN (B).

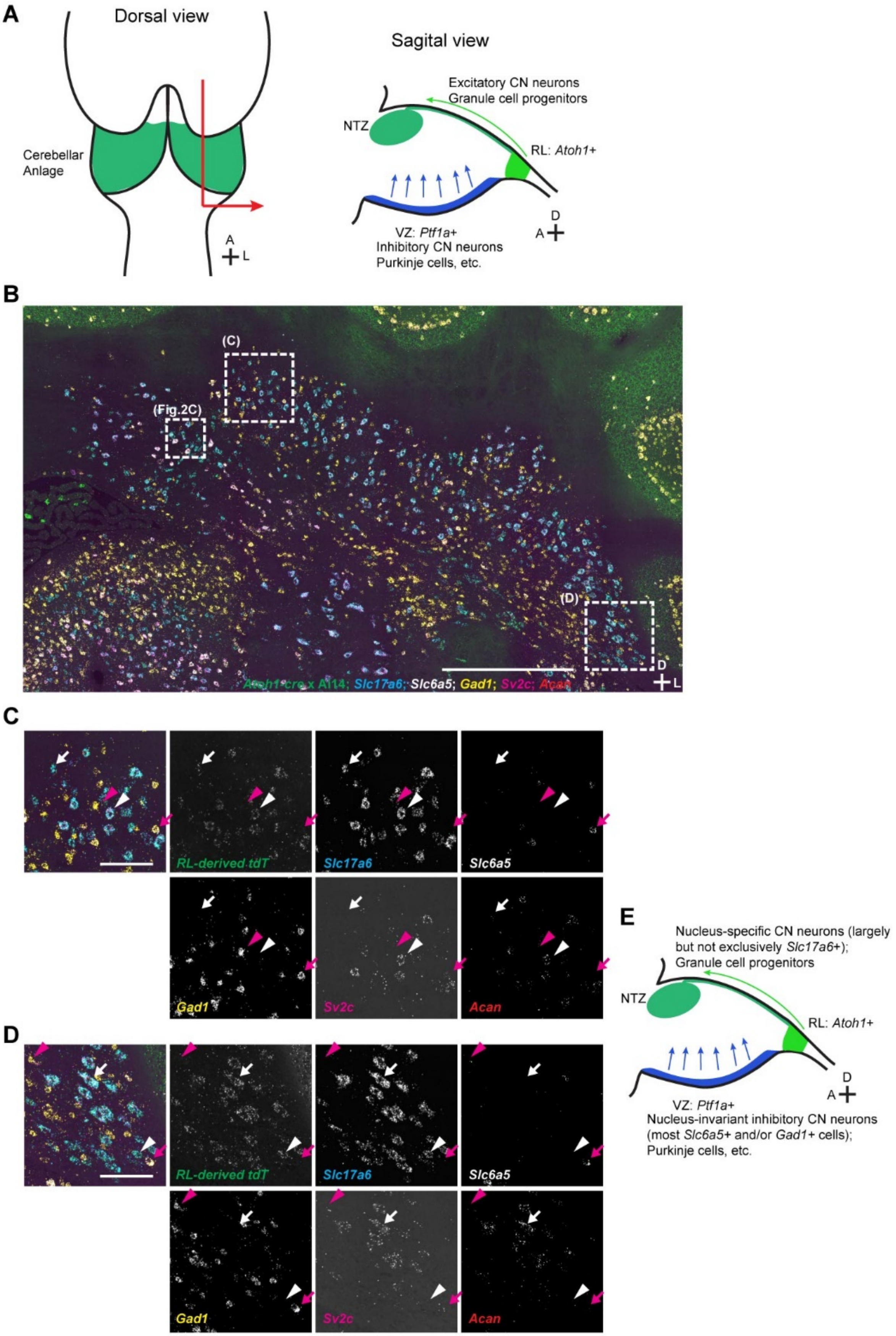


**Figure S10: Developmental origin of mouse CN cell types.** (A) Schematic of the current model of CN development. Left, dorsal view of ~E12 mouse brain; right, sagittal section through cerebellar anlage. Excitatory CN neurons are born from *Atoh1*+ progenitors in the rhombic lip, and migrate into the nuclear transitory zone (NTZ). Inhibitory CN neurons derive from the ventricular zone. (B) Representative STARmap image of *Atoh1-Cre* × Ai14 cerebellar nuclei. *Slc17a6* labels excitatory cells, *Gad1* inhibitory cells, *Slc6a5* glycinergic cells, and *Acan* and *Sv2c* label, in combination with *Slc17a6*, Class B cells. e9* neurons are *Slc6a5*+, *Acan*+, *Sv2c*+; N = 2 sections. Scale bar = 1 μm. (C, D) Magnified view of ventral Lateral CN (C) and Interposed CN (D). In all cases *tdTomato*+ Class A neurons (white arrowhead) and B neurons (white arrow), as well as *tdTomato*– inhibitory (magenta arrowhead) and glycinergic neurons (magenta arrow) are visible. (E) Updated schematic of CN development. RL-derived cells include all excitatory and glycinergic e9* neurons that are CN-specific, whereas VZ-derived cells are all inhibitory and common to all CN. Scale bars = 100 μm.

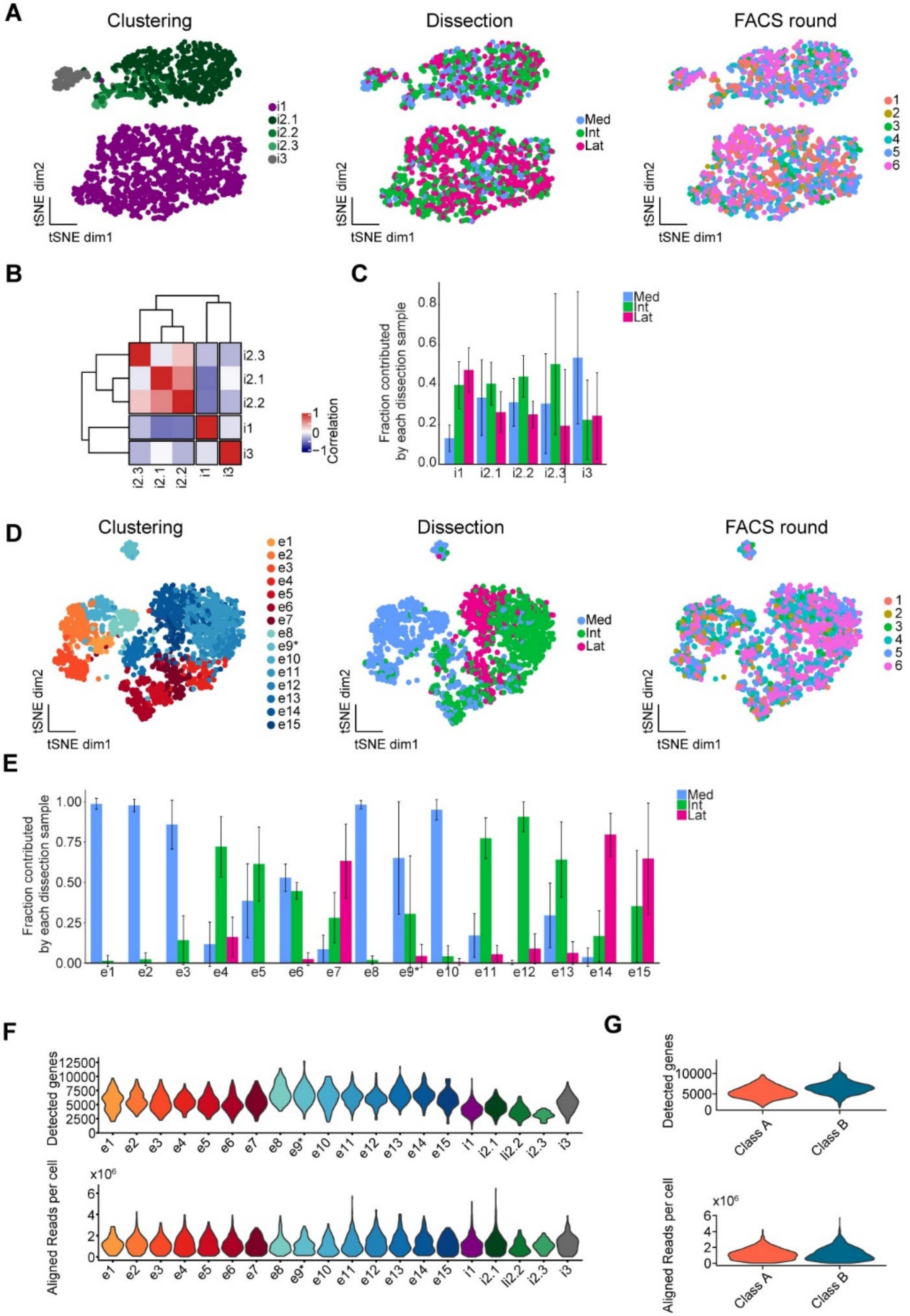


**Figure S11: Details on mouse CN snRNAseq data.** (A) Clustering of inhibitory cell types is independent of potentially confounding factor FACS round/date of collection. (B) Mouse inhibitory neurons fall into three classes, with major cell type i2.1 being more similar to rare cell types i2.2 and i2.3 than to cell types i1 and i3. (C) The proportion of inhibitory cell types collected in Medial (Med), Interposed (Int), or Lateral (Lat) CN dissections across FACS rounds is relatively even, except for fewer Medial CN cells in cluster i1. Mean ± SD is shown. (D) Clustering of excitatory cell types is independent of FACS round/date of collection. (E) Proportion of excitatory cell types collected in Medial, Interposed, or Lateral CN dissections across FACS rounds shows strong biases for individual nuclei contributing differentially to different cell types. (F, G) The number of detected genes and aligned reads per cell broken down by (F) cluster or (G) class.

**A**

| Cell type from this study | Tenative match to reviously reported cell type | Citation |
|---|---|---|
| e7 (Lat.A1) | GADnS | Uusiaari and Knopfel 2011 |
| e14/e15 (Lat.B1/2) | GADnL | |
| i3 | Gly-I | |
| i1 | IO-projecting GABAergic neurons | |
| e9* (MedL.Bgly) | Medial nucleus glycinergic neurons | Bagnall *et al.* 2009 |

**B**

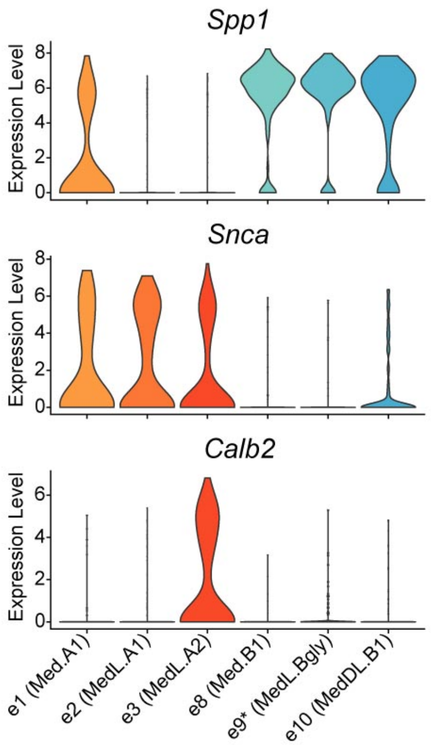


**Figure S12: Mapping of CN cell types in the literature.** (A) Table containing the tentative mapping of cell types obtained by unbiased clustering of transcriptomic data in this work (see Fig. 3G for the renaming of excitatory cell types in parenthesis) to cell types described in the literature. (B) Fujita *et. al* (Fujita, Kodama, and Lac 2020) cell type marker gene expression maps one-to-one to the Medical CN cell types defined in this study by unbiased clustering.

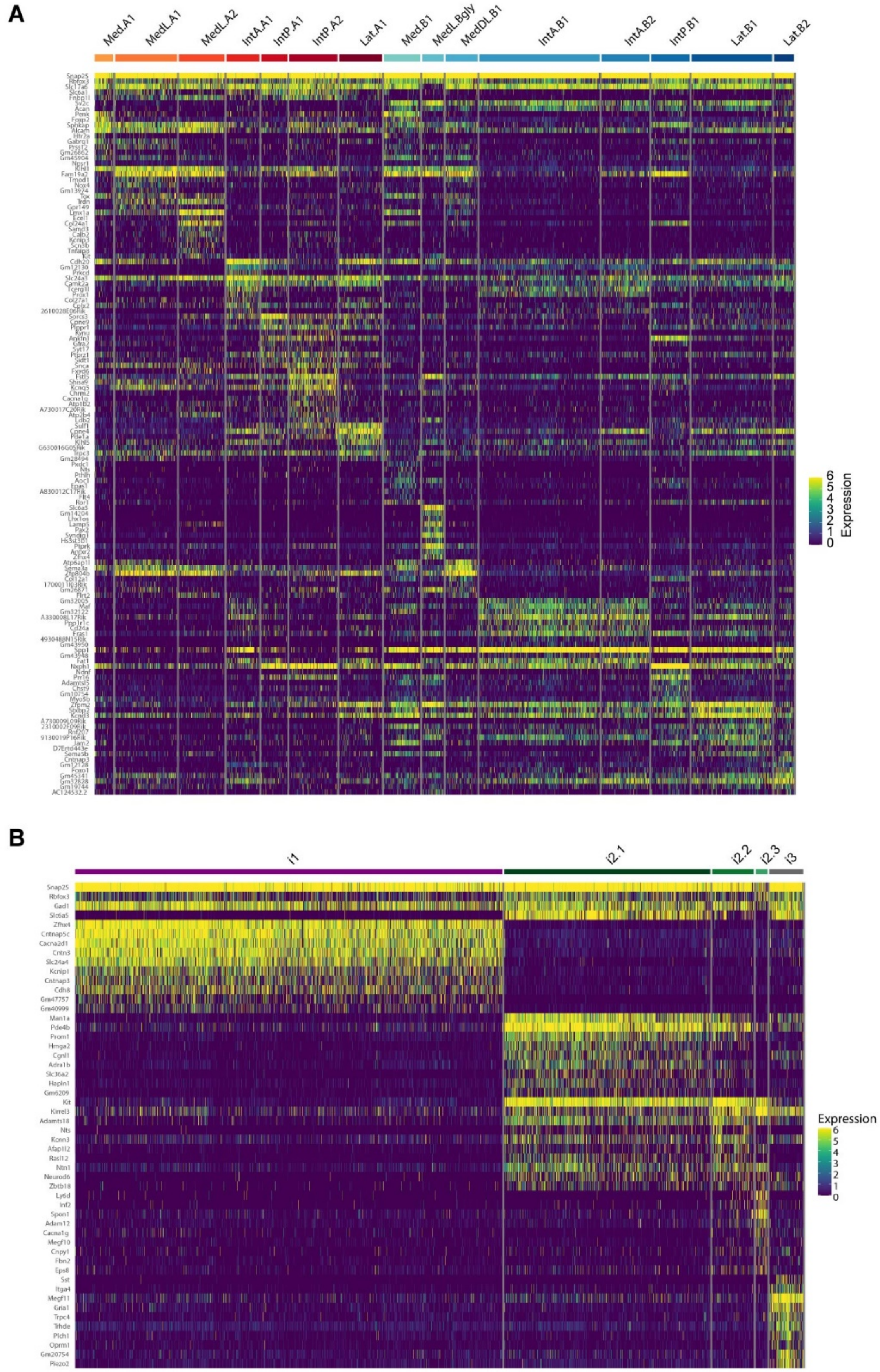


**Figure S13: Differential gene expression in mouse CN.** Heat map of differentially expressed genes for (A) excitatory and (B) inhibitory cell types.

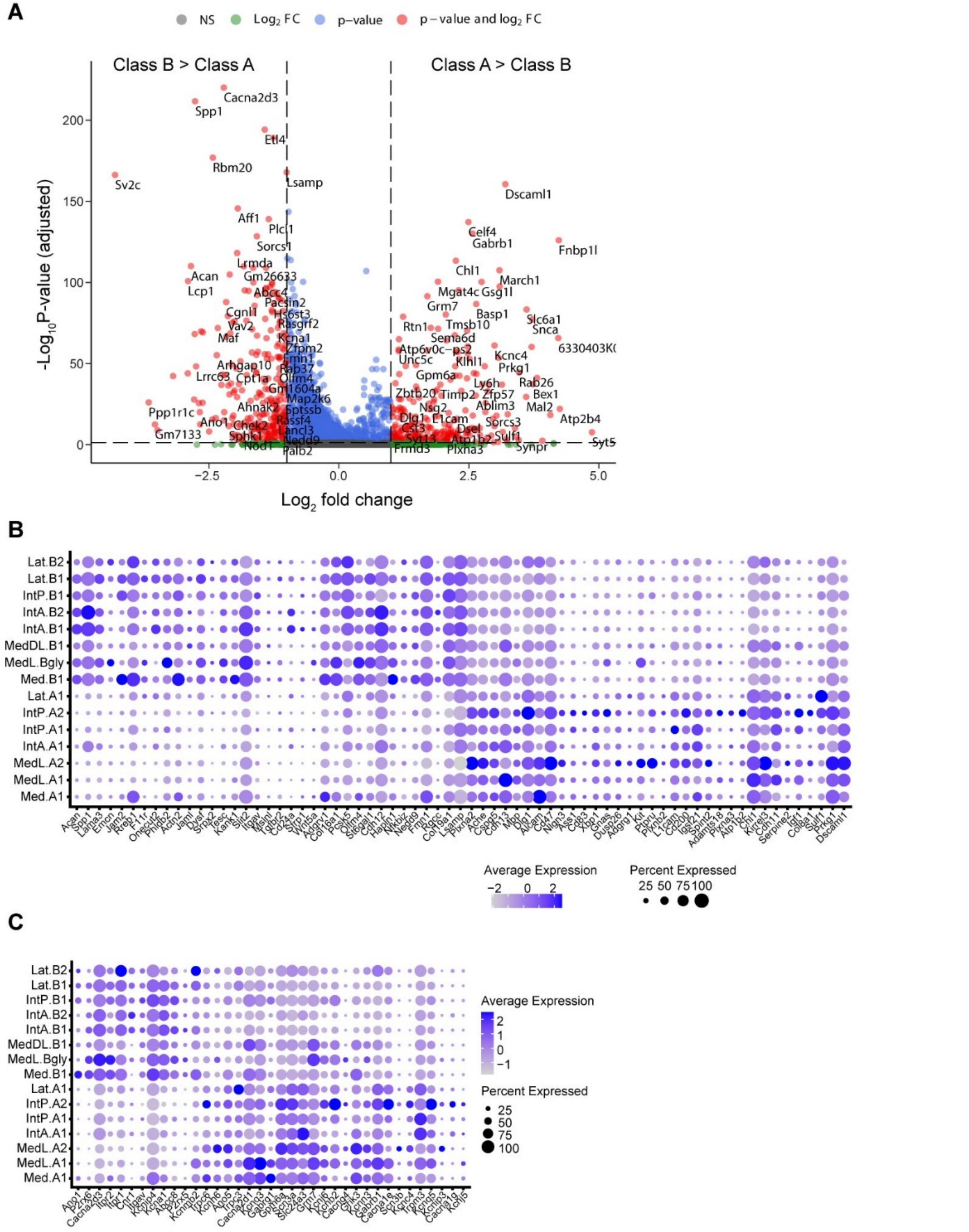


**Figure S14: Differential gene expression between Class A and Class B neurons, part 1.** (A) Volcano plot showing all differentially expressed genes between Class A and Class B neurons after collapsing cell types by class assignment. (B, C) Cluster level expression of Class A vs B differentially expressed genes with (B) cell adhesion (GO:0007155) and (C) ion channel activity (GO:0005216).

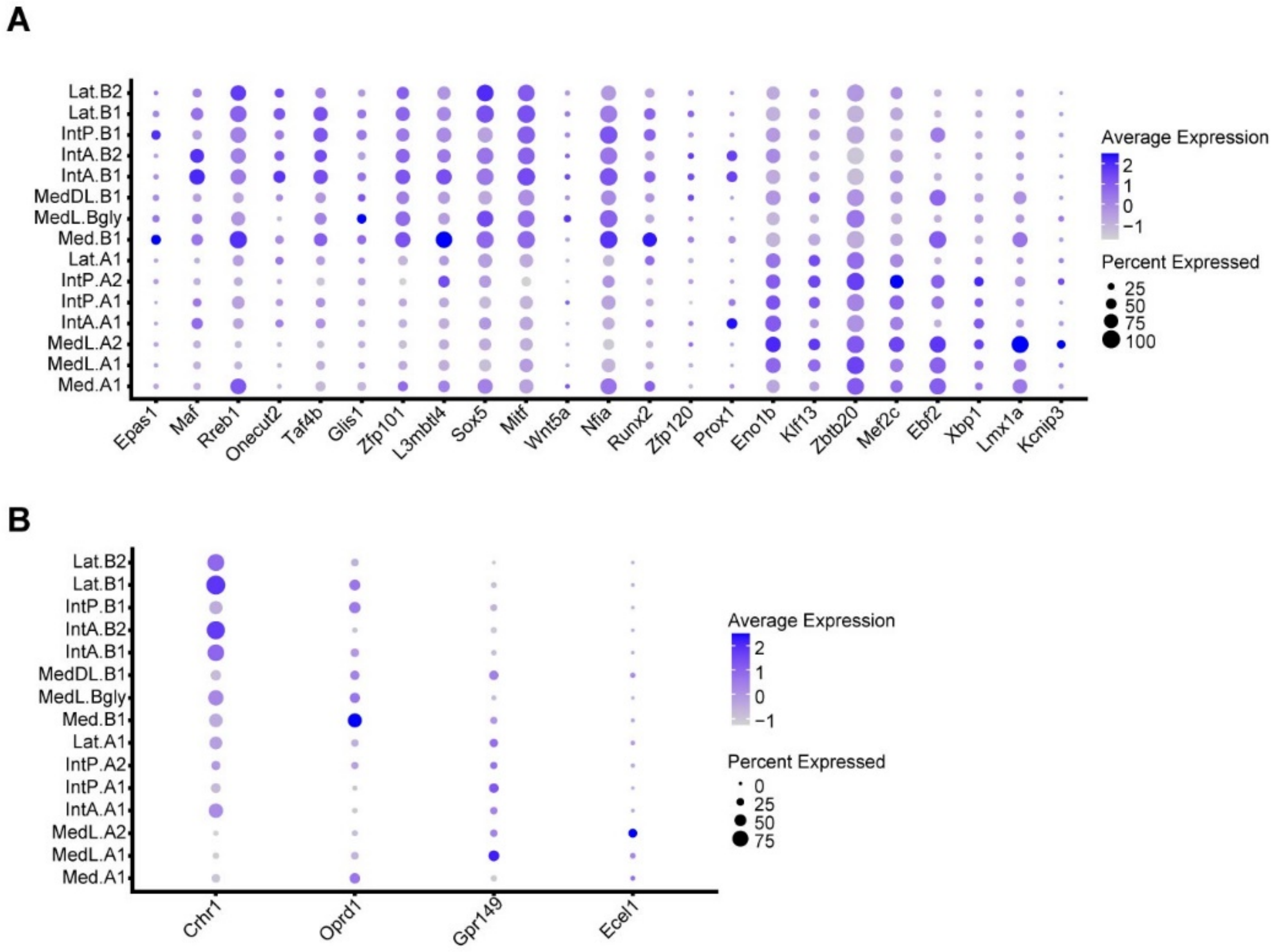


**Figure S15: Differential gene expression between Class A and Class B neurons, part 2.** Cluster level expression of Class A vs B differentially expressed (A) transcription factors (GO:0003700; GO:0003702; GO:0003709; GO:0016563; GO:0016564) and (B) genes with neuropeptide signaling peptide pathway activity (GO:0007218).

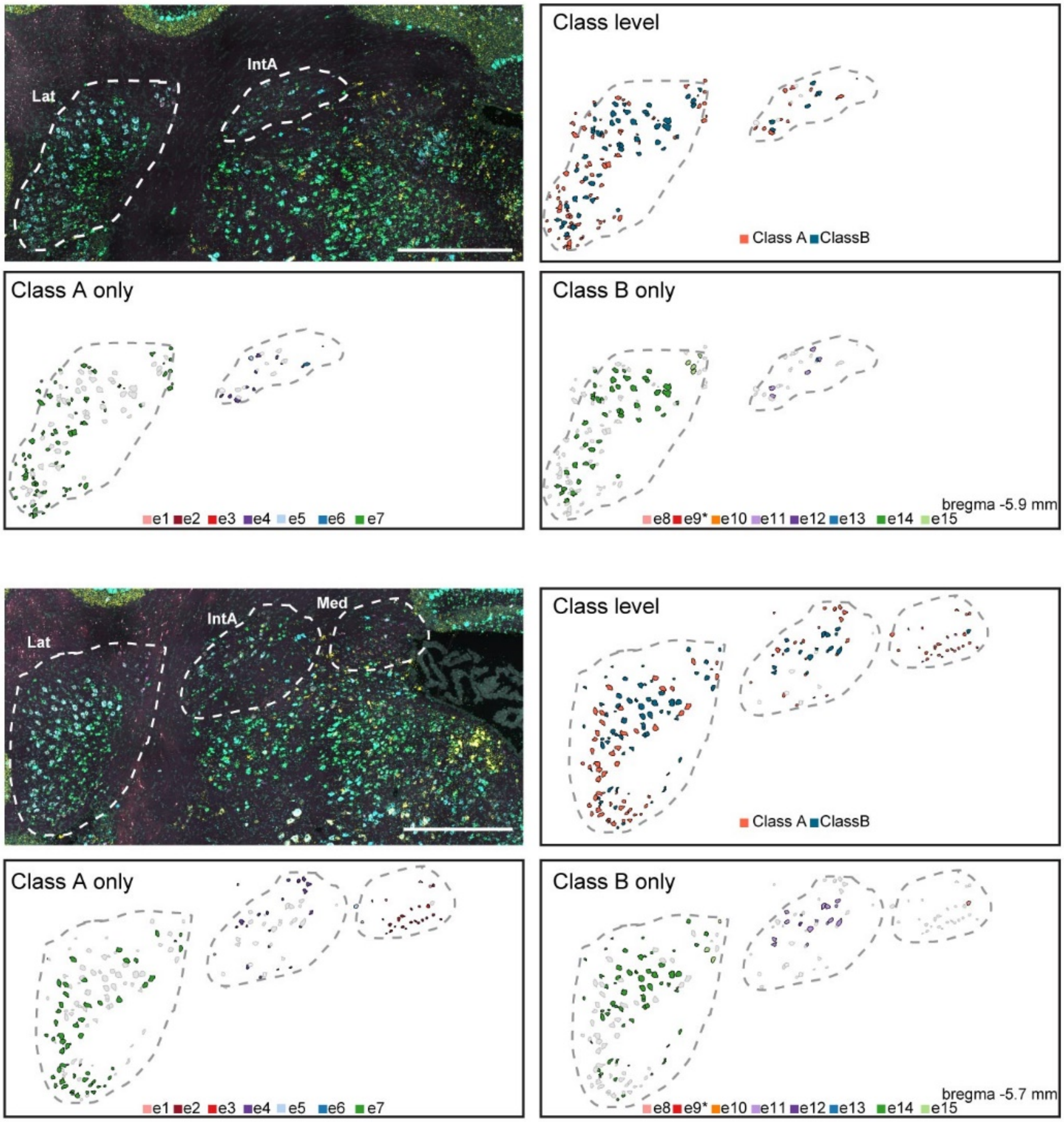


**Figure S16: Spatial location of CN excitatory cell types, part 1.** Representative coronal sections of the CN moving from rostral to caudal illustrating the distribution of excitatory cell classes and of individual cell types within these. Excitatory cells are segmented based on Nissl images and are assigned to a cell type based on gene expression measured by sequential STARmap readout of 12 core marker genes per section. Cell type assignment is based on criteria derived from snRNAseq. Excitatory cells are colored by class/cell type. Cells of the other class and unassigned excitatory cells are shown in light grey. N = 2 mice. Scale bars = 500 µm. Fig. S17 is a continuation of this figure representing more caudal sections.

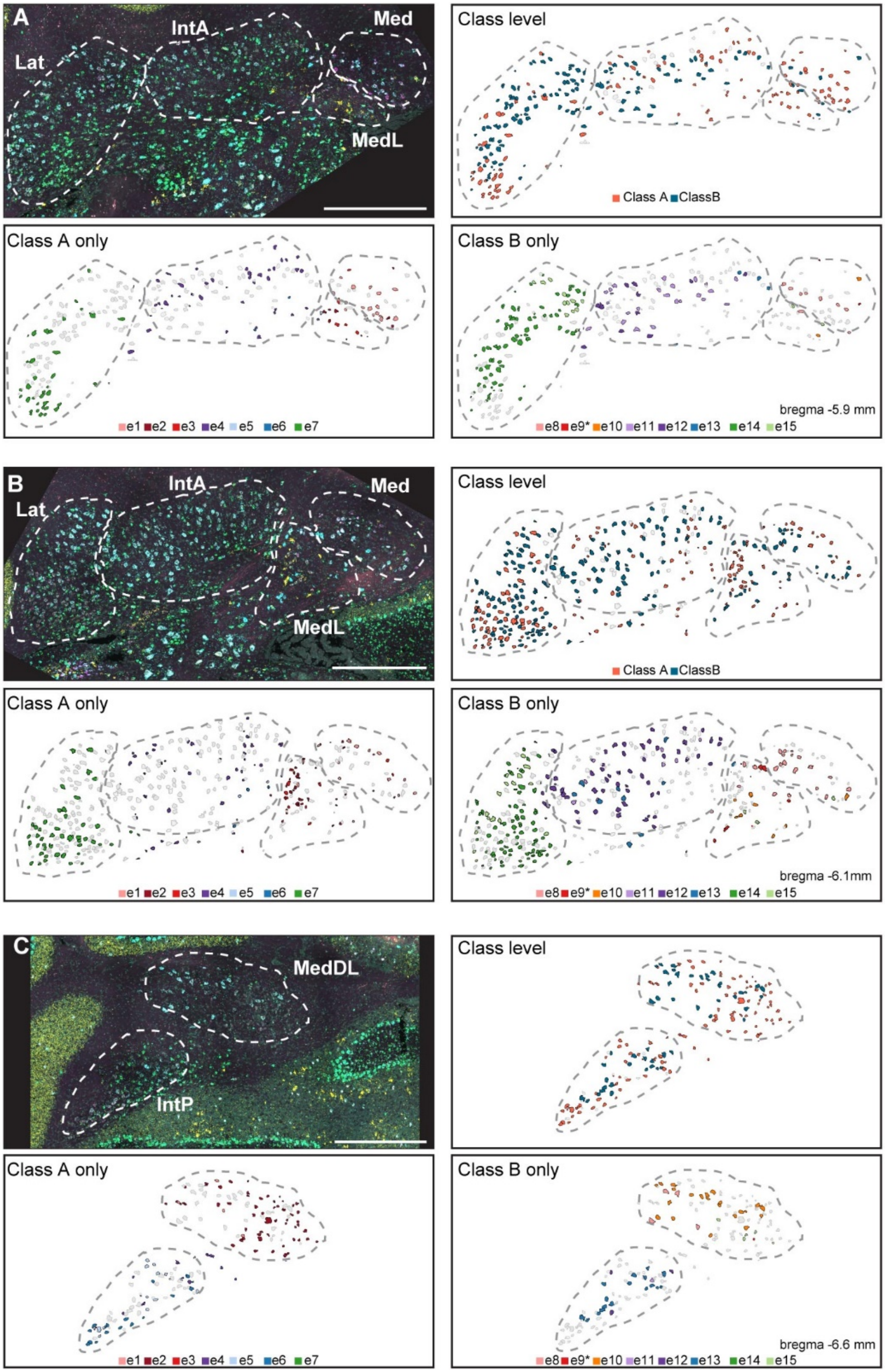


**Figure S17: Spatial location of CN excitatory cell types, part 2.** see Figure S16 legend.

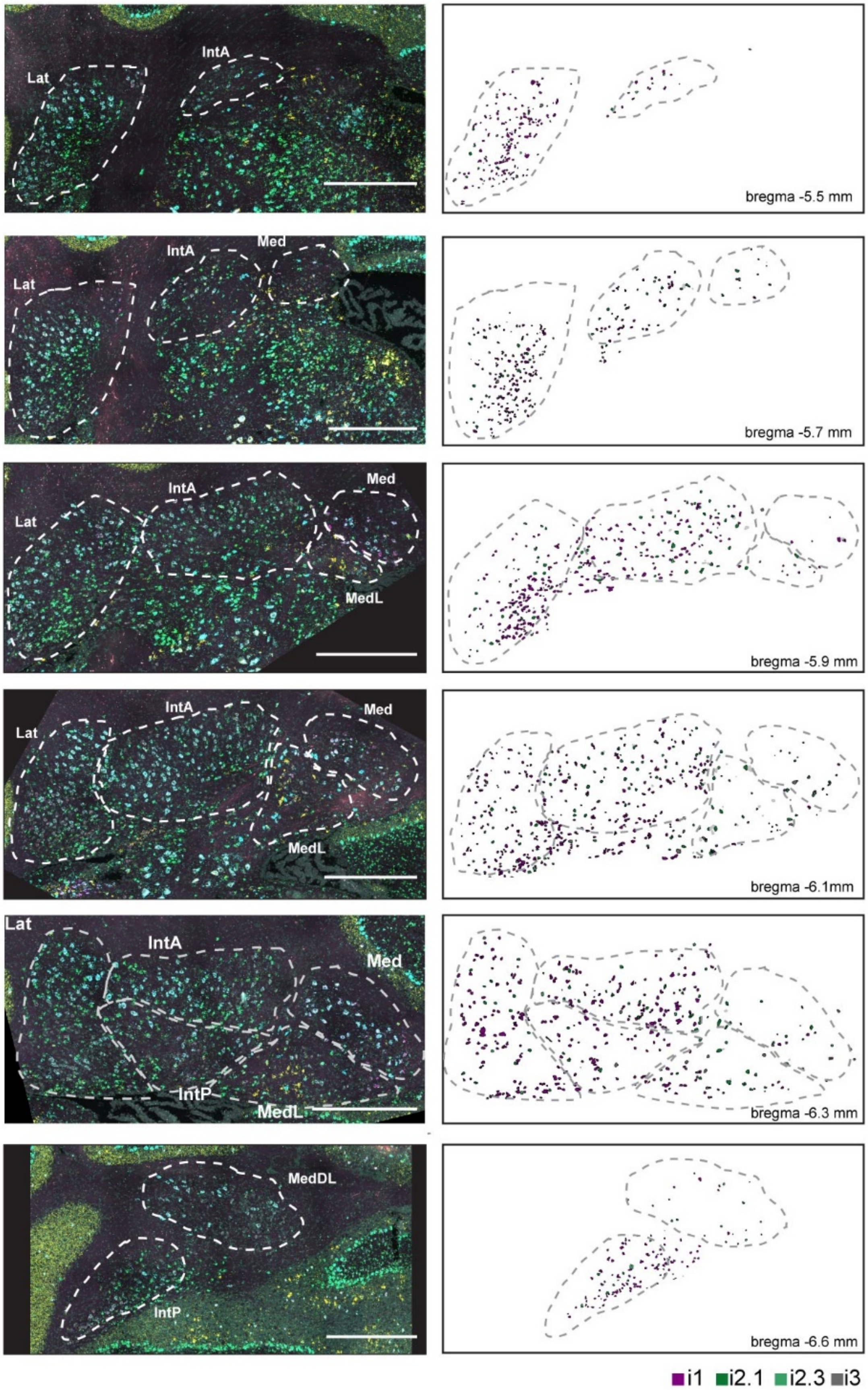


**Figure S18: Spatial location of CN inhibitory cell types.** Representative coronal sections of the CN moving from rostral to caudal illustrating the distribution of inhibitory cells. Inhibitory cells are segmented based on Nissl images and are assigned to a cell type based on gene expression measured by sequential STARmap readout of 12 core marker genes per section. Cell type assignment is based on criteria derived from snRNAseq. Inhibitory cells are colored by cell type. Unassigned inhibitory cells are shown in light grey. We did not attempt to resolve the rare i2.2 cell type. N = 2 mice. Scale bars = 500 µm.

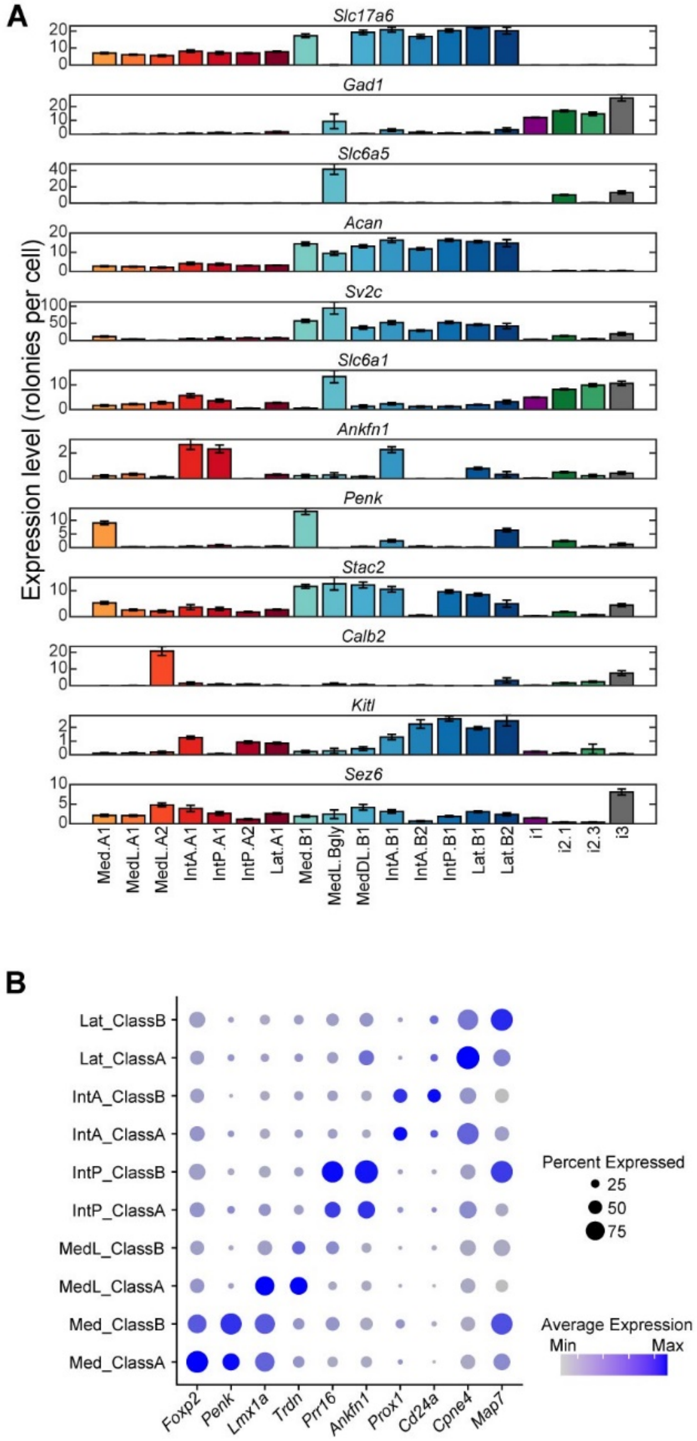


**Figure S19: Cluster-wise gene expression measured by STARmap.** (A) Detected gene expression across cell types from a representative STARmap experiment. Mean ± SEM shown for one representative experiment containing 6 sections spanning the CN. (B) Dot plot showing CN subnucleus marker gene expression derived from snRNAseq data. For each subnucleus, expression of each marker gene in Class A and Class B neurons is indicated. MedDL was merged with MedL for this analysis. There are two axes of variation in gene expression of excitatory CN neurons. One is the difference between Class A and Class B cells (Figs. S14, S15). The other is the difference between subnuclei that affect Class A and Class B cells equally, as shown here.

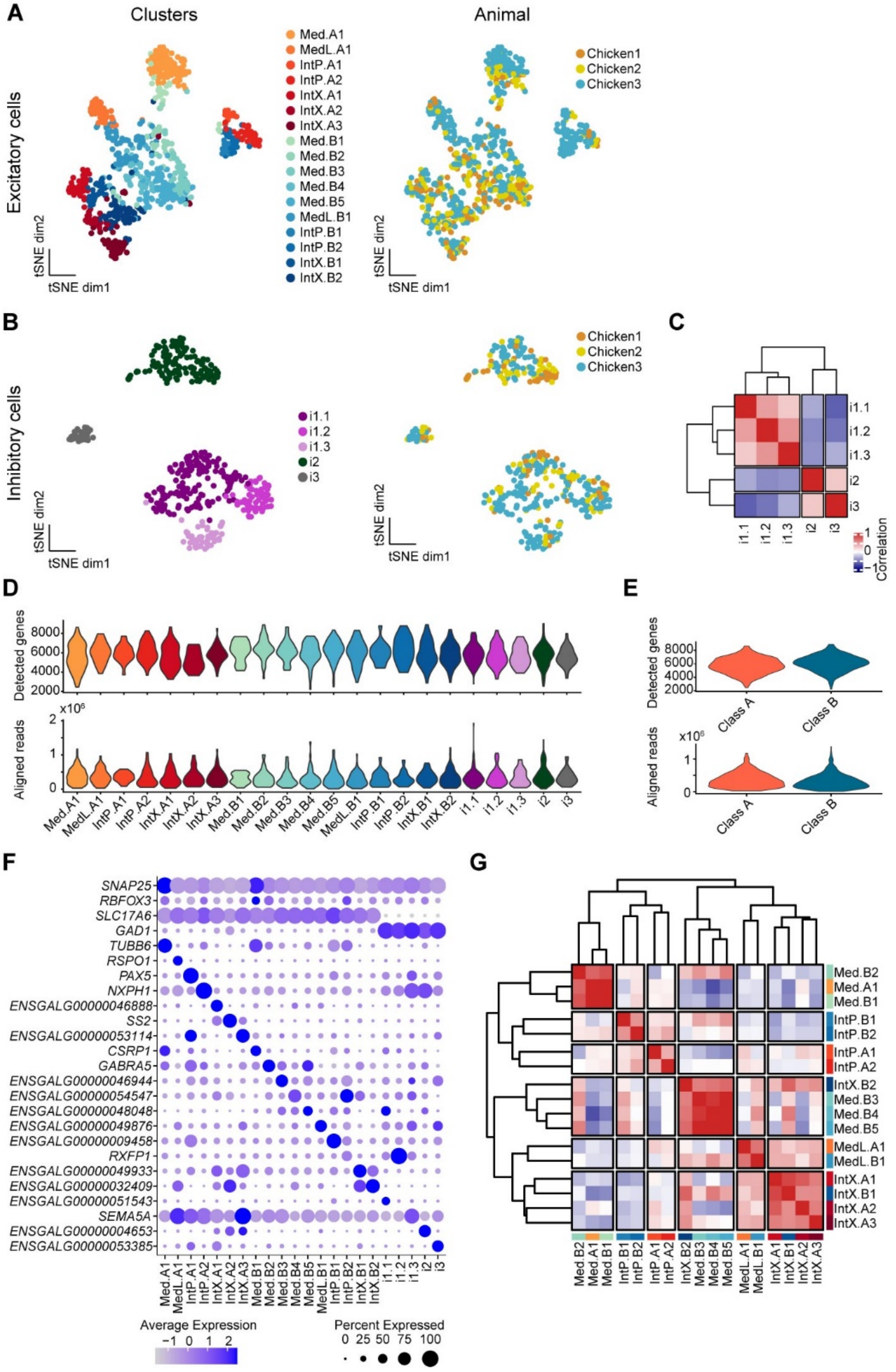


**Figure S20: Details on chicken CN sequencing.** (A, B) Clustering of excitatory (A) and inhibitory (B) CN neurons is independent of the donor animal. (C) Correlation matrix of inhibitory cell types shows that chicken inhibitory cells fall into three classes, with i1.1, i1.2, and i1.3 more similar to each other than to i2 or i3. (D, E) The number of detected genes and aligned reads per cell broken up by cell type (D) or class (E). (F) Marker genes for detected CN cell types. (G) Correlation matrix of all chicken excitatory cell types. In contrast to the mouse excitatory cell types, chicken excitatory cell types group first by brain region followed by class.

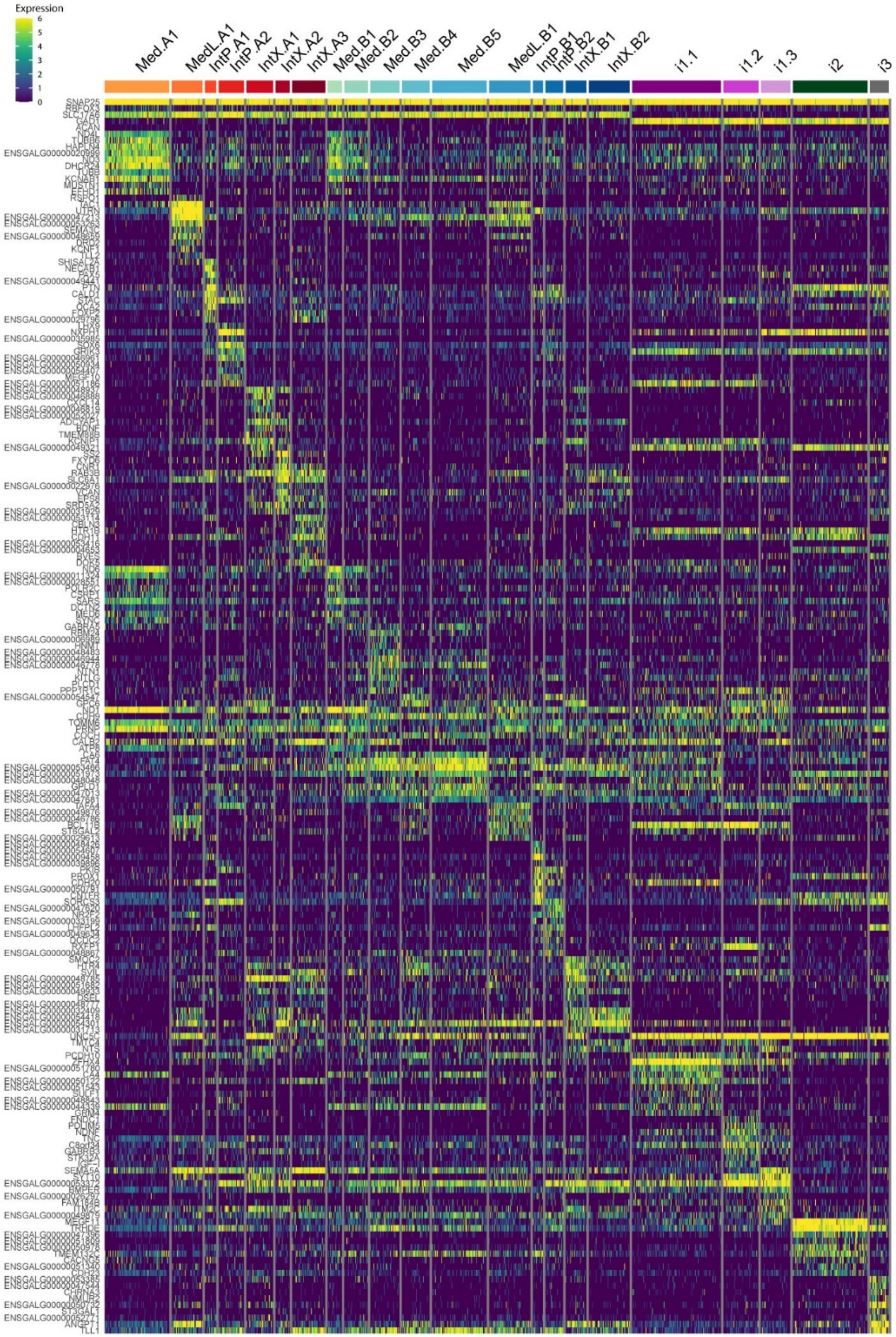


**Figure S21: Differentially expressed genes between chicken CN cell types.**

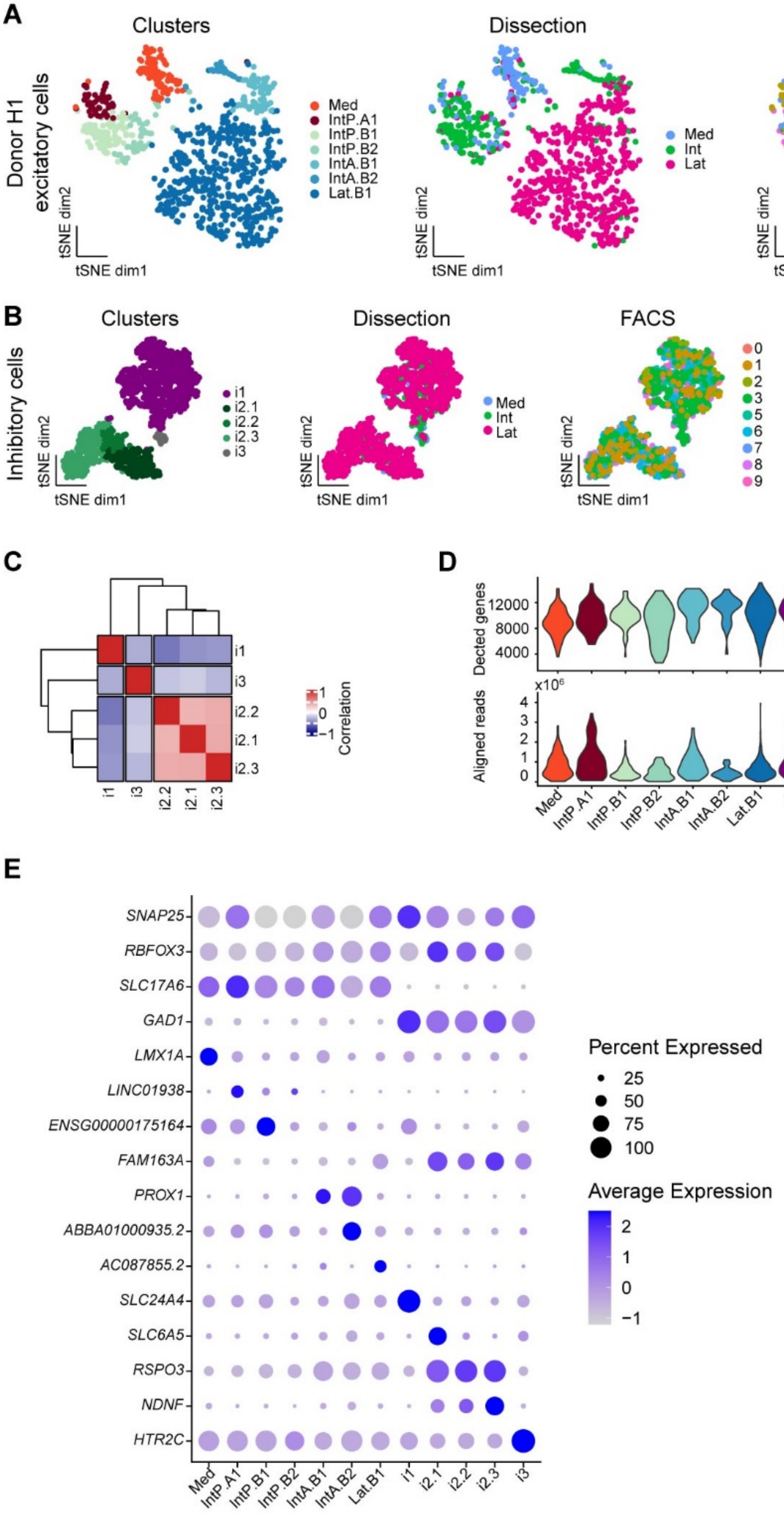


**Figure S22: Details on human CN sequencing.** (A, B) Clustering of donor H1 excitatory cells (A) or all inhibitory cells (B) is independent of FACS round and donor. Note FACS round 9 included only H1

Interposed CN neurons. We focused the analysis of excitatory cells on donor H1 as it contained data for both Interposed and Lateral CN. Donor H2 and H3 only yielded cells from the Lateral CN. (C) Correlation matrix of inhibitory cell types shows that human inhibitory cells fall into three classes, with i2.1, i2.2, i2.3 more similar to each other than to i1 or i3. (D) The number of detected genes and aligned reads per cell broken up by cluster. (E) Marker genes for detected CN cell types.

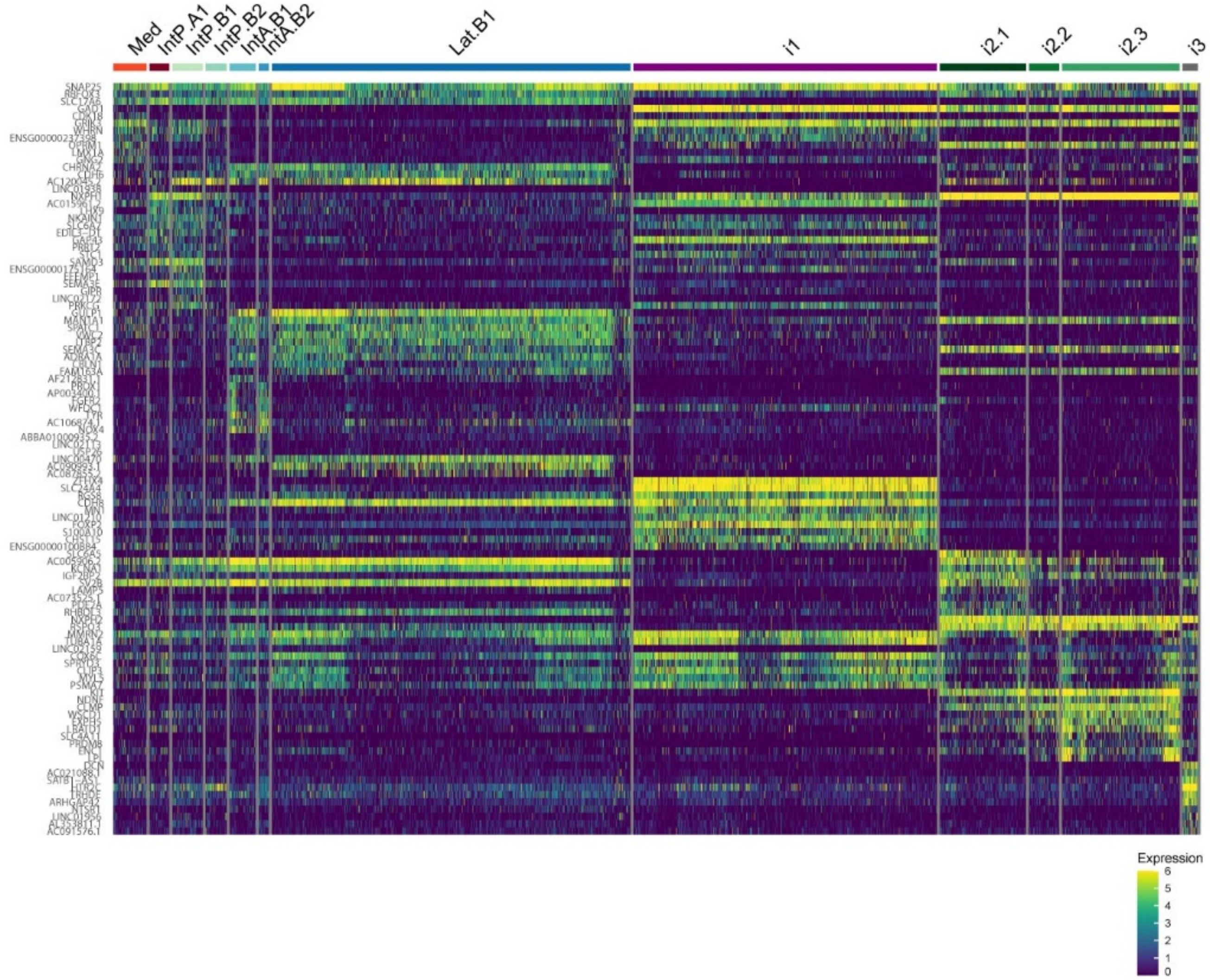


**Figure S23: Differentially expressed genes for human CN cell types.**

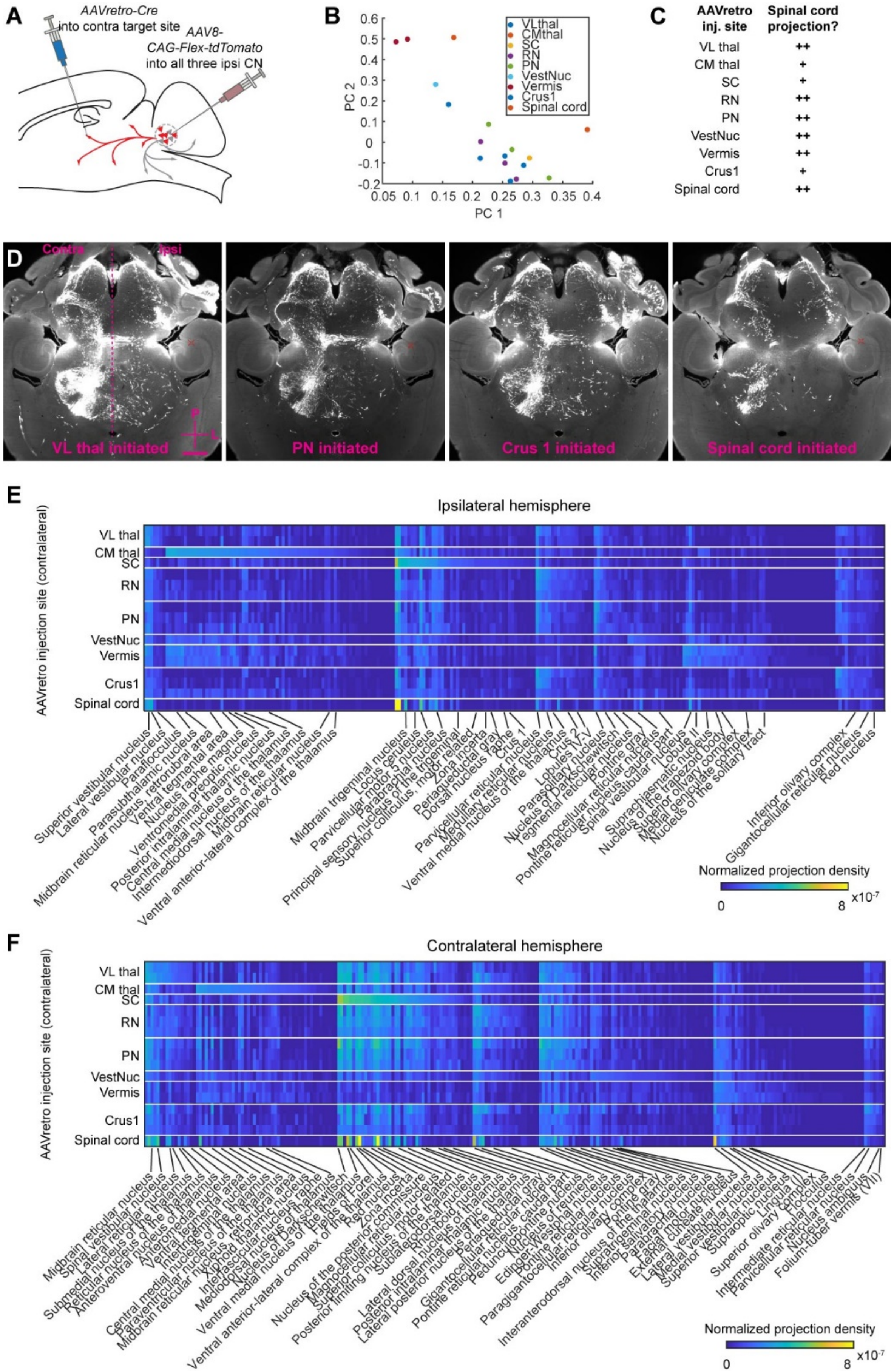
A
AAVretro-Cre
into contra target site
AAV8-
CAG-Flex-tdTomato
into all three ipsi CN
B
VLthal
CMthal
SC
RN
PN
VestNuc
Vermis
Crus1
Spinal cord
PC 1
PC 2
C
AAVretro inj. site
Spinal cord projection?
VL thal ++
CM thal +
SC +
RN ++
PN ++
VestNuc ++
Vermis ++
Crus1 +
Spinal cord ++
D
Contra
Ipsi
VL thal initiated
PN initiated
Crus 1 initiated
Spinal cord initiated
E
Ipsilateral hemisphere
AAVretro injection site (contralateral)
Normalized projection density
x10^-7
F
Contralateral hemisphere
AAVretro injection site (contralateral)
Normalized projection density
x10^-7

**Figure S24: CN neurons collateralize broadly.** (A) Schematic of the experimental workflow. We injected *AAV8-CAG-FLEX-tdTomato* into the ipsilateral mouse CN, covering all three nuclei. We then injected *AAVretro-Ef1a-Cre* into one of several regions contralateral to the CN injections, to label the CNS-wide projection patterns of CN neurons projecting to a given region. After expression, we cleared and imaged brains and spinal cords as before and quantified the CNS-wide innervation projections. (B) Principal component analysis of the brain-wide projection patterns of all processed brains. Different *Cre* injection sites are hard to differentiate in principal component space, indicating overall similarities between the collateralization patterns. (C) Qualitative assessment of spinal cord projections for every *Cre* injection site. (D) Representative horizontal section through brains injected with Cre in the ventrolateral thalamus, pontine nuclei, crus 1 of the cerebellar cortex, or the cervical spinal cord. Raw fluorescence data is shown. Broad and overall strikingly similar projection patterns are apparent. Scale bar = 1 mm. (E, F) Normalized projection densities to all brain regions ipsi- (E) and contralateral (F) to the CN injection sorted by their mean projections per *Cre* injection site. Many regions receive similar input irrespective of where *Cre* was injected. Data is shown for *Cre* injections it ventrolateral (VL) thalamus (N = 2), central medial (CM) thalamus (N = 1), red nucleus (RN; N = 3), pontine nuclei (PN; N = 3), vestibular nuclei (VestNuc; N = 1), vermis (N = 2), crus 1 of the cerebellar cortex (N = 3), and cervical spinal cord (N = 1). Brain regions of interest are indicated.

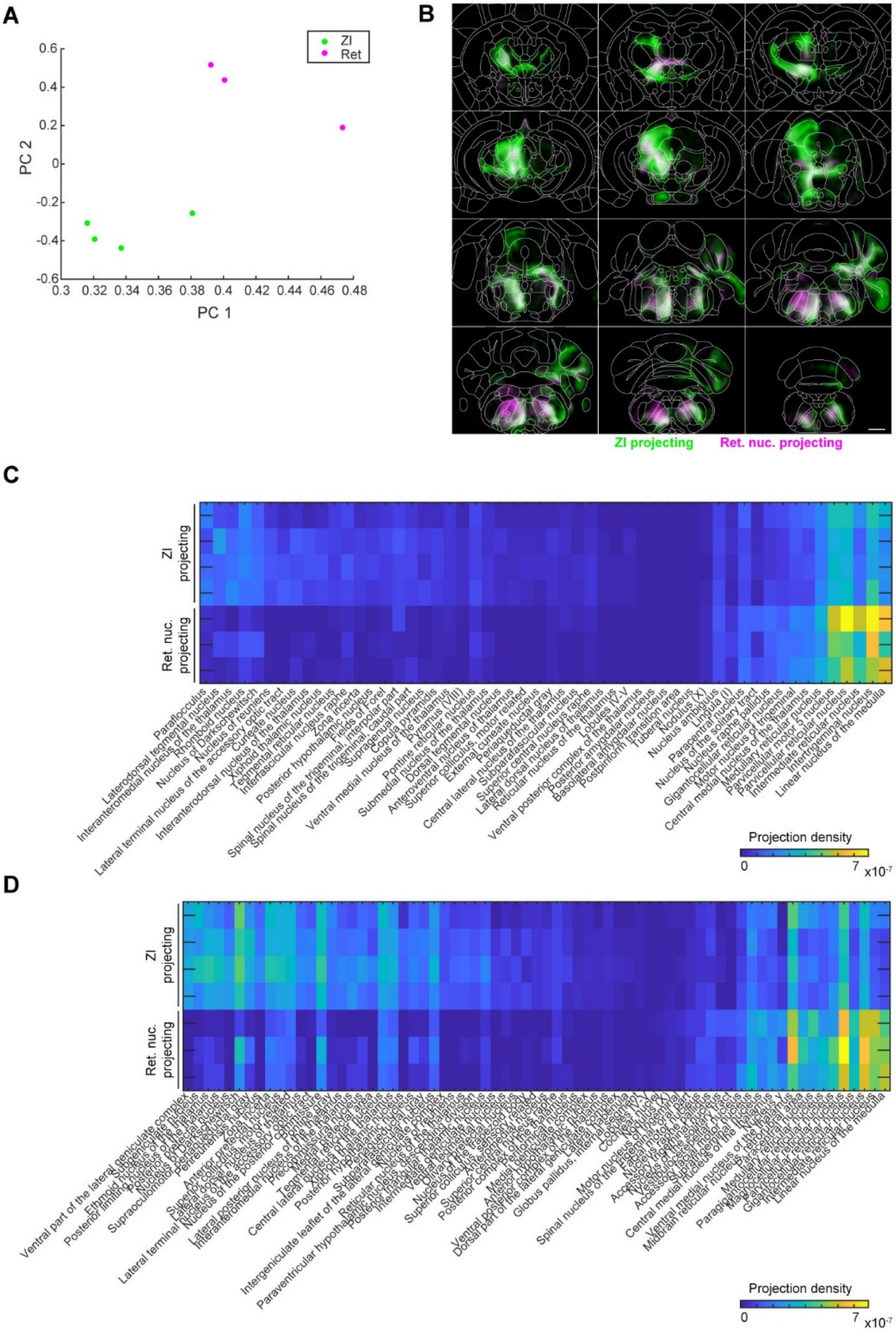


**Figure S25: Differential projections of ZI- and Ret-projecting Lateral CN neurons.** (A) Principal component analysis of 4 brains with ZI injections and 3 brains with Ret injections based on their of the brain-wide normalized innervation densities. The two kinds of brains are easily separated with the first two PCs. (B) Projection probability maps for ZI- and Ret-projecting Lateral CN neurons. Coronal sections separated by 625 μm are shown. Scale bar = 1 mm. (C) Ipsi- and (D) contralateral differentially innervated brain regions ($p < 0.05$, two-sample t-test) sorted by their mean difference in innervation strength.

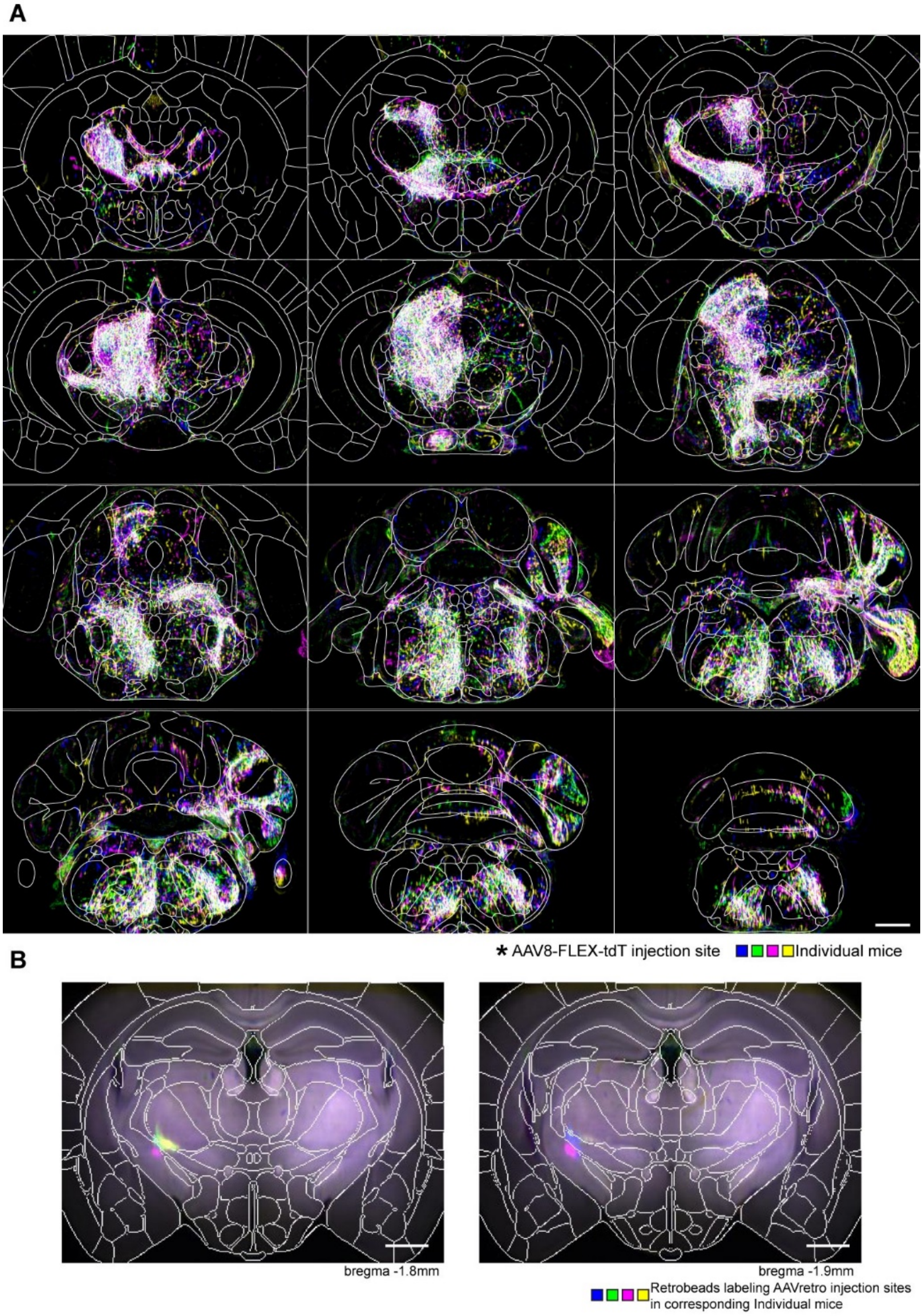


**Figure S26: Brain-wide collateralization patterns of ZI-projecting Lateral CN neurons.** (A) Overlay of detected axons for N = 4 independent injections. Coronal sections are shown, spaced 625 μm apart along the anterior–posterior (A–P) axis, with Allen compartments overlaid in white. Scale bar = 1 mm. (B) Coronal sections at the indicated A–P positions showing the center of the *AAVretro-Ef1a-cre* injection sites for each animal as marked by co-injected retrobeads. Scale bar = 1 mm. Color of axons and retrobeads indicate the mouse of origin.

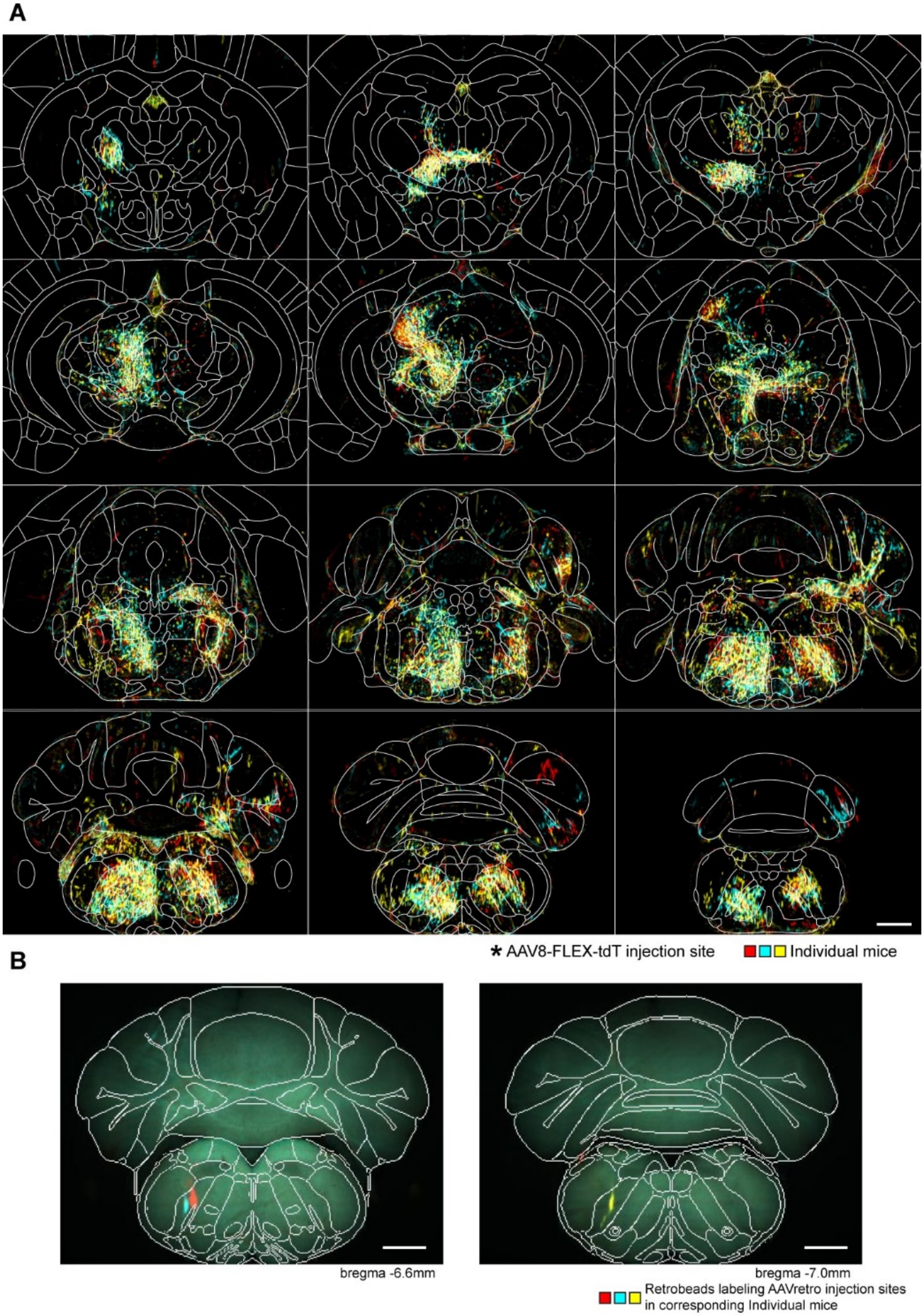


**Figure S27: Brain-wide collateralization patterns of Ret projecting Lateral CN neurons.** N = 3 brains, see Fig. S26 for caption.

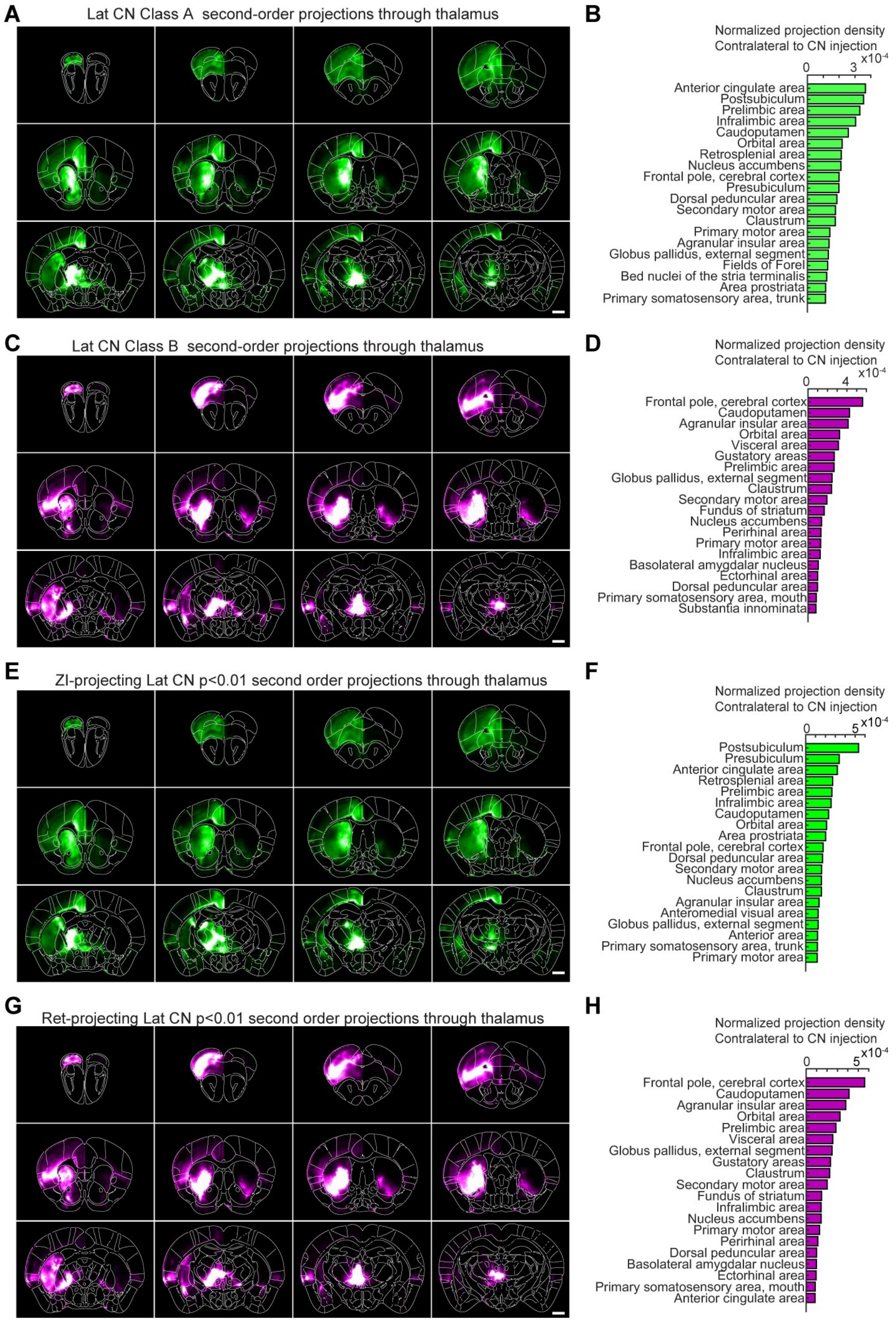


**Figure S28: Differential Lateral CN Class A/B→thalamus→forebrain projections revealed by *in***

***silico* tracing of second-order projections.** (A) Projection probability maps and (B) quantification of contralateral innervation densities from *in silico* tracing of thalamic voxels preferentially innervated by Class A neurons of the Lateral CN. (C, D) Projection probability maps and quantification of contralateral innervation densities from *in silico* tracing of thalamic voxels preferentially innervated by Class B neurons of the Lateral CN. (E–H) Projection probability maps and quantification of *in silico* tracing of second-order projections from thalamic voxels more innervated by ZI-projecting cells than Ret-projecting cells (E, F) and more innervated by Ret-projecting cells than ZI-projecting cells (G, H); starting voxels for *in silico* tracing had $p < 0.01$ without multiple comparison correction using a voxel-wise t-test between ZI and Ret projection maps. Scale bars = 1 mm.

### Supplemental File Captions

**Table S2:** Spreadsheet containing the sequences for all STARmap oligos used in this study.
**Table S3:** Spreadsheet containing the brain-wide normalized projection densities for every anterograde tracing experiment in this study.
**Table S4:** Spreadsheet containing the brain-wide normalized projection densities for the ZI/Ret collateralization mapping experiments.

**Movie S1: Fly-through of aligned raw axonal projections of the anterior Medial CN.** Axons from each mouse are shown in a distinct color. N=5.
**Movie S2: Fly-through of aligned raw axonal projections of the posterior Medial CN.** Axons from each mouse are shown in a distinct color. N=5.
**Movie S3: Fly-through of aligned raw axonal projections of the Interposed CN.** Axons from each mouse are shown in a distinct color. N=6.
**Movie S4: Fly-through of aligned raw axonal projections of the Lateral CN.** Axons from each mouse are shown in a distinct color. N=7.
**Movie S5: Fly-through of aligned raw axonal projections of ZI-projecting neurons from the Lateral CN.** Axons from each mouse are shown in a distinct color. N=4.
**Movie S6: Fly-through of aligned raw axonal projections of Ret-projecting neurons from the Lateral CN.** Axons from each mouse are shown in a distinct color. N=3.
**Movie S7: Fly-through of second-order projection from thalamic voxels that were primarily innervated by Class A (green) or Class B (magenta) Lateral CN neurons.**
